# Effects of 4 testing arena sizes and 11 types of embryo media on sensorimotor behaviors in wild-type and *chd7* mutant zebrafish larvae

**DOI:** 10.1101/2023.07.31.551330

**Authors:** Dana R. Hodorovich, Tiara Fryer Harris, Derek Burton, Katie Neese, Rachael Bieler, Vimal Chudasama, Kurt. C Marsden

## Abstract

The larval zebrafish is a highly versatile model across research disciplines, and the expanding use of behavioral analysis has contributed to many advances in neuro-psychiatric, developmental, and toxicological studies, often through large-scale chemical and genetic screens. In the absence of standardized approaches to larval zebrafish behavior analysis, however, it is critical to understand the impact on behavior of experimental variables such as the size of testing arenas and the choice of embryo medium. Using a custom-built, modular high-throughput testing system, we examined the effects of 4 testing arena sizes and 11 types of embryo media on conserved sensorimotor behaviors in zebrafish larvae. Our data show that testing arena size impacts acoustic startle sensitivity and kinematics as well as spontaneous locomotion and thigmotaxis, with fish tested in larger arenas displaying reduced startle sensitivity and increased locomotion. We also find that embryo media can dramatically affect startle sensitivity, kinematics, habituation, and pre-pulse inhibition, as well as spontaneous swimming, turning, and overall activity. Common media components such as methylene blue and high calcium concentration consistently reduced startle sensitivity and locomotion. To further address how the choice of embryo medium can impact phenotype expression in zebrafish models of disease, we reared *chd7* mutant larvae, a model of CHARGE syndrome with previously characterized morphological and behavioral phenotypes, in 5 different types of media and observed impacts on all phenotypes. By defining the effects of these key extrinsic factors on larval zebrafish behavior, these data can help researchers select the most appropriate conditions for their specific research questions, particularly for genetic and chemical screens.

## Introduction

The teleost *Danio rerio* or zebrafish has become a powerful model in biological and biomedical research. The larval zebrafish is an accessible and versatile model across multiple research fields including neuroscience, genetics, developmental biology, toxicology, immunology, and more^1–7^. Advantages of the zebrafish model include external fertilization and development, transparent embryos and larvae, high fecundity, cost-effective maintenance^8^, and defined orthologs for ∼70% of human genes^9^. As use of the larval zebrafish model continues to increase, particularly with high-throughput approaches using larval zebrafish behavior as a readout of neural function, it has become more important to define how key experimental variables affect larval development and behavior.

By 5 days post fertilization (dpf), larvae reliably perform conserved auditory- and visually-mediated behaviors^10–13^, making them a powerful tool for multiple research questions in developmental neuroscience, toxicology, and chemical or genetic screening^1, 4, 14, 15^. For example, high-throughput screening of visual and locomotor behavior of larvae treated with various compounds is an established measure of toxicity^6, 14^. However, while standardized protocols for comparable experiments with rodents have been well-established^16^, protocols for raising and testing zebrafish larvae can vary widely, and experimental variables such as testing arena size and embryo medium have been reported to impact larval zebrafish behavior^17, 18^. The influence of media on phenotypes is especially critical when screening mutant lines, where media may have a genotype-specific effect that could mask potential differences. Therefore, to ensure the reliability and reproducibility of findings from labs around the world, it is necessary to define the effects that testing arena size and embryo media type have on a range of larval zebrafish behaviors.

In this study, we assayed multiple behaviors including the acoustic startle response, short-term habituation, pre-pulse inhibition, spontaneous locomotion, as well as kinematic performance in larvae tested in 4 arena sizes and raised in 11 different media. Additionally, we tested an established zebrafish model of CHARGE Syndrome^19^ (*chd7^ncu^*^101^*^/^*) with defined morphological and behavioral phenotypes in different media to assess the impact on phenotype expression. Our results show arena- and media-dependent differences affecting multiple sensorimotor behaviors. Increasing testing arena diameter significantly decreased startle frequency and increased thigmotaxis. Additionally, media with additives including methylene blue, higher calcium concentrations, and sodium bicarbonate exerted significant and bidirectional changes in both conserved auditory and general locomotor behaviors compared to control embryo media (1x E3). Finally, homozygous *chd7* mutants (*chd7^ncu^*^101^*^/ncu^*^101^) displayed the highest penetrance of morphological phenotypes and significant changes in auditory-driven behaviors when reared in media containing methylene blue. We observed high degrees of variation within the same genotype across media types, indicating a genotype-specific influence on auditory and locomotor behaviors. Together these results demonstrate how extrinsic experimental factors impact larval zebrafish behavior as well as phenotypic penetrance in a zebrafish disease model.

## Materials and methods

### Zebrafish Husbandry and Maintenance

All animal use and procedures approved by the North Carolina State University Institutional Animal Care and Use Committee. All testing was performed with the *Tüpfel long fin* (TLF) strain. The wild-type TLF line originated from University of Pennsylvania stocks, and *chd7^ncu^*^101^*^/+^* line was maintained in the TLF strain which originated from Zebrafish International Resource Center (ZIRC) stocks. Adult zebrafish were housed in 5 fish/L density under a 14 hr:10hr light:dark cycle at ∼28°C, and were fed rotifers, *artemia* brine shrimp (Brine Shrimp Direct), and GEMMA micro 300 (Skretting).

To generate embryos for larval testing, male and female pairs were placed in mating boxes (Aquaneering) containing system water and artificial grass. 1-2 hours into the subsequent light cycle of the following day, embryos were collected and placed into petri dishes containing 1x E3 media with 0.05% methylene blue. Embryos were sorted for fertilization under a dissecting scope at ∼6 hours post fertilization (hpf) and placed into 10 cm petri dishes with n ≤ 60. For media experiments, 1x E3 media was completely removed and embryos were placed in experimental media following sorting. All embryos were reared in a temperature-controlled incubator at 29°C on a 14h:10h light:dark cycle. Each day until testing, a 50-75% media change was performed. All experiments were performed blind to genotype, when applicable.

### Embryo Media

For media experiments, 11 commonly used laboratory media were selected with 1x E3 as the control media. The 11 solutions included: 1x E3, 1x E3 with 0.05% methylene blue (1x E3 MB), 1x E2, 1x E2 with 0.05% methylene blue (1x E2 MB), 0.5x E2 with 0.05% methylene blue (0.5x E2 MB), Normal Ringer’s solution (Ringer’s), High calcium Ringer’s (Hi Ca^2+^ Ringer’s), Bath solution, 10% Hank’s, Egg water, and system water from our laboratory’s fish facility. Details about each medium, including solute concentrations, are reported in Supplemental Table 1.

### Behavioral Assays and Analyses

Larvae were tested at 5-6 dpf within their normal light cycle. Larvae were screened for morphological defects including uninflated swim bladder, pericardial edema, and bent tail. For media and *chd7^ncu^*^101^*^/^* experiments, swim bladder inflation and less severe morphological phenotypes (those not affecting mobility) were ignored. Post-screening, larvae adapted to the testing lighting and temperature conditions for 30 minutes before placement in a custom laser-cut acrylic grid consisting of 36 circular wells each with 9 mm diameter and 2 mm depth. For arena size experiments, 4 well diameters were tested: 9 mm (36 wells), 13 mm (16 wells), 18 mm (9 wells), and 28 mm (4 wells). All wells had 2 mm depth. The grid was anchored to a 60×5×5 mm aluminum rod attached to an acoustic shaker (Bruel and Kjaer). Spontaneous movement, acoustic startle, short-term habituation, and pre-pulse inhibition assays were tracked and analyzed using FLOTE-software as previously described^11, 12, 20^.

### Spontaneous Movement and Thigmotaxis

Once individually placed in 9 mm round wells for all media experiments, or 9-, 13-, 18-, and 28-mm wells for testing arena experiments, larvae acclimated for 3 minutes and were then recorded for 18.5 minutes at 50 fps at 640×640 px resolution with a Photron mini-UX50 high-speed camera. Thigmotaxis was analyzed by dividing the area of each well into its center and periphery, with each accounting for 50% of the total area and calculating the distance from center throughout the time course (√[(□ - 50)^2^ + (□ - 50)^2^]), where □ and □ represent the coordinates of the larvae relative to the center of the arena. If the calculated distance from center was less than 25, or half of the center area, larvae were classified as in the center. If the distance from center was greater than 25, larvae were classified as in the perimeter.

### Acoustic Startle Response and Auditory behaviors

Larvae acclimated for 2 minutes prior to auditory stimulus onset. Larvae received a total of 60 acoustic stimuli: 10 pseudorandomized trials of 6 intensities (13.6, 25.7, 29.2, 35.5, 39.6, and 53.6 dB), with a 20 s inter-stimulus interval (ISI). Immediately following the acoustic startle response assay, larvae received 10 trials of a pre-pulse stimulus at 29.2 dB and a pulse of 53.6 dB 300 ms later, with each trial separated by a 20s ISI to assess pre-pulse inhibition. At the end of the assay, larvae received 30 additional acoustic stimuli (53.6 dB) with a 1s ISI to induce short-term habituation. Recordings were captured at 1000 fps and 640×640 px resolution with a Photron mini-UX50 high-speed camera.

### Morphological phenotype assessment of *chd7^ncu^*^101^*^/^*

Larvae were assessed for morphological phenotypes after behavior testing. Larvae were anesthetized in 0.2% Tricaine (MS-222) and manually assessed by two independent experimenters, blinded to genotype. Morphological phenotypes were determined as previously described^19^.

### DNA extraction and Genotyping *chd7^ncu^*^101^*^/^*

Larvae were individually placed in 96-well plates containing methanol (MeOH) for tissue fixation. DNA was extracted from whole larvae using a lysis buffer of 25mM sodium hydroxide, 0.2 mM EDTA (base solution), and 40mM Tris-HCl (neutralization solution). PCR and gel electrophoresis was performed as previously described^19^.

### Statistics

Statistical analyses were completed using Prism 8 (GraphPad) and JMP Pro 14 and JMP Pro 16 (SAS). All datasets were tested for normality using the Shapiro-Wilk test. One-way ANOVA compared to controls and multiple comparisons or nonparametric tests (Wilcoxon each pair) were used, respectively. For media experiments, p-value threshold was set α=0.01, and testing arena and *chd7^ncu^*^101^*^/^* experiments was α=0.05. All data is represented as mean ± standard deviation, unless otherwise noted in the figure legends.

## Results

### Larger testing arenas cause acoustic hypo-responsiveness in zebrafish larvae

Some neuropsychiatric and congenital disorders such as schizophrenia and CHARGE syndrome commonly present with auditory-driven behavioral symptoms^21, 22^. Symptoms are typically in the form of dysregulated sensorimotor gating, and conductive/sensorineural hearing loss, respectively^21–23^. Larval zebrafish are a robust model to study clinically relevant auditory-derived symptoms because by 5 days post fertilization (dpf) larvae can perform multiple auditory-driven behaviors including two types acoustic startle response (short- and long-latency C-bends; SLC and LLCs), pre-pulse inhibition, and short-term habituation^11, 24, 25^. To determine the effect that testing arena diameter has on the acoustic startle response and its kinematics, we assayed 6 dpf wild-type TLF-strain larvae that were reared together in 1x E3 medium in 10 cm petri dishes and then tested individually in one of four testing arena sizes: 9 mm, 13 mm, 18 mm, and 28 mm diameter circular wells. All arenas had 2 mm depth. Larvae received 60 acoustic stimuli: 10 trials at each of 6 intensities, with a 20 second interstimulus interval (ISI). Following an acoustic stimulus, zebrafish larvae perform one of two distinct response types: 1) Short-latency C-bends (SLCs) and 2) Long-latency c-bends^11^. SLCs and LLCs are defined by their response latency and kinematics and are mediated by separate neural circuits^11, 26^. As decibel value increases, larvae bias their responses towards SLCs, whereas LLCs are more likely to be performed following lower intensity stimuli^11^ (Fig. 1A). To assess individual response rates of SLCs and LLCs, we calculated sensitivity indices by taking the area under the curve (AUC) for individual frequency curves (Fig. 1B,C). Compared to the 9 mm arena, larvae tested in all three larger arenas (13, 18, and 28 mm) showed a reduction in SLC frequency as decibel value increased (Fig. 1A,B), but LLCs were unaffected (Fig. 1C,D). Statistical analyses for multiple comparisons are reported in Supplemental Table 2. The SLC response is a stereotyped series of movements following an intense acoustic stimulus including an initial C-bend (C1), counter-bend (C2), and swim bout^10–12^. Because of the highly stereotyped nature of the SLC response movements, we compared specific kinematic parameters of SLCs performed by larvae in different arena sizes, including latency of response initiation, bend angles (C1 and C2), C1 bend curvature, distance traveled, and C1 angular velocity (Fig. 2). The initial C-bend (C1) can be quantified using the angle, or the changes in head orientation, and curvature, or the summed angle calculated from the orientation change of the head/body and body/tail positions of the larvae^12^. Although we detected differences in SLC responsiveness, SLC kinematics were mostly unchanged including latency, C1 curvature, and SLC distance (Fig. 2A,C,D). We observed significant differences for C1 angle in the 13 mm arena (Fig. 2B), and maximum angular velocity in the 13 mm and 18 mm arenas (Fig. 2E). Larvae in the three largest arena sizes displayed significant increases in C2 bend angle (Fig. 2F). This may be at least partly due to the fact that in a larger testing arena larvae are less likely to contact the wall of the arena following the initial C1-bend.

**Figure 1.**
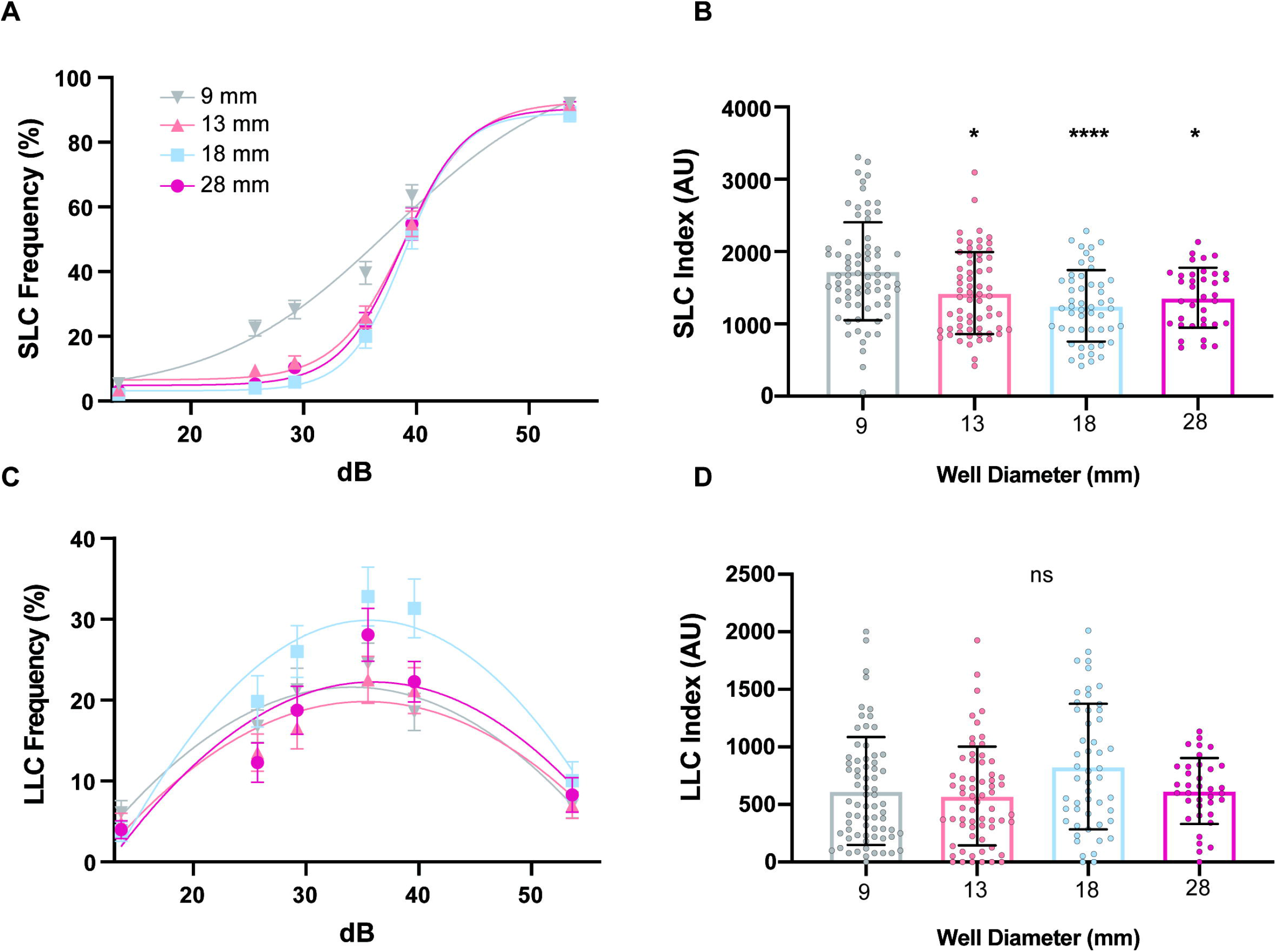
Testing arena size affects short-latency c-bends. **(A)** Acoustic startle responses, average short-latency c-bend (SLC) frequency comparing testing arena sizes (9 mm n=72, 13 mm n=64, 18 mm n=51, 28 mm n=36) (mean ± SEM). **(B)** Short-latency c-bend sensitivity index, calculated by the area under the SLC frequency curve for individual larvae (mean ± SD). **(C)** Average long-latency c-bend (LLC) frequency comparing testing arena sizes (mean ± SEM). **(D)** Long-latency c-bend sensitivity index, calculated by the area under the LLC frequency curve. Asterisks represent statistical significance for arena sizes compared to 9 mm arena (mean ± SD, Wilcoxon/ Kruskal-Wallis tests with Wilcoxon each pair for nonparametric multiple comparisons, *p<0.05, **p<0.01, ****p<0.0001).

**Figure 2.**
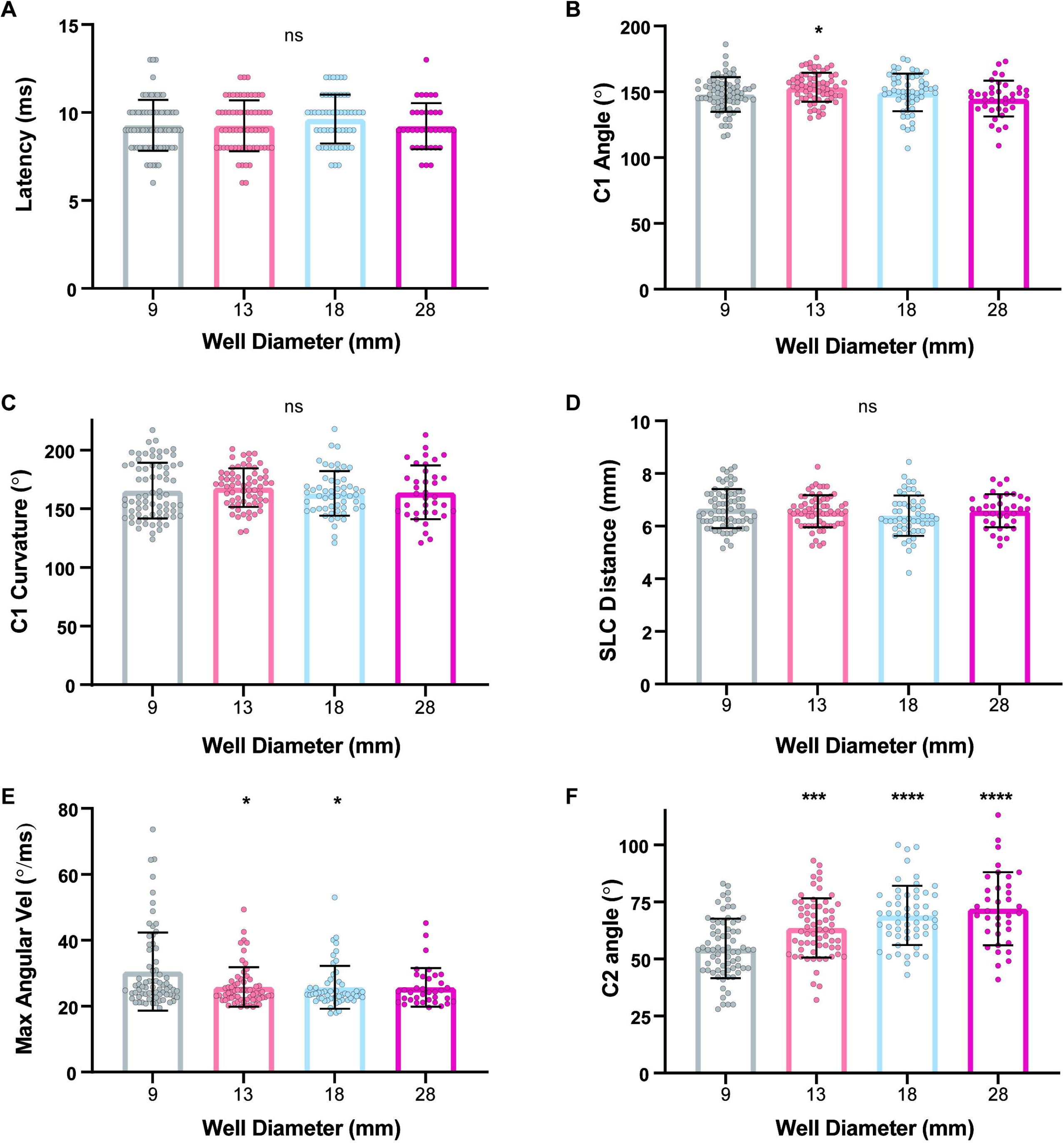
SLC kinematics compared across testing arena sizes. **(A)** SLC response kinematics of individual larvae including latency of response initiation **(B)** C1 bend angle, **(C)** C1 bend curvature, **(D)** average distance traveled during response, **(E)** maximum angular velocity of C1 bend, and **(F)** C2 bend angle (9 mm n=71, 13 mm n=64, 18 mm n=54, 28 mm n=36). Asterisks represent statistical significance for arena sizes compared to 9 mm arena (mean ± SD, One-way ANOVA with student’s t each pair test for multiple comparisons, Wilcoxon/ Kruskal-Wallis tests with Wilcoxon Each Pair for nonparametric multiple comparisons, *p<0.05, ***p<0.001, ****p<0.0001).

### Activity levels and thigmotaxis behavior are dependent on arena size

Previous work has demonstrated differences in overall activity levels of larval zebrafish as arena size changes^17, 27, 28^, but to determine if arena size alters the types of movement performed and/or the location of larvae within the arena, we recorded and measured spontaneous locomotion at 50 frames per second for 18.5 min and analyzed the frequency of swim and turn movements^12, 29^, total distance traveled, and thigmotaxis (preference for proximity to the wall) behavior. We found robust differences in total distance traveled as arena size increased (Fig. 3A,B), compared to the 9 mm arena. Larvae tested in 13 mm and 28 mm arenas displayed increased swim and turn frequencies (Fig. 3C,D), whereas the larvae in the 18 mm arena performed more turns but similar swim frequency (Fig. 3D). In addition to general locomotor behaviors, we analyzed thigmotaxis, defined as the percentage of time spent in the perimeter of the arena and a common measurement used in both rodent and zebrafish testing as an indicator of exploratory behavior and anxiety-like states^30–32^. Center and perimeter were defined such that each accounted for 50% of the total area (Fig. 3E), and thus if the larva had no preference, it would be equally likely to be in the center or perimeter. Larvae in the 28 mm arena exhibited similar thigmotaxis rates compared to the 9 mm arena, while larvae in the 13 mm and 18 mm arenas spent significantly more time in the center (Fig. 3F). While larvae may be expected to spend the vast majority of their time in the perimeter of the smallest 9 mm arena since the well is only ∼2.5 times their body length, the increased thigmotaxis in the largest arena (28 mm) relative to the medium-sized 13- and 18-mm arenas suggests that larvae strongly prefer to remain close to the wall when exposed to a large, open space.

**Figure 3.**
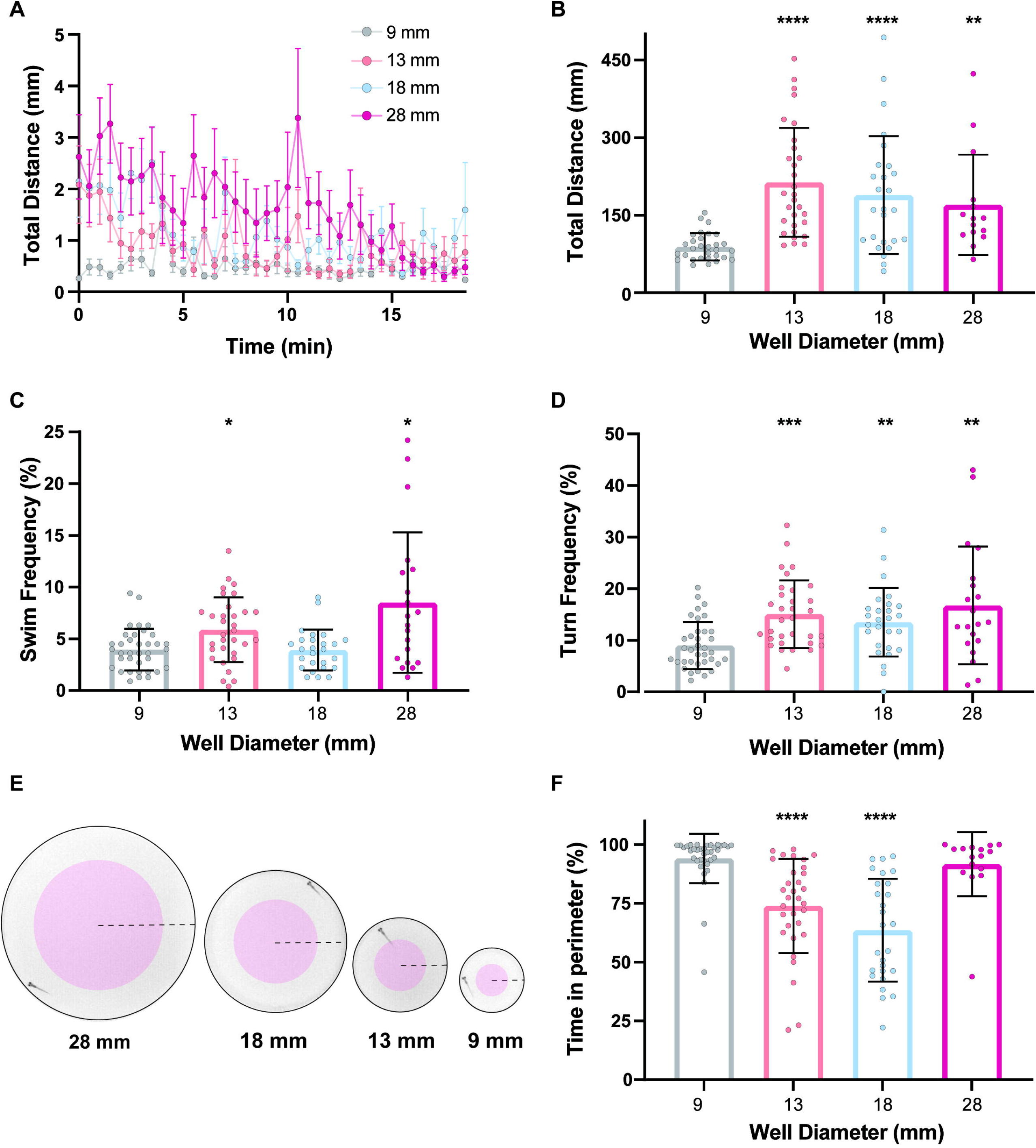
Arena size influences general locomotor activity and thigmotaxis behavior. **(A)** Total distance time plot during 18.5 minutes of recording comparing testing arena sizes (9 mm n=30, 13 mm n=30, 18 mm n=26, 28 mm n=20)(mean ± SEM). **(B)** Sum of distance traveled for individual larvae, and **(C)** swim and **(D)** turn frequencies (mean ± SD). **(E)** Representative diagrams of defined perimeter (white), center (red), and radius (dashed line) of each testing arena size. **(F)** Percent of time spent in the perimeter during the 18.5 min testing period. Asterisks represent statistical significance for arena sizes compared to 9 mm arena (mean ± SD, Wilcoxon/ Kruskal-Wallis tests with Wilcoxon Each Pair for nonparametric multiple comparisons, *p<0.05, **p<0.01, ***p<0.001, ****p<0.0001).

### Embryo medium influences auditory-driven behaviors and kinematics

Besides the testing arena size, embryo medium is another controllable variable that may influence larval behavior. To determine if media type affects auditory-driven behavioral responses, we performed the same acoustic startle assay as above on larvae reared in 11 different types of commonly used media. To account for unequal sample sizes (Table S3), we normalized AUC data to 1x E3. Larvae reared in 1x E3 MB, 0.5 X E2 MB, and Hi Ca^2+^ Ringer’s displayed a significant reduction in SLC responses, while those raised in Hank’s medium showed an increas in SLCs compared to 1x E3 (Fig. 4A,B). 0.5x E2 MB-reared larvae also performed more LLCs compared to 1x E3 (Fig. 4C), indicating a shift in response type bias. Additionally, 1x E2 MB- and Bath solution-reared larvae displayed a significant increase or decrease in LLC responses, respectively. (Fig. 4C). The independent effect of a medium on one of the two startle responses further illustrates the distinction between the two behaviors and suggests that media type may specifically impact the development and/or function of either the SLC or LLC circuit.

**Figure 4.**
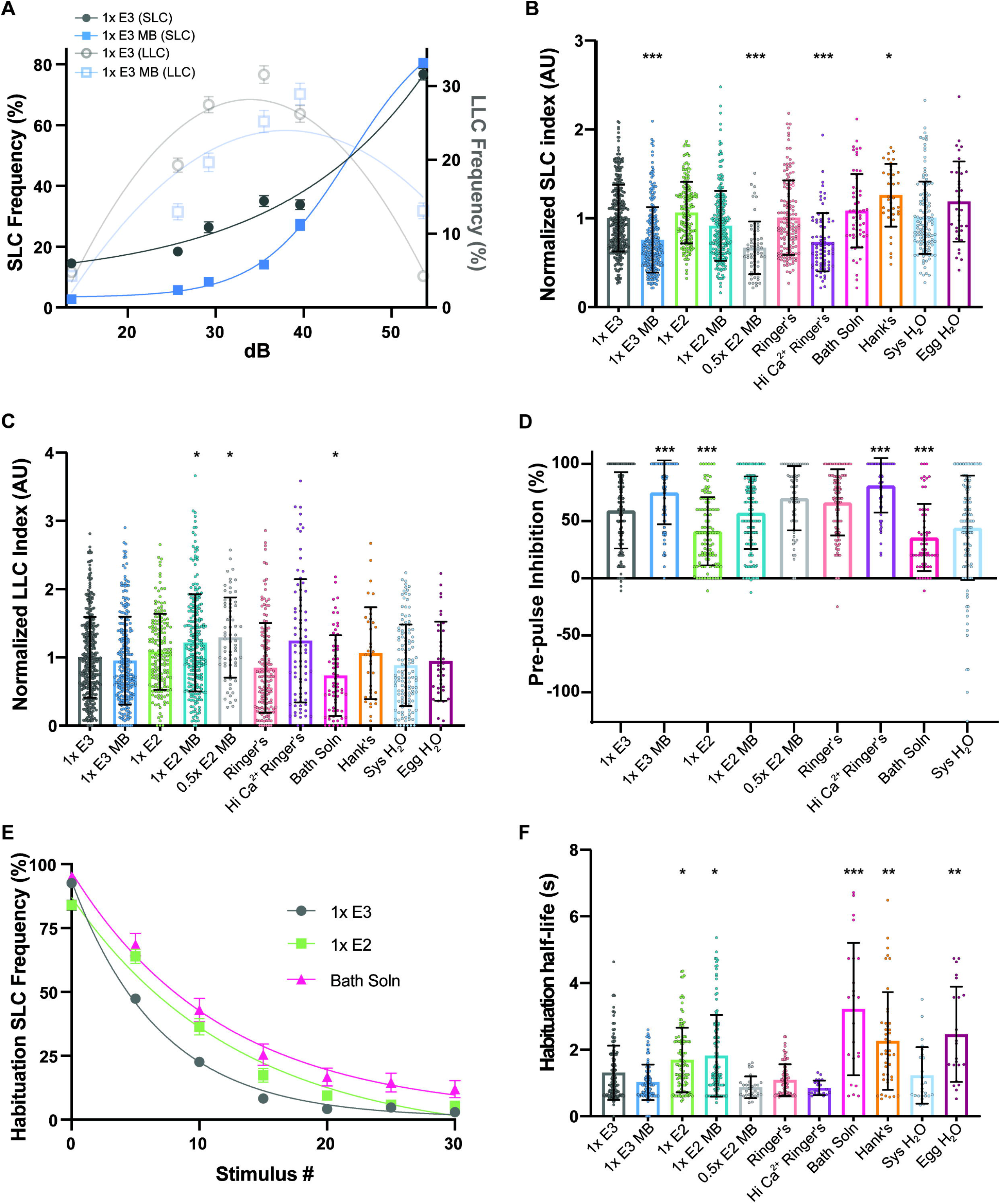
Embryo media type impacts auditory-driven behaviors. **(A)** Acoustic startle responses, average short-latency c-bend (SLC) frequency (left y-axis) and long-latency c-bend (LLC) frequency (right y-axis) as acoustic stimulus intensity increases in 1x E3 or 1x E3 with methylene blue treated larvae (mean ± SEM). **(B)** Short-latency c-bend and **(C)** Long-latency c-bend sensitivity indices, calculated by the area under the SLC or LLC frequency curves, respectively, for individual larvae, normalized to 1x E3 (1x E3 n= 397, 1x E3 MB n=245, 1x E2 n=166, 1x E2 MB n=231, 0.5x E2 MB n=64, Ringer’s n=141, Hi Ca^2+^ Ringer’s n =79, Bath Soln n=61, Hank’s n=36, Sys H_2_O n=123, Egg H_2_O n=36) (mean ± SD). **(D)** Rate of pre-pulse inhibition (1x E3 n= 263, 1x E3 MB n=118, 1x E2 n=108, 1x E2 MB n=137, 0.5x E2 MB n=63, Ringer’s n=115, Hi Ca^2+^ Ringer’s n =62, Bath Soln n=61, Sys H_2_O n=123)(mean ± SD). **(E)** Short-term habituation, average SLC frequency during 30 acoustic stimuli at highest intensity in 1x E3, 1x E2 and Bath Solution treated larvae (mean ± SEM). **(F)** SLC half-life calculated by nonlinear regression (one-phase exponential decay) of SLC frequency curves for individual larvae. Asterisks represent statistical significance for media type compared to 1x E3 (mean ± SD, Wilcoxon/ Kruskal-Wallis tests with Wilcoxon Each Pair for nonparametric multiple comparisons, *p<0.01, **p<0.001, ***p<0.0001).

To further examine the influence media may have on sensorimotor gating, we performed a pre-pulse inhibition (PPI) assay. Larvae received 10 trials in which a weak stimulus (29.2 dB) was immediately followed by an intense stimulus (53.6 dB) 300 ms later. We calculated %PPI with larval SLC response rates following an intense stimulus alone compared to the paired stimuli. Relative to larvae raised in 1x E3, PPI was significantly increased in 1x E3 MB and Hi Ca^2+^ Ringer’s larvae, while 1x E2 and Bath solution reduced PPI (Fig. 4D). Hank’s and Egg H_2_O were not tested for PPI. Given that 1x E3 MB and Hi Ca^2+^ Ringer’s reduced SLC responses, an increase in PPI suggests these conditions suppress activity in the SLC circuit in both contexts.

Finally, we analyzed short-term habituation, a simple form of non-associative learning in which SLC responses rapidly decrease following repeated strong acoustic stimuli^24, 25^. Larvae received 30 strong acoustic stimuli (53.6 dB) separated by a 1s ISI (Fig. 4E). To measure the rate of habituation we calculated the half-life of SLC frequency during the assay and found that larvae reared in 1x E2, 1x E2 MB, Bath soln, Hank’s, and Egg H_2_O displayed slower habituation rates compared to 1x E3. The assay- and context-dependent differences in auditory behaviors suggests that embryo media can influence specific characteristics of these behavioral responses, rather than having a global effect on response rates.

Larvae raised in 1x E3 MB, 0.5x E2 MB, and Hi Ca^2+^ Ringer’s initiated SLCs with significantly longer latencies, while Bath solution, Hank’s, and Egg water-raised larvae responded with shorter latencies compared to 1x E3 (Fig. 5A). We found that larvae reared in Ringer’s, Hi Ca^2+^ Ringer’s, and Bath solution displayed a reduction in both C1 angle and C1 curvature (Fig. 5B,C). Additionally, System H_2_O decreased C1 curvature but not C1 head angle, compared to 1x E3 (Fig. 5B,C), indicating a possible specific effect on tail bending. 1X E3 MB-, Ringer’s- and Hi Ca^2+^ Ringer’s-raised larvae traveled less distance during SLC responses compared to 1x E3 (Fig. 5D). 1x E3 MB, 1x E2 MB, 0.5x E2 MB, Ringer’s, Hi Ca^2+^ Ringer’s, and Bath solution larvae all displayed reduced maximum angular velocities (Fig. 5E), further illustrating specific kinematic differences resulting from the choice of media. Finally, larvae reared in 1x E3 MB performed weaker counter-bends (C2), while Bath solution and System H_2_O increased C2 angle (Fig. 5F). The media-specific variation in SLC kinematic parameters suggests that the embryo medium can selectively influence the development and/or function of motor pathways that determine the performance of the response.

**Figure 5.**
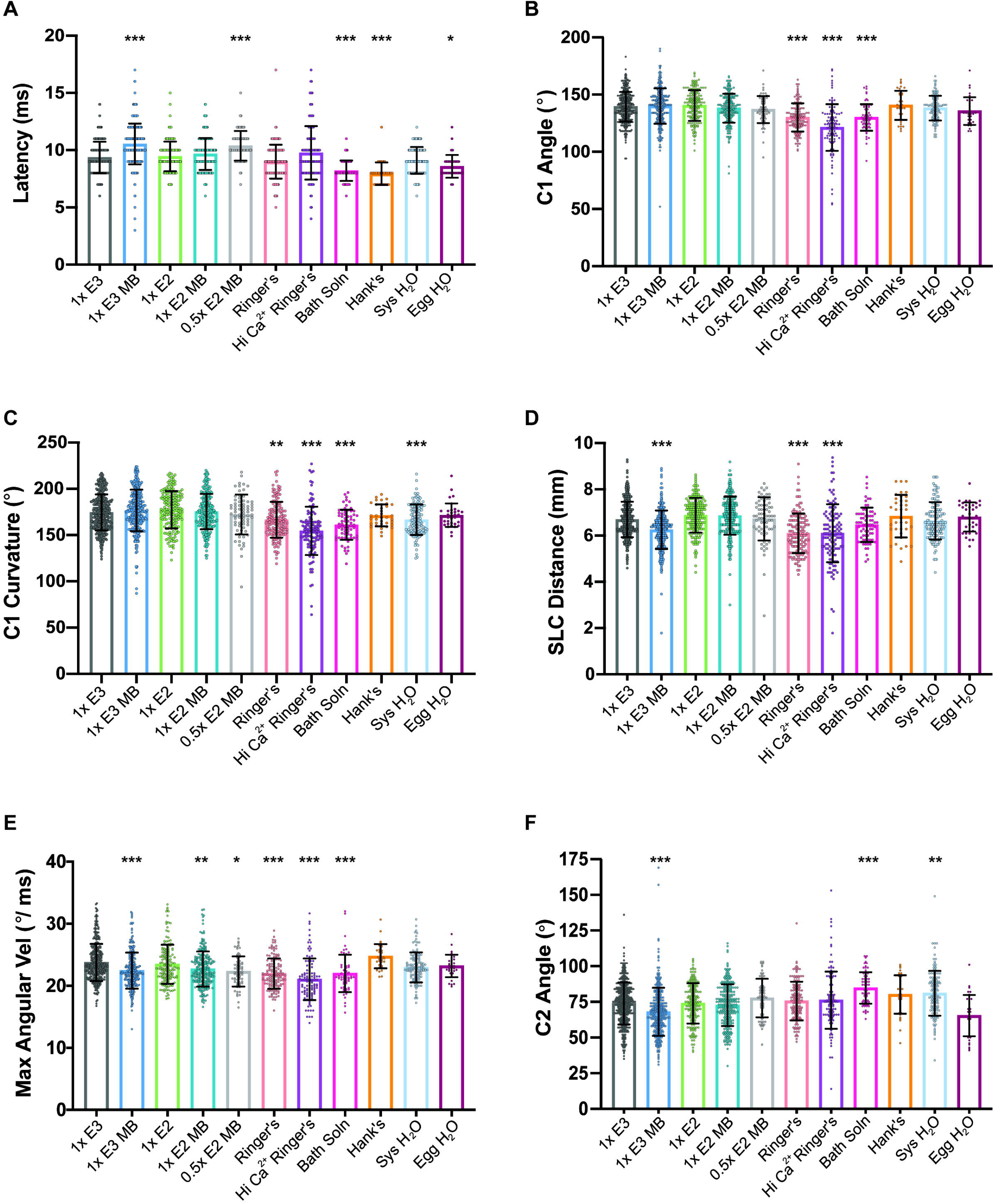
SLC kinematics compared across media types. **(A)** SLC response kinematics of individual larvae including latency of response initiation **(B)** C1 bend angle, **(C)** C1 bend curvature, **(D)** average distance traveled during response, **(E)** maximum angular velocity of C1 bend, and **(F)** C2 bend angle (1x E3 n= 380, 1x E3 MB n=314, 1x E2 n=180, 1x E2 MB n=250, 0.5x E2 MB n=70, Ringer’s n=150, Hi Ca^2+^ Ringer’s n =120, Bath Soln n=68, Hank’s n=36, Sys H_2_O n=138, Egg H_2_O n=36). Asterisks represent statistical significance for media type compared to 1x E3 (mean ± SD, Wilcoxon/ Kruskal-Wallis tests with Wilcoxon Each Pair for nonparametric multiple comparisons, *p<0.01, **p<0.001, ***p<0.0001).

### Embryo medium alters spontaneous locomotor activity and behaviors

To further investigate media-specific influences on general locomotion, we recorded spontaneous locomotion and analyzed the frequency of swim and turn movements^12, 29^ and total distance traveled (Fig. 6A). Normalized to 1x E3 larvae, larvae raised in 1x E2, Hank’s, and Egg H_2_O larvae were hyperactive and displayed greater distance traveled, while 0.5x E2 MB, Ringer’s, Hi Ca^2+^ Ringer’s, Bath solution, and System H_2_O larvae were hypoactive during the recorded period (Fig. 6B). Larvae reared in 1x E3 MB displayed significantly reduced swim and turn frequencies (Fig. 6C,D), and while larvae in 1x E2 moved significantly farther, their swim and turn frequencies were similar to 1x E3 (Fig. 6C,D), suggesting more vigorous swims/turns that propelled them farther. We also observed significantly increased swim frequencies in 0.5x E2 MB and System H_2_O larvae, while Egg H_2_O larvae displayed a reduction (Fig. 6C). Ringer’s, Hi Ca^2+^ Ringer’s, Bath solution, and Egg H_2_O larvae performed fewer turns, largely consistent with their trend of decreased total movement (Fig. 6D). Taken together, our data show that embryo medium can impact nearly every aspect of auditory-driven and locomotor behaviors that we analyzed, including startle response threshold, bias, and modulation, as well as kinematic performance and spontaneous movement initiation.

**Figure 6.**
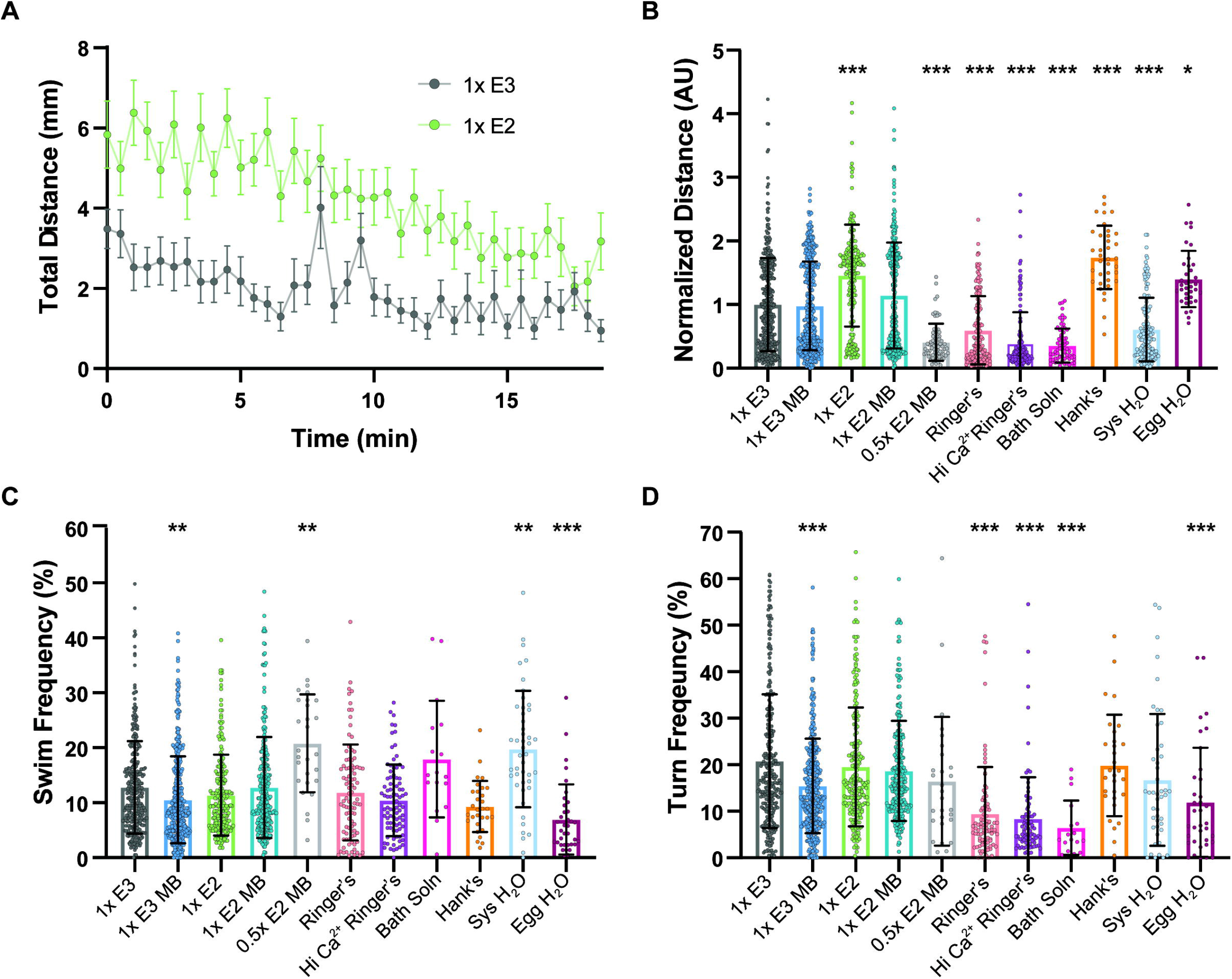
Media type induces hypo- and hyper-activity in general locomotor behaviors. **(A)** Total distance time plot during 18.5 minutes of recording in 1x E3 and 1x E2 treated larvae (mean ± SEM). **(B)** Sum of distance traveled for individual larvae, normalized to 1x E3 (1x E3 n= 398, 1x E3 MB n=382, 1x E2 n=193, 1x E2 MB n=265, 0.5x E2 MB n=72, Ringer’s n=154, Hi Ca^2+^ Ringer’s n =140, Bath Soln n=69, Hank’s n=36, Sys H_2_O n=141, Egg H_2_O n=36). **(C)** Swim and **(D)** turn frequencies. Asterisks represent statistical significance for media type compared to 1x E3 (mean ± SD, Wilcoxon/ Kruskal-Wallis tests with Wilcoxon Each Pair for nonparametric multiple comparisons, *p<0.01, **p<0.001, ***p<0.0001).

### Media type influences morphological and behavioral phenotypes in a CHARGE syndrome model

CHARGE syndrome is a rare and heterogeneous disorder characterized by a spectrum of physical and behavioral manifestations^21, 33, 34^. The majority of CHARGE cases (∼70%) arise from *de novo* loss-of-function mutations in the chromatin remodeler, *Chromodomain-helicase-DNA-binding-protein-7* (CHD7)^35, 36^. Previous work in our lab established a stable *chd7* mutant line (*chd7^ncu^*^101^*^/^*) in larval zebrafish to study the role of *chd7* in CHARGE-related sensorimotor behaviors^19^. Homozygous *chd7* mutants recapitulate CHARGE-related behavioral and morphological phenotypes such as auditory responsiveness and craniofacial, ear, and heart abnormalities, but phenotype penetrance is influenced by genetic background and/or mutation location^17^. To determine if an experimental variable such as media type can affect CHARGE-related phenotype penetrance, we reared and tested *chd7^ncu^*^101^*^/^* and wild-type siblings in 5 media types (1x E3, 1x E3 MB, 1x E2, 1x E2 MB, Hank’s). By 6 dpf, *chd7* homozygous mutants displayed multiple morphological phenotypes with varying frequencies across media types (Table 1), including craniofacial abnormalities (highest frequency: 40% 1x E2 MB), enlarged pericardial area (54% in 1x E2), uninflated swim bladder (75% in 1x E2 MB), and abnormal otoliths, or hard crystal-like structures within the otic vesicle^37, 38^ (23% in 1X E3). 58% of homozygous mutants reared in 1x E3 displayed at least one morphological phenotype while 77-81% of homozygous mutants displayed phenotypes in the other media types.

**Table 1.**
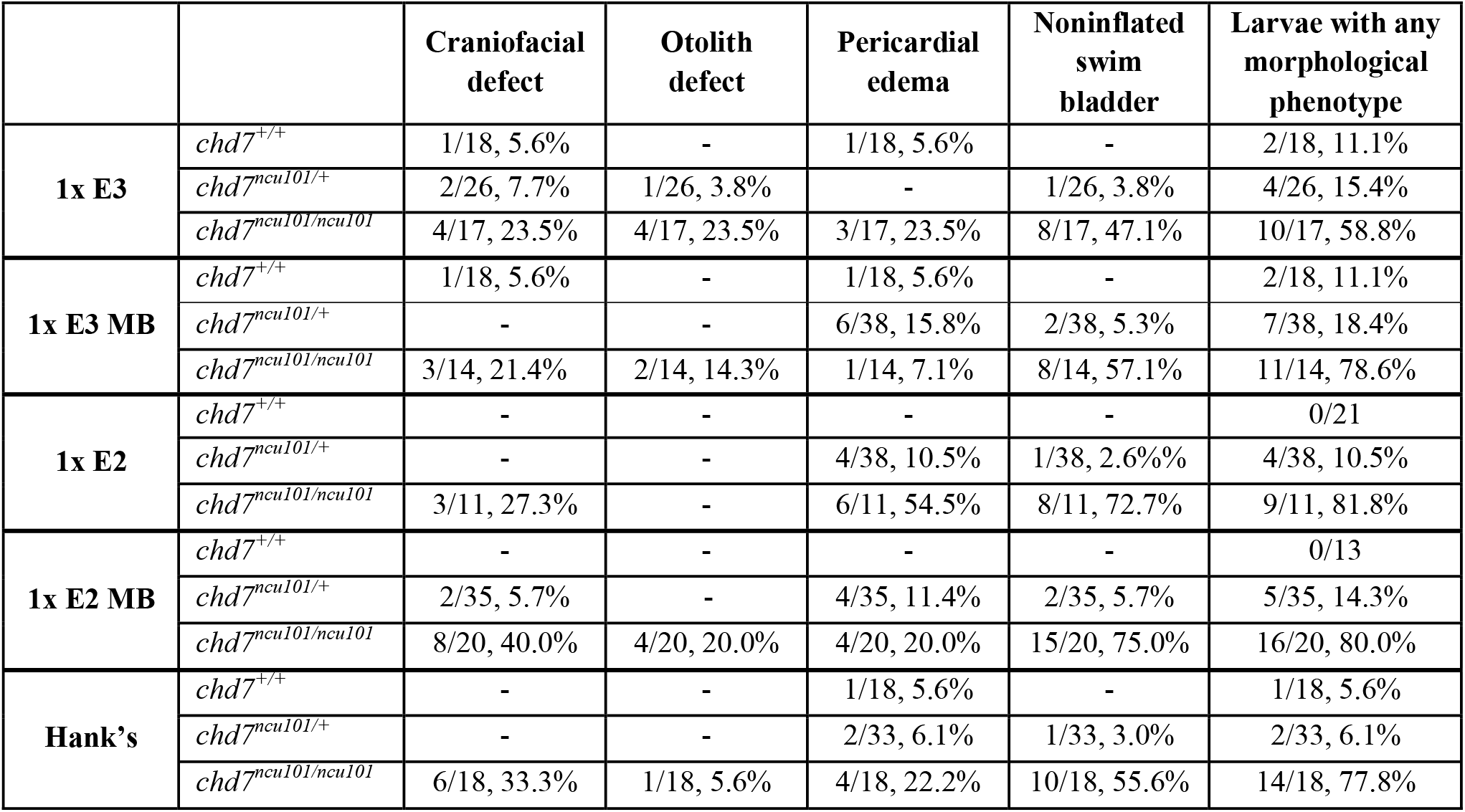
Media type influences morphological phenotype frequencies in *chd7* mutants. Morphological phenotype ratio and frequencies in 6 dpf *chd7^+/+^, chd7^ncu^*^101^*^/+,^ and chd7^ncu^*^101^*^/ncu^*^101^. Explanation and discussion of phenotype presence in wild-type siblings has been previously described ^19^.

|  |  | Craniofacial defect | Otolith defect | Pericardial edema | Noninflated swim bladder | Larvae with any morphological phenotype |
| --- | --- | --- | --- | --- | --- | --- |
| **1x E3** | *chd7<sup>+/+</sup>* | 1/18, 5.6% | - | 1/18, 5.6% | - | 2/18, 11.1% |
|  | *chd7<sup>ncu101/+</sup>* | 2/26, 7.7% | 1/26, 3.8% | - | 1/26, 3.8% | 4/26, 15.4% |
|  | *chd7<sup>ncu101/ncu101</sup>* | 4/17, 23.5% | 4/17, 23.5% | 3/17, 23.5% | 8/17, 47.1% | 10/17, 58.8% |
| **1x E3 MB** | *chd7<sup>+/+</sup>* | 1/18, 5.6% | - | 1/18, 5.6% | - | 2/18, 11.1% |
|  | *chd7<sup>ncu101/+</sup>* | - | - | 6/38, 15.8% | 2/38, 5.3% | 7/38, 18.4% |
|  | *chd7<sup>ncu101/ncu101</sup>* | 3/14, 21.4% | 2/14, 14.3% | 1/14, 7.1% | 8/14, 57.1% | 11/14, 78.6% |
| **1x E2** | *chd7<sup>+/+</sup>* | - | - | - | - | 0/21 |
|  | *chd7<sup>ncu101/+</sup>* | - | - | 4/38, 10.5% | 1/38, 2.6% | 4/38, 10.5% |
|  | *chd7<sup>ncu101/ncu101</sup>* | 3/11, 27.3% | - | 6/11, 54.5% | 8/11, 72.7% | 9/11, 81.8% |
| **1x E2 MB** | *chd7<sup>+/+</sup>* | - | - | - | - | 0/13 |
|  | *chd7<sup>ncu101/+</sup>* | 2/35, 5.7% | - | 4/35, 11.4% | 2/35, 5.7% | 5/35, 14.3% |
|  | *chd7<sup>ncu101/ncu101</sup>* | 8/20, 40.0% | 4/20, 20.0% | 4/20, 20.0% | 15/20, 75.0% | 16/20, 80.0% |
| **Hank's** | *chd7<sup>+/+</sup>* | - | - | 1/18, 5.6% | - | 1/18, 5.6% |
|  | *chd7<sup>ncu101/+</sup>* | - | - | 2/33, 6.1% | 1/33, 3.0% | 2/33, 6.1% |
|  | *chd7<sup>ncu101/ncu101</sup>* | 6/18, 33.3% | 1/18, 5.6% | 4/18, 22.2% | 10/18, 55.6% | 14/18, 77.8% |

To determine the impact of media type on previously described behavioral phenotypes, we performed the acoustic startle response and spontaneous locomotor assays on *chd7^ncu^*^101^*^/^*and wild-type siblings. Following acoustic stimulation, *chd7^ncu^*^101^*^/ncu^*^101^ show normal SLC response frequency but reduced LLC response frequency^19^, so we selected 5 media to test based on their influence on the acoustic startle response (Fig. 1). Wild-type siblings and homozygous mutants performed SLCs at similar rates regardless of media type (Fig. 7A,B), but mutants performed significantly fewer LLCs when reared in 1x E3 MB and 1x E2 MB, compared to wild-type siblings (Fig. 7C). Homozygous mutants in all other media groups displayed a trend toward reduced LLCs that did not reach statistical significance at the p<0.05 level compared to wild-type siblings. We did find that when we compared wild-types between media groups (ex. 1x E3 wild-type vs. 1x E3 MB wild-type), we observed the greatest number of differences as compared to other genotypes (Table S4). This media-dependent shift in responsiveness of wild-types thus might mask the reduced LLC phenotype in *chd7* mutants. We saw similar results in spontaneous locomotor behaviors in which hyperactivity was seen in mutants reared in 1x E3 compared to wild-type siblings (Fig. 7D), but there was a high degree of variability between the same genotypes (Table S4). Mutants in 1x E3 MB also performed more swims (Fig. 7E), while mutants in 1x E2 MB performed fewer turns (Fig. 7F) compared to wild-type siblings, phenotypes previously not reported in the *chd7^ncu^*^101^*^/^* line. Taken together, these data show that media type influences phenotype penetrance and may mask behavioral differences through genotype-specific effects in a zebrafish disease model.

**Figure 7.**
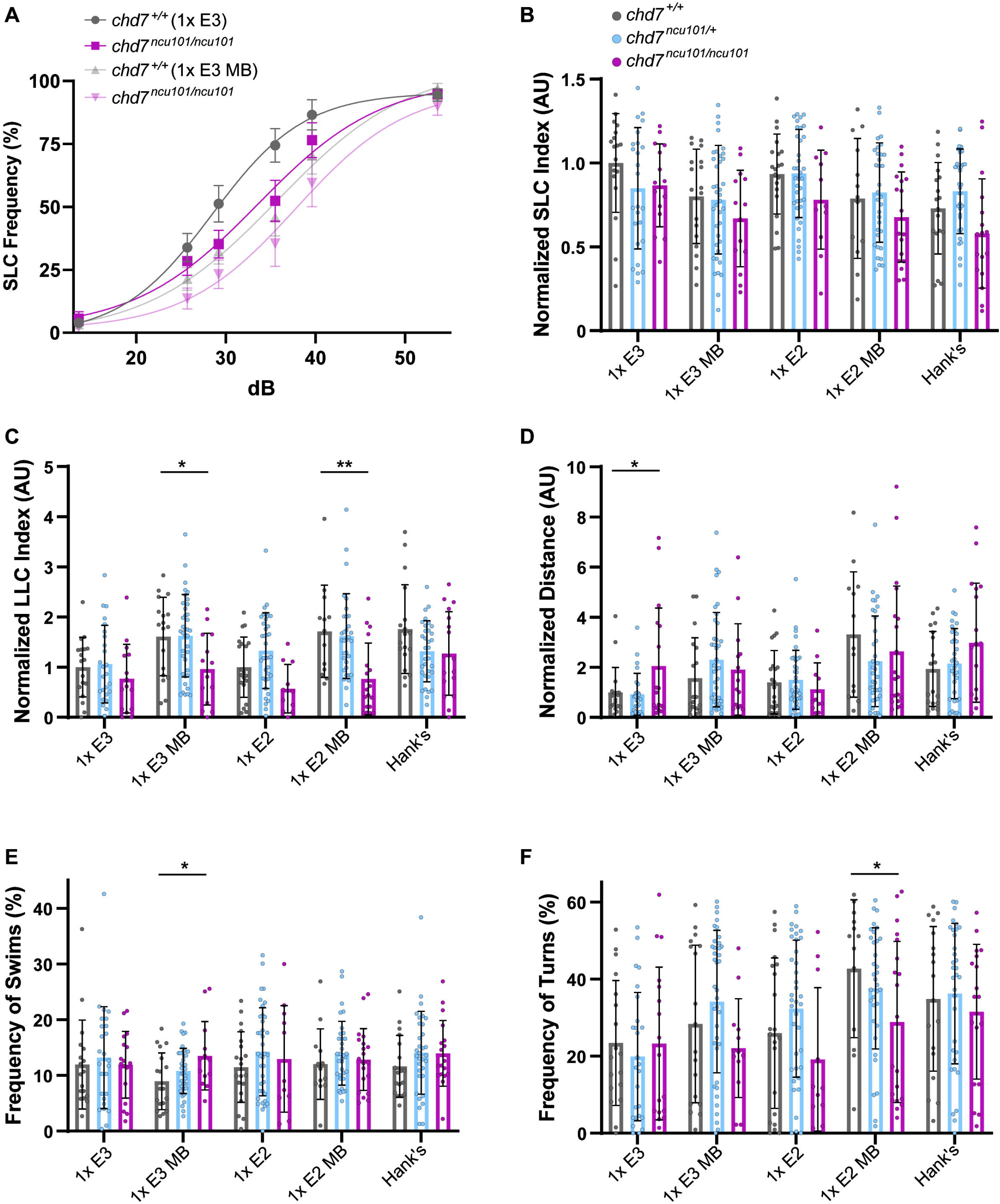
Media type impacts behavioral phenotype penetrance in *chd7* mutants. **(A)** Acoustic startle responses, average short-latency c-bend (SLC) frequency of *chd7*^+/+^ and *chd7^ncu^*^101^*^/ncu^*^101^ treated in 1x E3 (*chd7^+/+^* n=18, *chd7^ncu^*^101^*^/ncu^*^101^ n=15) or 1x E3 with methylene blue (*chd7^+/+^* n=18, *chd7^ncu^*^101^*^/ncu^*^101^ n=14) (mean ± SEM). **(B)** Short-latency c-bend and **(C)** Long-latency c-bend sensitivity indices, calculated by the area under the SLC or LLC frequency curves, respectively, for individual larvae, normalized to *chd7^+/+^* [1x E3]. Asterisks represent statistical significance for *chd7^ncu^*^101^*^/+, ncu^*^101^*^/ncu^*^101^ compared to wild-type siblings within media type (1x E3: *chd7^+/+^* n=18*, chd7^ncu^*^101^*^/+^* n=25, *chd7^ncu^*^101^*^/ncu^*^101^ n=15; 1x E3 MB: *chd7^+/+^*n=18*, chd7ncu101/+* n=38, *chd7ncu101/ncu101* n=14; 1x E2: *chd7+/+* n=20*, chd7ncu101/+* n=36, *chd7ncu101/ncu101* n=11; 1x E2 MB: *chd7^+/+^* n=13*, chd7^ncu^*^101^*^/+^* n=33, *chd7^ncu^*^101^*^/ncu^*^101^ n=18; Hank’s: *chd7^+/+^* n=18*, chd7^ncu^*^101^*^/+^* n=33, *chd7^ncu^*^101^*^/ncu^*^101^ n=17). **(D)** Spontaneous locomotion, sum of total distance traveled, normalized to *chd7^+/+^* [1x E3], **(E)** swim and **(F)** turn frequencies for individual larvae. Asterisks represent statistical significance for *chd7^ncu^*^101^*^/+, ncu^*^101^*^/ncu^*^101^ compared to wild-type siblings within media type (1x E3: *chd7^+/+^* n=18*, chd7^ncu^*^101^*^/+^*n=26, *chd7^ncu^*^101^*^/ncu^*^101^ n=17; 1x E3 MB: *chd7^+/+^*n=18*, chd7^ncu^*^101^*^/+^*n=38, *chd7^ncu^*^101^*^/ncu^*^101^ n=14; 1x E2: *chd7^+/+^* n=21*, chd7^ncu^*^101^*^/+^* n=38, *chd7^ncu^*^101^*^/ncu^*^101^ n=11; 1x E2 MB: *chd7^+/+^*n=13*, chd7^ncu^*^101^*^/+^*n=26, *chd7^ncu^*^101^*^/ncu^*^101^ n=20; Hank’s: *chd7^+/+^* n=18*, chd7^ncu^*^101^*^/+^* n=33, *chd7^ncu^*^101^*^/ncu^*^101^ n=18) (mean ± SD, One-way ANOVA with student’s t each pair test for multiple comparisons, *p<0.05, **p<0.01).

## Discussion

In this study we sought to define the impact of experimental variables such as testing arena size and embryo media type on wild-type and mutant larval zebrafish behavior. As the use of the larval zebrafish continues to expand in multiple fields in which behavior is often used as an experimental output including neuroscience, toxicology, and developmental biology, it is imperative to understand the influence experimental factors may have on results. In this study, we have demonstrated arena size- and media-dependent differences in auditory and locomotor behaviors of larval zebrafish. Additionally, media type influences the expression of both morphological and behavioral phenotypes in a zebrafish CHARGE syndrome model in a genotype-specific manner. These results highlight the substantial impacts that these variables can have on experimental outcomes, and they underscore the importance of carefully designing studies to mitigate these impacts to ensure rigor and reproducibility in the zebrafish field.

A major advantage of the larval zebrafish model is that it is amenable for high-throughput testing, particularly in toxicological and neurobehavioral research^6, 14^. These studies typically require small-diameter wells and use locomotor behavior as an output for measuring toxicity. A few studies have reported significant changes in locomotion and activity levels when arena size increased, while exposure concentrations and other variables remained constant^17, 27, 28, 39^. Another study reported significant changes in locomotion and an interaction between well size and the concentration of a neurostimulant^28^. In these studies locomotion of larvae was tested in dim light for an extended perior^39^, or in variations of the visuomotor response (VMR) assay in which locomotion is analyzed during repeated cycles of light and dark illumination^17, 27^. Although our study did not include this assay, our spontaneous locomotion data are consistent with these in showing that locomotion increases as arena size increases. Here we have expanded on this finding to demonstrate how arena size can also influence behavioral responses in multiple sensory modalities, highlighting the sensitivity of these behaviors to methodological variations.

In the acoustic startle response assay, larvae tested in the smallest arena size (9 mm diameter) were the most responsive, although this difference was exclusively for SLC and not LLC responses. It is not clear what may cause this hyperresponsiveness, but one possible explanation is that larvae may be more likely to be in contact with the wall of the smaller arena and therefore could receive both tactile and auditory stimulation. C2 bend angle was also decreased for larvae tested in the smallest arena. This is likely due to the confined space such that when larvae are close to the wall when the stimulus arrives, typically the C1 bend will be away from the wall and the C2 bend will turn larvae back toward the wall, sometimes hitting the wall and interrupting the full bend. Larvae in larger arenas showed hyperactivity and increased turn frequencies, but thigmotaxis behavior was only decreased in the medium-sized arenas (13 and 18 mm). Thigmotaxis, or “wall-hugging” behavior has been previously characterized in 5 dpf zebrafish larvae and is thought to be a measure of exploratory behavior and anxiety-like states^32^. Larvae in the 9 mm arenas exhibit strong thigmotaxis most likely because their overall activity is reduced and their space is limited, rather than resulting from an “anxiety-like” choice to stay at the perimeter and not explore the center. For the largest arena size (28 mm), though, overall activity was increased while thigmotaxis was equivalent to that seen in the smallest arena. This heightened activity while maintaining proximity to the wall indicates a clear preference for avoiding the large open field. Taken together, these data indicate how arena size influences both spontaneous locomotor activity and stimulus-evoked responses. Overall, larger arenas may be more reliable for studies examining stimulus-evoked and spontaneous locomotor behaviors, as these exert fewer constraints on movement, although this would come at the expense of reduced throughput.

Another experimental factor we considered was the type of media in which larvae were raised. We selected 11 media that are commonly used in zebrafish laboratories. Methylene blue is a synthetic cationic compound with a range of clinical and research applications, including as an antifungal agent and biocide in aquaculture^40, 41^. It is biologically active in zebrafish, and its toxicity in zebrafish cell cultures has been previously described^41^. We observed significant differences in auditory and locomotor behaviors in larvae raised in 1x E3 medium containing methylene blue (1x E3 MB) compared to the 1x E3 control. Larvae raised in 1x E3 MB were hyporesponsive in the acoustic startle response assay and showed increased pre-pulse inhibition (PPI). Similar trends were observed when comparing 1x E2 and 0.5x E2 with methylene blue. These data are consistent with a role for methylene blue in suppressing the function of the SLC circuit. Methylene blue also caused significant changes in SLC kinematic performance (increased latency, reduced angular velocity), and these overall weakened responses also reflect suppressed activity in the startle circuit. A previous study found that 4 dpf larvae treated with varying concentrations of methylene blue were hypo-active during the VMR assay^18^, which further illustrates the influence methylene blue has on stimulus-evoked behaviors and their underlying circuits. A more recent study, however, did not report any effects of methylene blue exposure from 1-4 dpf on distance traveled in the VMR assay by 6 dpf larvae^42^. This discrepancy could be because methylene blue was not present during testing and/or because it was used in 10% Hanks media, a combination which we did not test. We did observe that methylene blue impacted some behaviors but not others when in 1 x E3, and that it had a different set of impacts in 1x E2. It is unlikely that the effects we observed were due to a higher dosage of methylene blue, since the concentration of methylene blue we used (1.5 uM) was at the low end of those tested by Hedge et al. (0.6-10 uM)^42^. On balance it appears clear that methylene blue does affect some larval zebrafish behaviors, and that this may be dependent on the embryo medium used, timing of exposure, and the behavioral analysis methods. By using a high-speed camera we were able to detect subtle differences in movement that may not have been identified in other studies. Since methylene blue acts as an antifungal agent and biocide, it is possible that altered microbial makeup could contribute to the observed behavioral differences between fish raised with and without methylene blue. However, we did not observe any fungal growth in any of our media conditions. Zebrafish larvae raised in germ-free conditions have been shown to display hyperactivity and modulation of stress-like behaviors^43^, but the mechanisms driving behavioral differences in the presence or absence of causative microbes remain undetermined.

The presence of sodium bicarbonate (NaHCO_3_) and monopotassium phosphate (KH_2_PO_4_) define the E2 medium, compared to E3. The impact of sodium bicarbonate and phosphate buffers have long been studied in the optimization of cell culturing techniques including chick tissue and mouse embryos^44, 45^. The presence of NaHCO_3_ significantly increases the confluence of cells^44^, while higher concentrations of KH_2_PO_4_ is detrimental to embryo development^45, 46^. Relatively low concentrations seem to be the best environment for *in vitro* cell growth, but the effects of NaHCO_3_ past embryonic development *in vivo* are not well defined. E2 media seemingly had no effect on the acoustic startle response and kinematics in zebrafish larvae but did influence both pre-pulse inhibition and short-term habituation. Additionally, we observed hyperactivity in 1x E2 raised larvae compared to 1x E3. Although we cannot determine which component of E2 medium drives these behavioral differences, a contributing factor may be pH. E2 medium has a slightly lower pH than E3 (pH 7.6 vs. pH 7.8), but changes in extra- and intracellular H^+^ can impact neuronal function^47^. Alternatively, embryonic NaCOH_3_ exposure may have a developmental effect on behavior-specific circuits, rather than globally influencing sensorimotor behaviors.

Ringer’s and High Calcium Ringer’s contain higher Ca^2+^ concentrations compared to E2 and E3 media. Extracellular Ca^2+^ influences neuronal function and plays critical roles in synaptic transmission, neuronal survival, and axon growth^48^. Since decreased extracellular Ca^2+^ generally increases neuronal excitability^49^, it was not surprising to see multiple behavioral differences in higher Ca^2+^ containing solutions like High Calcium Ringer’s. Larvae reared in High Calcium Ringer’s performed fewer SLCs in the acoustic startle assay, and their responses were less kinematically robust. Higher Ca^2+^ concentrations caused an increase in pre-pulse inhibition but had no effect on short-term habituation, demonstrating the distinct regulatory mechanisms for these two types of behavioral modulation. Both Ringer’s and High Calcium Ringer’s caused a significant reduction in distance traveled and turn frequency compared to 1x E3, indicating they also impact the function of downstream motor pathways.

A possible explanation for the differences we observed is that some media differentially affect external sensory organ function, including hair cells of the ear and lateral line^50^. However, we did not observe consistent changes across all auditory responses with any one media type, and so the selective differences we observed in distinct auditory-driven behaviors (SLCs and LLCs) and in the modulation of those behaviors during pre-pulse inhibition and short-term habituation indicate that media more likely impacts the development and/or function of specific behavioral circuits rather than that of hair cells. Additionally, the behavior most impacted by media type was locomotion, with 8 out of 11 altering total distance traveled relative to 1x E3. This robust change in response could reflect that distributed networks across the brain drive these behaviors, but it could also be a cautionary note to researchers about the sensitivity of this assay to different conditions.

The media and arena size experiments used wild-type TLF-strain larvae, but because larval zebrafish have become a powerful model for human genetic conditions, we also sought to define how embryo media might affect a mutant line with defined disease-related morphological and behavioral phenotypes. The *chd7* mutant line (*chd7^ncu^*^101^*^/^*) has previously been characterized to have variable phenotype penetrance^19^, similar to the heterogeneous disorder it models, CHARGE syndrome^51–53^. Media type largely increases behavioral variance between wild-type genotypes, which highlights how media may mask certain mutant behavioral phenotypes. We also observed robust differences in specific morphological phenotypes in *chd7* mutants and in the overall penetrance of morphological phenotypes between media groups. CHARGE syndrome is a very heterogeneous disorder, and external factors such as retinoic acid exposure and vitamin D deficiency can produce symptoms similar to those associated with CHARGE^53–55^. In our study methylene blue exposure may similarly contribute to morphological differences in *chd7* mutants. A previous study found impaired t-cell development and possible immunological defects in *chd7* morphant and knock-out models^56^. The use of methylene blue or other biocide agents should be carefully considered when screening mutant lines with suspected immunological or developmental phenotypes.

## Conclusion

Larval zebrafish behavior is a versatile and powerful readout of neural development and function, but there are no standardized methods for raising larvae and testing their behavior. In this study, we have demonstrated that two key experimental variables, embryo media and testing arena size, can exert significant effects on the performance of multiple conserved auditory and locomotor behaviors. We also show how different media can impact the expression of behavioral and developmental phenotypes in a zebrafish disease model. Our data add to a growing body of literature that highlights the importance of not only taking these experimental variables into account in designing larval zebrafish behavior studies but also of clearly reporting these methodological details to ensure rigor and reproducibility of results across laboratories.

## Acknowledgements

We would like to thank Kim Scofield and Jackie McCawley for their additional help with behavior testing, morphological phenotyping, and genotyping of the *chd7^ncu^*^101^*^/^* line.

## Author contributions

**D.R.H.:** Formal analysis, Investigation, Visualization, Writing - Original draft, Writing - Review & editing. **T.F.H.:** Methodology, Investigation, Formal analysis. **D.B.:** Resources, Investigation, Formal analysis, Visualization. **K.N.:** Investigation, Formal analysis. **R.B.:** Investigation, Formal analysis, **V.C.:** Methodology, Investigation, Formal analysis. **K.C.M.:** Conceptualization, Supervision, Writing - Review & editing, Funding acquisition.

## Data availability

All data is available upon request.

