## Supplemental Information for "Effects of 4 testing arena sizes and 11 types of embryo media on sensorimotor behaviors in wild-type and *chd7* mutant zebrafish larvae"

| **Media** | **Components** | **Abbreviation** |
| --- | --- | --- |
| 1x E3 | 5 mM NaCl, 0.17 mM KCl, 0.33 mM CaCl_2 *_ 2H_2_O,  0.33 mM MgSO_4_ | - |
| 1x E3, 0.05% methylene blue | 5 mM NaCl, 0.17 mM KCl, 0.33 mM CaCl_2 *_ 2H_2_O,  0.33 mM MgSO4, 0.5 mg/L methylene blue | 1x E3 MB |
| 1x E2 | 15 mM NaCl, 0.5 mM KCl, 1.0 mM MgSO_4_,  150 μM KH_2_PO_4_, 50 μM Na_2_HPO_4_, 1.0 mM CaCl_2,_ 0.7 mM NaHCO_3_ | - |
| 1x E2, 0.05% methylene blue | 15 mM NaCl, 0.5 mM KCl, 1.0 mM MgSO_4_, 150 μM KH_2_PO_4_,  50 μM Na_2_HPO_4_, 1.0 mM CaCl_2,_ 0.7 mM NaHCO_3_,  0.5 mg/L methylene blue | 1x E2 MB |
| 0.5x E2, 0.05% methylene blue | 7.5 mM NaCl, 0.25 mM KCl, 0.5 mM MgSO_4_, 75μM KH_2_PO_4_,  25 μM Na_2_HPO_4_, 0.5 mM CaCl_2,_ 0.35 mM NaHCO_3_,  0.5 mg/L methylene blue | 0.5x E2 MB |
| Normal Ringer’s | 116 mM NaCl, 2.9 mM KCl, 1.8 mM CaCl_2_, 5.0 mM HEPES | Ringer’s |
| High calcium Ringer’s | 116 mM NaCl, 2.9 mM KCl, 10.0 mM CaCl_2_, 5.0 mM HEPES | Hi Ca^2+^ Ringer’s |
| Bath solution | 112 mM NaCl, 5 mM HEPES, 2 mM CaCl_2_, 3 mM Glucose,  2 mM KCl, 1 mM MgCl_2_ | Bath soln |
| 10% Hank’s | 0.137 M NaCl, 5.4 mM KCl, 0.25 mM Na_2_HPO_4_, 0.44 mM KH_2_PO_4_, 1.3 mM CaCl_2_, 1.0 mM MgSO_4,_ 4.2 mM NaHCO_3_ | Hank’s |
| Egg water | 60μg/mL “Instant Ocean” in dH_2_O | Egg H_2_O |
| System water | pH= 7.59, conductivity- 595 μS, nitrate- 5 ppm, and  nitrite- 0 ppm | Sys H_2_O |

**Supplemental Table 1.** **List of media types and solute concentrations.** For each media type, pH was measured and ranged between 7.5-8.3. For system water, specific water quality details are listed.

| **SLC Index** | | | | | | |
| --- | --- | --- | --- | --- | --- | --- |
| **Comparison** | **(n)** | **p-Value** | **Mean Dif** | **Std Error Dif** | **Lower CL** | **Upper CL** |
| 9 mm – 13 mm | 72, 64 | **0.0023** | 302.4097 | 97.835 | 109.591 | 495.2284 |
| 9 mm– 18 mm | 72, 51 | **<.0001** | 479.1065 | 104.2275 | 273.689 | 684.5239 |
| 9 mm – 28 mm | 72, 36 | **0.0018** | 367.5389 | 116.2453 | 138.436 | 596.6416 |
| 13 mm – 18 mm | 64, 51 | 0.0998 | 176.6968 | 106.8944 | -33.977 | 387.3702 |
| 13 mm – 28 mm | 64, 36 | 0.5836 | 65.1292 | 118.6424 | -168.698 | 298.9562 |
| 18 mm – 28 mm | 51, 36 | 0.5836 | 65.1292 | 118.6424 | -168.698 | 298.9562 |
| **LLC Index** | | | | | | |
| **Comparison** | **(n)** | **p-Value** | **Mean Dif** | **Std Error Dif** | **Lower CL** | **Upper CL** |
| 9 mm – 13 mm | 70, 64 | 0.5816 | 43.213 | 78.29139 | -111.096 | 197.5219 |
| 9 mm– 18 mm | 70, 51 | **0.0112** | 213.2141 | 83.34109 | 48.952 | 377.4757 |
| 9 mm – 28 mm | 70, 36 | 0.9928 | 0.8441 | 92.84387 | -182.147 | 183.8353 |
| 13 mm – 18 mm | 64, 51 | **0.0029** | 256.4271 | 84.97173 | 88.951 | 423.9026 |
| 13 mm – 28 mm | 64, 36 | 0.6409 | 44.0571 | 94.31035 | -141.824 | 229.9387 |
| 18 mm – 28 mm | 51, 36 | **0.0323** | 212.3699 | 98.54257 | 18.147 | 406.593 |
| **Latency** | | | | | | |
| **Comparison** | **(n)** | **p-Value** | **Mean Dif** | **Std Error Dif** | **Lower CL** | **Upper CL** |
| 9 mm – 13 mm | 71, 64 | 0.9424 | 0.017606 | 0.24341 | -0.4621 | 0.497307 |
| 9 mm– 18 mm | 71, 54 | 0.1571 | 0.362024 | 0.254988 | -0.1405 | 0.864543 |
| 9 mm – 28 mm | 71, 36 | 0.8753 | 0.045383 | 0.288936 | -0.52404 | 0.614806 |
| 13 mm – 18 mm | 64, 54 | 0.1471 | 0.37963 | 0.260942 | -0.13462 | 0.893883 |
| 13 mm – 28 mm | 64, 36 | 0.9249 | 0.027778 | 0.294204 | -0.55203 | 0.607583 |
| 18 mm – 28 mm | 54, 36 | 0.1814 | 0.407407 | 0.303853 | -0.19141 | 1.006227 |
| **C1 Angle** | | | | | | |
| **Comparison** | **(n)** | **p-Value** | **Mean Dif** | **Std Error Dif** | **Lower CL** | **Upper CL** |
| 9 mm – 13 mm | 71, 64 | **0.0149** | 5.475132 | 2.232163 | 1.07608 | 9.87418 |
| 9 mm– 18 mm | 71, 54 | 0.4838 | 1.640063 | 2.338338 | -2.96823 | 6.24836 |
| 9 mm – 28 mm | 71, 36 | 0.2418 | 3.109937 | 2.649657 | -2.11189 | 8.33177 |
| 13 mm – 18 mm | 64, 54 | 0.1104 | 3.835069 | 2.392943 | -0.88084 | 8.55098 |
| 13 mm – 28 mm | 64, 36 | **0.0017** | 8.585069 | 2.697969 | 3.26803 | 13.90211 |
| 18 mm – 28 mm | 54, 36 | 0.0897 | 4.75 | 2.786451 | -0.74142 | 10.24142 |
| **C1 Curvature** | | | | | | |
| **Comparison** | **(n)** | **p-Value** | **Mean Dif** | **Std Error Dif** | **Lower CL** | **Upper CL** |
| 9 mm – 13 mm | 71, 64 | 0.4739 | 2.552377 | 3.557781 | -4.45914 | 9.5639 |
| 9 mm– 18 mm | 71, 54 | 0.513 | 2.441836 | 3.727009 | -4.90319 | 9.78686 |
| 9 mm – 28 mm | 71, 36 | 0.7265 | 1.478873 | 4.223213 | -6.84405 | 9.8018 |
| 13 mm – 18 mm | 64, 54 | 0.1917 | 4.994213 | 3.814044 | -2.52234 | 12.51076 |
| 13 mm – 28 mm | 64, 36 | 0.3495 | 4.03125 | 4.300216 | -4.44343 | 12.50593 |
| 18 mm – 28 mm | 54, 36 | 0.8285 | 0.962963 | 4.441244 | -7.78965 | 9.71557 |
| **SLC Distance** | | | | | | |
| **Comparison** | **(n)** | **p-Value** | **Mean Dif** | **Std Error Dif** | **Lower CL** | **Upper CL** |
| 9 mm – 13 mm | 70, 64 | 0.4049 | 0.100446 | 0.120375 | -0.13679 | 0.337682 |
| 9 mm– 18 mm | 70, 54 | **0.0352** | 0.267113 | 0.126063 | 0.018668 | 0.515559 |
| 9 mm – 28 mm | 70, 36 | 0.5776 | 0.079613 | 0.14275 | -0.20172 | 0.360945 |
| 13 mm – 18 mm | 64, 54 | 0.1964 | 0.166667 | 0.128611 | -0.0868 | 0.420133 |
| 13 mm – 28 mm | 64, 36 | 0.8859 | 0.020833 | 0.145004 | -0.26494 | 0.306609 |
| 18 mm – 28 mm | 54, 36 | 0.2119 | 0.1875 | 0.14976 | -0.10765 | 0.482648 |
| **C1 Max Angular Velocity** | | | | | | |
| **Comparison** | **(n)** | **p-Value** | **Mean Dif** | **Std Error Dif** | **Lower CL** | **Upper CL** |
| 9 mm – 13 mm | 71, 64 | **0.0013** | 4.695095 | 1.443905 | 1.84951 | 7.540679 |
| 9 mm– 18 mm | 71, 54 | **0.0016** | 4.830029 | 1.512585 | 1.84909 | 7.810966 |
| 9 mm – 28 mm | 71, 36 | **0.0055** | 4.808204 | 1.713967 | 1.43039 | 8.186014 |
| 13 mm – 18 mm | 64, 54 | 0.9306 | 0.134934 | 1.547908 | -2.91561 | 3.185483 |
| 13 mm – 28 mm | 64, 36 | 0.9484 | 0.113109 | 1.745218 | -3.32629 | 3.552508 |
| 18 mm – 28 mm | 54, 36 | 0.9903 | 0.021825 | 1.802453 | -3.53037 | 3.574021 |
| **C2 Angle** | | | | | | |
| **Comparison** | **(n)** | **p-Value** | **Mean Dif** | **Std Error Dif** | **Lower CL** | **Upper CL** |
| 9 mm – 13 mm | 71, 64 | **0.0002** | 8.94124 | 2.327049 | 4.3552 | 13.52729 |
| 9 mm– 18 mm | 71, 54 | **<.0001** | 14.4314 | 2.437737 | 9.6272 | 19.23559 |
| 9 mm – 28 mm | 71, 36 | **<.0001** | 17.36659 | 2.76229 | 11.9228 | 22.81039 |
| 13 mm – 18 mm | 64, 54 | **0.0288** | 5.49016 | 2.494664 | 0.5738 | 10.40654 |
| 13 mm – 28 mm | 64, 36 | **0.0031** | 8.42535 | 2.812656 | 2.8823 | 13.96841 |
| 18 mm – 28 mm | 54, 36 | 0.3134 | 2.93519 | 2.904899 | -2.7897 | 8.66003 |
| **Total Distance (spontaneous)** | | | | | | |
| **Comparison** | **(n)** | **p-Value** | **Mean Dif** | **Std Error Dif** | **Lower CL** | **Upper CL** |
| 9 mm – 13 mm | 30, 30 | **<.0001** | 124.68 | 23.39292 | 78.2516 | 171.1085 |
| 9 mm– 18 mm | 30, 26 | **<.0001** | 100.2328 | 24.27598 | 52.0517 | 148.4139 |
| 9 mm – 28 mm | 30, 15 | **0.0056** | 81.234 | 28.65036 | 24.371 | 138.097 |
| 13 mm – 18 mm | 30, 26 | 0.3164 | 24.4472 | 24.27598 | -23.7339 | 72.6283 |
| 13 mm – 28 mm | 30, 15 | 0.1327 | 43.446 | 28.65036 | -13.417 | 100.309 |
| 18 mm – 28 mm | 26, 15 | 0.5193 | 18.9988 | 29.3758 | -39.304 | 77.3016 |
| **Swim Frequency** | | | | | | |
| **Comparison** | **(n)** | **p-Value** | **Mean Dif** | **Std Error Dif** | **Lower CL** | **Upper CL** |
| 9 mm – 13 mm | 36, 32 | **0.0293** | 1.926042 | 0.872447 | 0.19706 | 3.655026 |
| 9 mm– 18 mm | 36, 26 | 0.9663 | 0.039103 | 0.924206 | -1.79246 | 1.870661 |
| 9 mm – 28 mm | 36, 20 | **<.0001** | 4.541667 | 1.001471 | 2.55699 | 6.526347 |
| 13 mm – 18 mm | 32, 26 | **0.0405** | 1.965144 | 0.948119 | 0.08619 | 3.844094 |
| 13 mm – 28 mm | 32, 20 | **0.012** | 2.615625 | 1.023581 | 0.58713 | 4.644123 |
| 18 mm – 28 mm | 26, 20 | **<.0001** | 4.580769 | 1.068041 | 2.46416 | 6.697376 |
| **Turn Frequency** | | | | | | |
| **Comparison** | **(n)** | **p-Value** | **Mean Dif** | **Std Error Dif** | **Lower CL** | **Upper CL** |
| 9 mm – 13 mm | 36, 32 | **0.0007** | 6.099306 | 1.740799 | 2.6498 | 9.54881 |
| 9 mm– 18 mm | 36, 27 | **0.0143** | 4.541667 | 1.824137 | 0.92702 | 8.15632 |
| 9 mm – 28 mm | 36, 20 | **0.0002** | 7.785556 | 1.998242 | 3.82591 | 11.74521 |
| 13 mm – 18 mm | 32, 27 | 0.4072 | 1.557639 | 1.872361 | -2.15257 | 5.26785 |
| 13 mm – 28 mm | 32, 20 | 0.4108 | 1.68625 | 2.042359 | -2.36082 | 5.73332 |
| 18 mm – 28 mm | 27, 20 | 0.1277 | 3.243889 | 2.113841 | -0.94483 | 7.43261 |
| **Thigmotaxis** | | | | | | |
| **Comparison** | **(n)** | **p-Value** | **Mean Dif** | **Std Error Dif** | **Lower CL** | **Upper CL** |
| 9 mm – 13 mm | 36, 32 | **<.0001** | 20.18532 | 4.163798 | 11.9311 | 28.43956 |
| 9 mm– 18 mm | 36, 27 | **<.0001** | 30.55153 | 4.363134 | 21.9021 | 39.20093 |
| 9 mm – 28 mm | 36, 16 | 0.6301 | 2.4872 | 5.149343 | -7.7208 | 12.69517 |
| 13 mm – 18 mm | 32, 27 | **0.0225** | 10.36621 | 4.478479 | 1.4881 | 19.24427 |
| 13 mm – 28 mm | 32, 16 | **0.001** | 17.69812 | 5.247435 | 7.2957 | 28.10055 |
| 18 mm – 28 mm | 27, 16 | **<.0001** | 28.06433 | 5.406967 | 17.3456 | 38.78301 |

**Supplemental Table 2.** **Multiple comparisons between testing arena sizes.** Multiple comparisons and their reported values for the acoustic startle response (SLC and LLC Index), SLC kinematics (latency, C1 angle, C1 curvature, SLC distance, C1 max angular velocity, and C2 angle), and general locomotor behaviors (total distance, swim and turn frequency) with statistically significant p-values in bold (α=0.05, ANOVA with student’s t each pair test for multiple comparisons).

| **Normalized SLC Index** | | | | | | |
| --- | --- | --- | --- | --- | --- | --- |
| **Comparison** | **(n)** | **p-Value** | **Mean Dif** | **Std Error Dif** | **Lower CL** | **Upper CL** |
| 1x E3 – 1x E3 MB | 397, 245 | **<.0001** | 0.243781 | 0.030774 | 0.164416 | 0.323145 |
| 1x E3 – 1x E2 | 397,166 | 0.075 | 0.062367 | 0.03501 | -0.02792 | 0.152657 |
| 1x E3 – 1x E2 MB | 397, 231 | **0.0059** | 0.086472 | 0.031345 | 0.005634 | 0.16731 |
| 1x E3 – 0.5x E2 MB | 397, 64 | **<.0001** | 0.334055 | 0.051022 | 0.202471 | 0.465639 |
| 1x E3 – Ringer’s | 397, 141 | 0.8716 | 0.006004 | 0.037135 | -0.08977 | 0.101773 |
| 1x E3 – Hi Ca^2+^ Ringer’s | 397, 79 | **<.0001** | 0.269536 | 0.046665 | 0.149189 | 0.389882 |
| 1x E3 – Bath Soln | 397, 61 | 0.1079 | 0.083797 | 0.052091 | -0.05055 | 0.218138 |
| 1x E3 – Hank’s | 397, 36 | **<.0001** | 0.258751 | 0.065931 | 0.088717 | 0.428785 |
| 1x E3 – Sys H_2_O | 397, 123 | 0.9049 | 0.004673 | 0.039088 | -0.09614 | 0.10548 |
| 1x E3 – Egg H_2_O | 397, 36 | **0.0045** | 0.187649 | 0.065931 | 0.017615 | 0.357682 |
| 1x E3 MB – 1x E2 | 245, 166 | **<.0001** | 0.306147 | 0.038078 | 0.207945 | 0.404349 |
| 1x E3 MB – 1x E2 MB | 245, 231 | **<.0001** | 0.157309 | 0.034738 | 0.06772 | 0.246897 |
| 1x E3 MB – 0.5x E2 MB | 245, 64 | 0.0898 | 0.090274 | 0.053174 | -0.04686 | 0.227408 |
| 1x E3 MB – Ringer’s | 245, 141 | **<.0001** | 0.249784 | 0.04004 | 0.146522 | 0.353046 |
| 1x E3 MB – Hi Ca^2+^ Ringer’s | 245, 79 | 0.5993 | 0.025755 | 0.049008 | -0.10064 | 0.152146 |
| 1x E3 MB – Bath Soln | 245, 61 | **<.0001** | 0.327577 | 0.054201 | 0.187795 | 0.467359 |
| 1x E3 MB – Hank’s | 245, 36 | **<.0001** | 0.502531 | 0.06761 | 0.328167 | 0.676896 |
| 1x E3 MB – Sys H_2_O | 245, 123 | **<.0001** | 0.248453 | 0.041858 | 0.140502 | 0.356404 |
| 1x E3 MB – Egg H_2_O | 245, 36 | **<.0001** | 0.431429 | 0.06761 | 0.257065 | 0.605793 |
| 1x E2 – 1x E2 MB | 166, 231 | **0.0001** | 0.148838 | 0.038541 | 0.049441 | 0.248235 |
| 1x E2 – 0.5x E2 MB | 166, 64 | **<.0001** | 0.396421 | 0.055733 | 0.252688 | 0.540155 |
| 1x E2 – Ringer’s | 166, 141 | 0.194 | 0.056363 | 0.043381 | -0.05552 | 0.168241 |
| 1x E2 – Hi Ca^2+^ Ringer’s | 166, 79 | **<.0001** | 0.331902 | 0.051774 | 0.19838 | 0.465425 |
| 1x E2 – Bath Soln | 166, 61 | 0.7056 | 0.02143 | 0.056714 | -0.12483 | 0.167693 |
| 1x E2 – Hank’s | 166, 36 | **0.0049** | 0.196384 | 0.069641 | 0.016783 | 0.375985 |
| 1x E2 – Sys H_2_O | 166, 123 | 0.2006 | 0.057694 | 0.045065 | -0.05853 | 0.173914 |
| 1x E2 – Egg H_2_O | 166, 36 | 0.0722 | 0.125282 | 0.069641 | -0.05432 | 0.304883 |
| 1x E2 MB – 0.5x E2 MB | 231, 64 | **<.0001** | 0.247583 | 0.053507 | 0.109591 | 0.385575 |
| 1x E2 MB – Ringer’s | 231, 141 | 0.0225 | 0.092476 | 0.040481 | -0.01192 | 0.196874 |
| 1x E2 MB – Hi Ca^2+^ Ringer’s | 231, 79 | **0.0002** | 0.183064 | 0.049369 | 0.055743 | 0.310385 |
| 1x E2 MB – Bath Soln | 231, 61 | **0.0018** | 0.170269 | 0.054527 | 0.029645 | 0.310892 |
| 1x E2 MB – Hank’s | 231, 36 | **<.0001** | 0.345223 | 0.067872 | 0.170183 | 0.520262 |
| 1x E2 MB – Sys H_2_O | 231, 123 | 0.0313 | 0.091144 | 0.04228 | -0.01789 | 0.200183 |
| 1x E2 MB – Egg H_2_O | 231, 36 | **<.0001** | 0.27412 | 0.067872 | 0.099081 | 0.44916 |
| 0.5x E2 MB – Ringer’s | 64, 141 | **<.0001** | 0.340059 | 0.057091 | 0.192822 | 0.487295 |
| 0.5x E2 MB – Hi Ca^2+^ Ringer’s | 64, 79 | 0.3113 | 0.064519 | 0.063703 | -0.09977 | 0.228806 |
| 0.5x E2 MB – Bath Soln | 64, 61 | **<.0001** | 0.417852 | 0.067779 | 0.243053 | 0.59265 |
| 0.5x E2 MB – Hank’s | 64, 36 | **<.0001** | 0.592806 | 0.078913 | 0.389291 | 0.796321 |
| 0.5x E2 MB – Sys H_2_O | 64, 123 | **<.0001** | 0.338728 | 0.058381 | 0.188165 | 0.48929 |
| 0.5x E2 MB – Egg H_2_O | 64, 36 | **<.0001** | 0.521703 | 0.078913 | 0.318188 | 0.725219 |
| Ringer’s – Hi Ca^2+^ Ringer’s | 141, 79 | **<.0001** | 0.27554 | 0.053233 | 0.138253 | 0.412826 |
| Ringer’s – Bath Soln | 141, 61 | 0.1804 | 0.077793 | 0.058049 | -0.07191 | 0.227499 |
| Ringer’s – Hank’s | 141, 36 | **0.0004** | 0.252747 | 0.070732 | 0.070331 | 0.435164 |
| Ringer’s – Sys H_2_O | 141, 123 | 0.9773 | 0.001331 | 0.046734 | -0.11919 | 0.121856 |
| Ringer’s – Egg H_2_O | 141, 36 | 0.0103 | 0.181645 | 0.070732 | -0.00077 | 0.364061 |
| Hi Ca^2+^ Ringer’s – Bath Soln | 79, 61 | **<.0001** | 0.353333 | 0.064562 | 0.186829 | 0.519836 |
| Hi Ca^2+^ Ringer’s – Hank’s | 79, 36 | **<.0001** | 0.528287 | 0.076169 | 0.33185 | 0.724723 |
| Hi Ca^2+^ Ringer’s – Sys H_2_O | 79, 123 | **<.0001** | 0.274208 | 0.054614 | 0.133361 | 0.415056 |
| Hi Ca^2+^ Ringer’s – Egg H_2_O | 79, 36 | **<.0001** | 0.457184 | 0.076169 | 0.260748 | 0.653621 |
| Bath Soln – Hank’s | 61, 36 | 0.0281 | 0.174954 | 0.079609 | -0.03036 | 0.380263 |
| Bath Soln – Sys H_2_O | 61, 123 | 0.1824 | 0.079124 | 0.059318 | -0.07385 | 0.232102 |
| Bath Soln – Egg H_2_O | 61, 36 | 0.1922 | 0.103852 | 0.079609 | -0.10146 | 0.309161 |
| Hank’s – Sys H_2_O | 36, 123 | **0.0004** | 0.254078 | 0.071777 | 0.068967 | 0.43919 |
| Hank’s – Egg H_2_O | 36, 36 | 0.4259 | 0.071102 | 0.08928 | -0.15915 | 0.301354 |
| Sys H_2_O – Egg H_2_O | 123, 36 | 0.0109 | 0.182976 | 0.071777 | -0.00214 | 0.368087 |
| **Normalized LLC Index** | | | | | | |
| **Comparison** | **(n)** | **p-Value** | **Mean Dif** | **Std Error Dif** | **Lower CL** | **Upper CL** |
| 1x E3 – 1x E3 MB | 377, 250 | 0.3468 | 0.04935 | 0.052439 | -0.08589 | 0.184586 |
| 1x E3 – 1x E2 | 377, 166 | 0.1743 | 0.081388 | 0.059887 | -0.07306 | 0.235834 |
| 1x E3 – 1x E2 MB | 377, 246 | **<.0001** | 0.212599 | 0.052695 | 0.076703 | 0.348494 |
| 1x E3 – 0.5x E2 MB | 377, 67 | **0.0007** | 0.289513 | 0.08524 | 0.069684 | 0.509341 |
| 1x E3 – Ringer’s | 377, 146 | 0.0136 | 0.154758 | 0.062671 | -0.00687 | 0.316381 |
| 1x E3 – Hi Ca^2+^ Ringer’s | 377, 80 | **0.0022** | 0.242634 | 0.079141 | 0.038534 | 0.446735 |
| 1x E3 – Bath Soln | 377, 64 | **0.002** | 0.268794 | 0.08692 | 0.044633 | 0.492955 |
| 1x E3 – Hank’s | 377, 36 | 0.5899 | 0.060453 | 0.112154 | -0.22878 | 0.34969 |
| 1x E3 – Sys H_2_O | 377, 129 | 0.073 | 0.117647 | 0.06558 | -0.05148 | 0.286773 |
| 1x E3 – Egg H_2_O | 377, 36 | 0.6079 | 0.057554 | 0.112154 | -0.23168 | 0.346791 |
| 1x E3 MB – 1x E2 | 250, 166 | 0.0424 | 0.130738 | 0.06437 | -0.03527 | 0.296744 |
| 1x E3 MB – 1x E2 MB | 250, 246 | **<.0001** | 0.261948 | 0.057738 | 0.113045 | 0.410852 |
| 1x E3 MB – 0.5x E2 MB | 250, 67 | **0.0001** | 0.338863 | 0.088447 | 0.110764 | 0.566962 |
| 1x E3 MB – Ringer’s | 250, 146 | 0.1157 | 0.105408 | 0.066967 | -0.0673 | 0.278112 |
| 1x E3 MB – Hi Ca^2+^ Ringer’s | 250, 80 | **0.0004** | 0.291984 | 0.082585 | 0.079003 | 0.504966 |
| 1x E3 MB – Bath Soln | 250, 64 | 0.0149 | 0.219444 | 0.090067 | -0.01283 | 0.451721 |
| 1x E3 MB – Hank’s | 250, 36 | 0.3382 | 0.109803 | 0.11461 | -0.18577 | 0.405374 |
| 1x E3 MB – Sys H_2_O | 250, 129 | 0.3273 | 0.068297 | 0.069697 | -0.11145 | 0.248042 |
| 1x E3 MB – Egg H_2_O | 250, 36 | 0.9429 | 0.008204 | 0.11461 | -0.28737 | 0.303775 |
| 1x E2 – 1x E2 MB | 166, 246 | 0.0423 | 0.13121 | 0.064578 | -0.03533 | 0.297754 |
| 1x E2 – 0.5x E2 MB | 166, 67 | 0.0255 | 0.208125 | 0.093057 | -0.03186 | 0.448111 |
| 1x E2 – Ringer’s | 166, 146 | **0.0012** | 0.236146 | 0.072947 | 0.048021 | 0.424272 |
| 1x E2 – Hi Ca^2+^ Ringer’s | 166, 80 | 0.0656 | 0.161246 | 0.087504 | -0.06442 | 0.386914 |
| 1x E2 – Bath Soln | 166, 64 | **0.0002** | 0.350182 | 0.094598 | 0.106221 | 0.594143 |
| 1x E2 – Hank’s | 166, 36 | 0.8594 | 0.020935 | 0.118204 | -0.2839 | 0.325775 |
| 1x E2 – Sys H_2_O | 166, 129 | **0.0084** | 0.199036 | 0.075461 | 0.004427 | 0.393645 |
| 1x E2 – Egg H_2_O | 166, 36 | 0.24 | 0.138942 | 0.118204 | -0.1659 | 0.443782 |
| 1x E2 MB – 0.5x E2 MB | 246, 67 | 0.3855 | 0.076914 | 0.088599 | -0.15158 | 0.305405 |
| 1x E2 MB – Ringer’s | 246, 146 | **<.0001** | 0.367356 | 0.067168 | 0.194136 | 0.540577 |
| 1x E2 MB – Hi Ca^2+^ Ringer’s | 246, 80 | 0.7167 | 0.030036 | 0.082748 | -0.18337 | 0.243437 |
| 1x E2 MB – Bath Soln | 246, 64 | **<.0001** | 0.481392 | 0.090216 | 0.248731 | 0.714054 |
| 1x E2 MB – Hank’s | 246, 36 | 0.185 | 0.152146 | 0.114727 | -0.14373 | 0.448019 |
| 1x E2 MB – Sys H_2_O | 246, 129 | **<.0001** | 0.330246 | 0.06989 | 0.150005 | 0.510487 |
| 1x E2 MB – Egg H_2_O | 246, 36 | 0.0187 | 0.270152 | 0.114727 | -0.02572 | 0.566026 |
| 0.5x E2 MB – Ringer’s | 67, 146 | **<.0001** | 0.444271 | 0.094872 | 0.199603 | 0.688938 |
| 0.5x E2 MB – Hi Ca^2+^ Ringer’s | 67, 80 | 0.6598 | 0.046878 | 0.106472 | -0.22771 | 0.321464 |
| 0.5x E2 MB – Bath Soln | 67, 64 | **<.0001** | 0.558307 | 0.112375 | 0.2685 | 0.848114 |
| 0.5x E2 MB – Hank’s | 67, 36 | 0.0849 | 0.22906 | 0.132859 | -0.11357 | 0.571694 |
| 0.5x E2 MB – Sys H_2_O | 67, 129 | **<.0001** | 0.40716 | 0.096818 | 0.157473 | 0.656848 |
| 0.5x E2 MB – Egg H_2_O | 67, 36 | **0.0091** | 0.347067 | 0.132859 | 0.004433 | 0.689701 |
| Ringer’s – Hi Ca^2+^ Ringer’s | 146, 80 | **<.0001** | 0.397392 | 0.089432 | 0.166753 | 0.628032 |
| Ringer’s – Bath Soln | 146, 64 | 0.2369 | 0.114036 | 0.096384 | -0.13453 | 0.362603 |
| Ringer’s – Hank’s | 146, 36 | 0.0722 | 0.215211 | 0.119638 | -0.09333 | 0.523749 |
| Ringer’s – Sys H_2_O | 146, 129 | 0.6329 | 0.03711 | 0.077688 | -0.16324 | 0.237463 |
| Ringer’s – Egg H_2_O | 146, 36 | 0.4166 | 0.097204 | 0.119638 | -0.21133 | 0.405742 |
| Hi Ca^2+^ Ringer’s – Bath Soln | 80, 64 | **<.0001** | 0.511428 | 0.107822 | 0.233363 | 0.789494 |
| Hi Ca^2+^ Ringer’s – Hank’s | 80, 36 | 0.1582 | 0.182182 | 0.129031 | -0.15058 | 0.514943 |
| Hi Ca^2+^ Ringer’s – Sys H_2_O | 80, 129 | **<.0001** | 0.360282 | 0.091494 | 0.124324 | 0.596239 |
| Hi Ca^2+^ Ringer’s – Egg H_2_O | 80, 36 | 0.0201 | 0.300188 | 0.129031 | -0.03257 | 0.63295 |
| Bath Soln – Hank’s | 64, 36 | 0.0141 | 0.329247 | 0.133943 | -0.01618 | 0.674676 |
| Bath Soln – Sys H_2_O | 64, 129 | 0.1243 | 0.151147 | 0.0983 | -0.10236 | 0.404656 |
| Bath Soln – Egg H_2_O | 64, 36 | 0.115 | 0.21124 | 0.133943 | -0.13419 | 0.55667 |
| Hank’s – Sys H_2_O | 36, 129 | 0.1419 | 0.1781 | 0.121187 | -0.13443 | 0.490634 |
| Hank’s – Egg H_2_O | 36, 36 | 0.4363 | 0.118007 | 0.151539 | -0.2728 | 0.508816 |
| Sys H_2_O – Egg H_2_O | 129, 36 | 0.6201 | 0.060094 | 0.121187 | -0.25244 | 0.372627 |
| **Pre-pulse Inhibition** | | | | | | |
| **Comparison** | **(n)** | **p-Value** | **Mean Dif** | **Std Error Dif** | **Lower CL** | **Upper CL** |
| 1x E3 – 1x E3 MB | 263, 118 | **<.0001** | 15.79586 | 3.606363 | 6.4894 | 25.1023 |
| 1x E3 – 1x E2 | 263, 108 | **<.0001** | 18.23723 | 3.719829 | 8.638 | 27.83648 |
| 1x E3 – 1x E2 MB | 263, 137 | 0.5551 | 2.02453 | 3.429396 | -6.8252 | 10.8743 |
| 1x E3 – 0.5x E2 MB | 263, 63 | 0.02 | 10.63711 | 4.56548 | -1.1444 | 22.41861 |
| 1x E3 – Ringer’s | 263, 115 | 0.0553 | 6.98002 | 3.638689 | -2.4098 | 16.36987 |
| 1x E3 – Hi Ca^2+^ Ringer’s | 263, 62 | **<.0001** | 21.86984 | 4.595087 | 10.0119 | 33.72774 |
| 1x E3 – Bath Soln | 263, 61 | **<.0001** | 23.81997 | 4.625466 | 11.8837 | 35.75626 |
| 1x E3 – Sys H_2_O | 263, 123 | **<.0001** | 15.19922 | 3.555404 | 6.0243 | 24.37415 |
| 1x E3 MB – 1x E2 | 118, 108 | **<.0001** | 34.0331 | 4.334381 | 22.848 | 45.21823 |
| 1x E3 MB – 1x E2 MB | 118, 137 | **<.0001** | 17.8204 | 4.087847 | 7.2715 | 28.36933 |
| 1x E3 MB – 0.5x E2 MB | 118, 63 | 0.31 | 5.15875 | 5.078714 | -7.9472 | 18.26468 |
| 1x E3 MB – Ringer’s | 118, 115 | 0.039 | 8.81585 | 4.264949 | -2.1901 | 19.8218 |
| 1x E3 MB – Hi Ca^2+^ Ringer’s | 118, 62 | 0.2344 | 6.07398 | 5.105346 | -7.1007 | 19.24863 |
| 1x E3 MB – Bath Soln | 118, 61 | **<.0001** | 39.61583 | 5.132706 | 26.3706 | 52.86109 |
| 1x E3 MB – Sys H_2_O | 118, 123 | **<.0001** | 30.99508 | 4.194119 | 20.1719 | 41.81826 |
| 1x E2 – 1x E2 MB | 108, 137 | **0.0001** | 16.2127 | 4.188289 | 5.4046 | 27.02083 |
| 1x E2 – 0.5x E2 MB | 108, 63 | **<.0001** | 28.87434 | 5.159905 | 15.5589 | 42.18979 |
| 1x E2 – Ringer’s | 108, 115 | **<.0001** | 25.21725 | 4.361315 | 13.9626 | 36.47189 |
| 1x E2 – Hi Ca^2+^ Ringer’s | 108, 62 | **<.0001** | 40.10707 | 5.18612 | 26.724 | 53.49017 |
| 1x E2 – Bath Soln | 108, 61 | 0.2845 | 5.58273 | 5.213055 | -7.8699 | 19.03534 |
| 1x E2 – Sys H_2_O | 108, 123 | 0.4792 | 3.03802 | 4.292075 | -8.0379 | 14.11398 |
| 1x E2 MB – 0.5x E2 MB | 137, 63 | 0.0107 | 12.66165 | 4.954619 | -0.124 | 25.44734 |
| 1x E2 MB – Ringer’s | 137, 115 | 0.0289 | 9.00455 | 4.116393 | -1.6181 | 19.62715 |
| 1x E2 MB – Hi Ca^2+^ Ringer’s | 137, 62 | **<.0001** | 23.89437 | 4.981914 | 11.0382 | 36.7505 |
| 1x E2 MB – Bath Soln | 137, 61 | **<.0001** | 21.79543 | 5.009948 | 8.867 | 34.7239 |
| 1x E2 MB – Sys H_2_O | 137, 123 | **0.0012** | 13.17468 | 4.042962 | 2.7416 | 23.60779 |
| 0.5x E2 MB – Ringer’s | 63, 115 | 0.4736 | 3.65709 | 5.10172 | -9.5082 | 16.82239 |
| 0.5x E2 MB – Hi Ca^2+^ Ringer’s | 63, 62 | 0.054 | 11.23273 | 5.822569 | -3.7928 | 26.25822 |
| 0.5x E2 MB – Bath Soln | 63, 61 | **<.0001** | 34.45708 | 5.846574 | 19.3696 | 49.54451 |
| 0.5x E2 MB – Sys H_2_O | 63, 123 | **<.0001** | 25.83633 | 5.042657 | 12.8234 | 38.84921 |
| Ringer’s – Hi Ca^2+^ Ringer’s | 115, 62 | **0.0038** | 14.88982 | 5.128232 | 1.6561 | 28.12353 |
| Ringer’s – Bath Soln | 115, 61 | **<.0001** | 30.79998 | 5.15547 | 17.496 | 44.10398 |
| Ringer’s – Sys H_2_O | 115, 123 | **<.0001** | 22.17923 | 4.221947 | 11.2842 | 33.07422 |
| Hi Ca^2+^ Ringer’s – Bath Soln | 62, 61 | **<.0001** | 45.68981 | 5.869723 | 30.5426 | 60.83698 |
| Hi Ca^2+^ Ringer’s – Sys H_2_O | 62, 123 | **<.0001** | 37.06906 | 5.069478 | 23.987 | 50.15115 |
| Bath Soln – Sys H_2_O | 61, 123 | 0.0911 | 8.62075 | 5.09703 | -4.5324 | 21.77394 |
| **Habituation Half-life** | | | | | | |
| **Comparison** | **(n)** | **p-Value** | **Mean Dif** | **Std Error Dif** | **Lower CL** | **Upper CL** |
| 1x E3 – 1x E3 MB | 163, 133 | 0.0102 | 0.285782 | 0.110966 | -0.00075 | 0.572318 |
| 1x E3 – 1x E2 | 163, 116 | **0.0009** | 0.383829 | 0.115357 | 0.08596 | 0.681702 |
| 1x E3 – 1x E2 MB | 163, 124 | **<.0001** | 0.509256 | 0.113162 | 0.21705 | 0.801461 |
| 1x E3 – 0.5x E2 MB | 163, 30 | 0.0216 | 0.434194 | 0.188664 | -0.05297 | 0.92136 |
| 1x E3 – Ringer’s | 163, 85 | 0.0823 | 0.221046 | 0.127054 | -0.10703 | 0.549123 |
| 1x E3 – Hi Ca^2+^ Ringer’s | 163, 22 | 0.0352 | 0.455014 | 0.215698 | -0.10196 | 1.011986 |
| 1x E3 – Bath Soln | 163, 23 | **<.0001** | 1.912316 | 0.211526 | 1.36612 | 2.458515 |
| 1x E3 – Hank’s | 163, 43 | **<.0001** | 0.952188 | 0.162806 | 0.53179 | 1.372583 |
| 1x E3 – Sys H_2_O | 163, 25 | 0.692 | 0.08083 | 0.203976 | -0.44587 | 0.607535 |
| 1x E3 – Egg H_2_O | 163, 22 | **<.0001** | 1.153322 | 0.215698 | 0.59635 | 1.710294 |
| 1x E3 MB – 1x E2 | 133, 116 | **<.0001** | 0.669611 | 0.120645 | 0.35808 | 0.981139 |
| 1x E3 MB – 1x E2 MB | 133, 124 | **<.0001** | 0.795038 | 0.118548 | 0.48892 | 1.101151 |
| 1x E3 MB – 0.5x E2 MB | 133, 30 | 0.4396 | 0.148412 | 0.191943 | -0.34722 | 0.644045 |
| 1x E3 MB – Ringer’s | 133, 85 | 0.6236 | 0.064736 | 0.131874 | -0.27579 | 0.405258 |
| 1x E3 MB – Hi Ca^2+^ Ringer’s | 133, 22 | 0.439 | 0.169232 | 0.218571 | -0.39516 | 0.733624 |
| 1x E3 MB – Bath Soln | 133, 23 | **<.0001** | 2.198098 | 0.214456 | 1.64433 | 2.751863 |
| 1x E3 MB – Hank’s | 133, 43 | **<.0001** | 1.23797 | 0.166595 | 0.80779 | 1.668149 |
| 1x E3 MB – Sys H_2_O | 133, 25 | 0.3225 | 0.204952 | 0.207013 | -0.32959 | 0.739498 |
| 1x E3 MB – Egg H_2_O | 133, 22 | **<.0001** | 1.439104 | 0.218571 | 0.87471 | 2.003497 |
| 1x E2 – 1x E2 MB | 116, 124 | 0.3069 | 0.125427 | 0.122668 | -0.19132 | 0.442178 |
| 1x E2 – 0.5x E2 MB | 116, 30 | **<.0001** | 0.818023 | 0.194514 | 0.31575 | 1.320295 |
| 1x E2 – Ringer’s | 116, 85 | **<.0001** | 0.604875 | 0.135589 | 0.25476 | 0.954991 |
| 1x E2 – Hi Ca^2+^ Ringer’s | 116, 22 | **0.0002** | 0.838843 | 0.220833 | 0.26861 | 1.409075 |
| 1x E2 – Bath Soln | 116, 23 | **<.0001** | 1.528487 | 0.21676 | 0.96877 | 2.088202 |
| 1x E2 – Hank’s | 116, 43 | **0.0008** | 0.568359 | 0.169551 | 0.13055 | 1.006171 |
| 1x E2 – Sys H_2_O | 116, 25 | 0.0268 | 0.464659 | 0.209399 | -0.07605 | 1.005367 |
| 1x E2 – Egg H_2_O | 116, 22 | **0.0005** | 0.769493 | 0.220833 | 0.19926 | 1.339725 |
| 1x E2 MB – 0.5x E2 MB | 124, 30 | **<.0001** | 0.94345 | 0.193221 | 0.44452 | 1.442382 |
| 1x E2 MB – Ringer’s | 124, 85 | **<.0001** | 0.730302 | 0.133727 | 0.385 | 1.075609 |
| 1x E2 MB – Hi Ca^2+^ Ringer’s | 124, 22 | **<.0001** | 0.96427 | 0.219694 | 0.39698 | 1.531562 |
| 1x E2 MB – Bath Soln | 124, 23 | **<.0001** | 1.40306 | 0.2156 | 0.84634 | 1.95978 |
| 1x E2 MB – Hank’s | 124, 43 | **0.0086** | 0.442932 | 0.168065 | 0.00896 | 0.876908 |
| 1x E2 MB – Sys H_2_O | 124, 25 | **0.0047** | 0.590086 | 0.208198 | 0.05248 | 1.127693 |
| 1x E2 MB – Egg H_2_O | 124, 22 | **0.0035** | 0.644066 | 0.219694 | 0.07677 | 1.211358 |
| 0.5x E2 MB – Ringer’s | 30, 85 | 0.2909 | 0.213148 | 0.201671 | -0.3076 | 0.733901 |
| 0.5x E2 MB – Hi Ca^2+^ Ringer’s | 30, 22 | 0.9378 | 0.02082 | 0.26656 | -0.66749 | 0.709127 |
| 0.5x E2 MB – Bath Soln | 30, 23 | **<.0001** | 2.34651 | 0.263195 | 1.66689 | 3.02613 |
| 0.5x E2 MB – Hank’s | 30, 43 | **<.0001** | 1.386382 | 0.225908 | 0.80305 | 1.969719 |
| 0.5x E2 MB – Sys H_2_O | 30, 25 | 0.1698 | 0.353364 | 0.257167 | -0.31069 | 1.017418 |
| 0.5x E2 MB – Egg H_2_O | 30, 22 | **<.0001** | 1.587516 | 0.26656 | 0.89921 | 2.275823 |
| Ringer’s – Hi Ca^2+^ Ringer’s | 85, 22 | 0.3033 | 0.233968 | 0.227162 | -0.35261 | 0.820543 |
| Ringer’s – Bath Soln | 85, 23 | **<.0001** | 2.133362 | 0.223205 | 1.55701 | 2.709718 |
| Ringer’s – Hank’s | 85, 43 | **<.0001** | 1.173234 | 0.177716 | 0.71434 | 1.632129 |
| Ringer’s – Sys H_2_O | 85, 25 | 0.5166 | 0.140216 | 0.216064 | -0.4177 | 0.698132 |
| Ringer’s – Egg H_2_O | 85, 22 | **<.0001** | 1.374368 | 0.227162 | 0.78779 | 1.960943 |
| Hi Ca^2+^ Ringer’s – Bath Soln | 22, 23 | **<.0001** | 2.36733 | 0.283202 | 1.63605 | 3.09861 |
| Hi Ca^2+^ Ringer’s – Hank’s | 22, 43 | **<.0001** | 1.407202 | 0.248929 | 0.76442 | 2.049984 |
| Hi Ca^2+^ Ringer’s – Sys H_2_O | 22, 25 | 0.1781 | 0.374184 | 0.277608 | -0.34265 | 1.09102 |
| Hi Ca^2+^ Ringer’s – Egg H_2_O | 22, 22 | **<.0001** | 1.608336 | 0.286331 | 0.86898 | 2.347697 |
| Bath Soln – Hank’s | 23, 43 | **<.0001** | 0.960128 | 0.245323 | 0.32666 | 1.593598 |
| Bath Soln – Sys H_2_O | 23, 25 | **<.0001** | 1.993146 | 0.274379 | 1.28465 | 2.701645 |
| Bath Soln – Egg H_2_O | 23, 22 | **0.0075** | 0.758994 | 0.283202 | 0.02771 | 1.490274 |
| Hank’s – Sys H_2_O | 43, 25 | **<.0001** | 1.033018 | 0.238844 | 0.41628 | 1.649759 |
| Hank’s – Egg H_2_O | 25, 22 | 0.4193 | 0.201134 | 0.248929 | -0.44165 | 0.843916 |
| Sys H_2_O – Egg H_2_O | 25, 22 | **<.0001** | 1.234152 | 0.277608 | 0.51732 | 1.950989 |
| **Latency** | | | | | | |
| **Comparison** | **(n)** | **p-Value** | **Mean Dif** | **Std Error Dif** | **Lower CL** | **Upper CL** |
| 1x E3 – 1x E3 MB | 380, 314 | **<.0001** | 1.183507 | 0.114471 | 0.88832 | 1.47869 |
| 1x E3 – 1x E2 | 380, 180 | 0.4977 | 0.092105 | 0.135812 | -0.25811 | 0.44232 |
| 1x E3 – 1x E2 MB | 380, 250 | 0.0124 | 0.306105 | 0.122231 | -0.00909 | 0.621299 |
| 1x E3 – 0.5x E2 MB | 380, 71 | **<.0001** | 1.022387 | 0.194062 | 0.52197 | 1.522809 |
| 1x E3 – Ringer’s | 380, 150 | 0.0119 | 0.364561 | 0.144735 | -0.00866 | 0.737785 |
| 1x E3 – Hi Ca^2+^ Ringer’s | 380, 122 | **0.0097** | 0.4044 | 0.15619 | 0.00164 | 0.807163 |
| 1x E3 – Bath Soln | 380, 68 | **<.0001** | 1.152012 | 0.197636 | 0.64237 | 1.66165 |
| 1x E3 – Hank’s | 380, 36 | **<.0001** | 1.41345 | 0.261744 | 0.7385 | 2.088402 |
| 1x E3 – Sys H_2_O | 380, 138 | 0.095 | 0.249199 | 0.149179 | -0.13548 | 0.633881 |
| 1x E3 – Egg H_2_O | 380, 36 | **0.0031** | 0.774561 | 0.261744 | 0.09961 | 1.449513 |
| 1x E3 MB – 1x E2 | 314, 180 | **<.0001** | 1.091401 | 0.140325 | 0.72955 | 1.453253 |
| 1x E3 MB – 1x E2 MB | 314, 250 | **<.0001** | 0.877401 | 0.127227 | 0.54933 | 1.205477 |
| 1x E3 MB – 0.5x E2 MB | 314, 71 | 0.4141 | 0.16112 | 0.197247 | -0.34751 | 0.669753 |
| 1x E3 MB – Ringer’s | 314, 150 | **<.0001** | 1.548068 | 0.148978 | 1.1639 | 1.932233 |
| 1x E3 MB – Hi Ca^2+^ Ringer’s | 314, 122 | **<.0001** | 0.779106 | 0.16013 | 0.36618 | 1.192027 |
| 1x E3 MB – Bath Soln | 314, 68 | **<.0001** | 2.335519 | 0.200764 | 1.81782 | 2.853222 |
| 1x E3 MB – Hank’s | 314, 36 | **<.0001** | 2.596957 | 0.264114 | 1.91589 | 3.278019 |
| 1x E3 MB – Sys H_2_O | 314, 138 | **<.0001** | 1.432706 | 0.153299 | 1.0374 | 1.828012 |
| 1x E3 MB – Egg H_2_O | 314, 36 | **<.0001** | 1.958068 | 0.264114 | 1.27701 | 2.63913 |
| 1x E2 – 1x E2 MB | 180, 250 | 0.1449 | 0.214 | 0.146724 | -0.16435 | 0.592353 |
| 1x E2 – 0.5x E2 MB | 180, 71 | **<.0001** | 0.930282 | 0.210351 | 0.38786 | 1.472707 |
| 1x E2 – Ringer’s | 180, 150 | **0.006** | 0.456667 | 0.165939 | 0.02877 | 0.884568 |
| 1x E2 – Hi Ca^2+^ Ringer’s | 180, 122 | 0.0762 | 0.312295 | 0.17602 | -0.1416 | 0.766191 |
| 1x E2 – Bath Soln | 180, 68 | **<.0001** | 1.244118 | 0.213653 | 0.69318 | 1.795057 |
| 1x E2 – Hank’s | 180, 36 | **<.0001** | 1.505556 | 0.274039 | 0.7989 | 2.212212 |
| 1x E2 – Sys H_2_O | 180, 138 | 0.0446 | 0.341304 | 0.169829 | -0.09663 | 0.779236 |
| 1x E2 – Egg H_2_O | 180, 36 | **0.0016** | 0.866667 | 0.274039 | 0.16001 | 1.573323 |
| 1x E2 MB – 0.5x E2 MB | 250, 71 | **0.0004** | 0.716282 | 0.201849 | 0.19578 | 1.236783 |
| 1x E2 MB – Ringer’s | 250, 150 | **<.0001** | 0.670667 | 0.15502 | 0.27092 | 1.070412 |
| 1x E2 MB – Hi Ca^2+^ Ringer’s | 250, 122 | 0.5533 | 0.098295 | 0.165766 | -0.32916 | 0.52575 |
| 1x E2 MB – Bath Soln | 250, 68 | **<.0001** | 1.458118 | 0.205288 | 0.92875 | 1.987486 |
| 1x E2 MB – Hank’s | 250, 36 | **<.0001** | 1.719556 | 0.267569 | 1.02959 | 2.409526 |
| 1x E2 MB – Sys H_2_O | 250, 138 | **0.0005** | 0.555304 | 0.159177 | 0.14484 | 0.965768 |
| 1x E2 MB – Egg H_2_O | 250, 36 | **<.0001** | 1.080667 | 0.267569 | 0.3907 | 1.770637 |
| 0.5x E2 MB – Ringer’s | 71, 150 | **<.0001** | 1.386948 | 0.216219 | 0.82939 | 1.944506 |
| 0.5x E2 MB – Hi Ca^2+^ Ringer’s | 71, 122 | **0.0059** | 0.617987 | 0.224049 | 0.04024 | 1.195734 |
| 0.5x E2 MB – Bath Soln | 71, 68 | **<.0001** | 2.174399 | 0.254681 | 1.51766 | 2.831138 |
| 0.5x E2 MB – Hank’s | 71, 36 | **<.0001** | 2.435837 | 0.307104 | 1.64392 | 3.227755 |
| 0.5x E2 MB – Sys H_2_O | 71, 138 | **<.0001** | 1.271586 | 0.219219 | 0.70629 | 1.836878 |
| 0.5x E2 MB – Egg H_2_O | 71, 36 | **<.0001** | 1.796948 | 0.307104 | 1.00503 | 2.588866 |
| Ringer’s – Hi Ca^2+^ Ringer’s | 150, 122 | **<.0001** | 0.768962 | 0.182992 | 0.29709 | 1.240838 |
| Ringer’s – Bath Soln | 150, 68 | **0.0003** | 0.787451 | 0.219433 | 0.22161 | 1.353295 |
| Ringer’s – Hank’s | 150, 36 | **0.0002** | 1.048889 | 0.278569 | 0.33055 | 1.767226 |
| Ringer’s – Sys H_2_O | 150, 138 | 0.5147 | 0.115362 | 0.177045 | -0.34118 | 0.571903 |
| Ringer’s – Egg H_2_O | 150, 36 | 0.1413 | 0.41 | 0.278569 | -0.30834 | 1.128337 |
| Hi Ca^2+^ Ringer’s – Bath Soln | 122, 68 | **<.0001** | 1.556413 | 0.227152 | 0.97066 | 2.142161 |
| Hi Ca^2+^ Ringer’s – Hank’s | 122, 36 | **<.0001** | 1.817851 | 0.284689 | 1.08373 | 2.551969 |
| Hi Ca^2+^ Ringer’s – Sys H_2_O | 122, 138 | **0.0005** | 0.653599 | 0.186527 | 0.17261 | 1.134589 |
| Hi Ca^2+^ Ringer’s – Egg H_2_O | 122, 36 | **<.0001** | 1.178962 | 0.284689 | 0.44484 | 1.91308 |
| Bath Soln – Hank’s | 68, 36 | 0.3982 | 0.261438 | 0.309374 | -0.53634 | 1.059212 |
| Bath Soln – Sys H_2_O | 68, 138 | **<.0001** | 0.902813 | 0.222389 | 0.32935 | 1.47628 |
| Bath Soln – Egg H_2_O | 68, 36 | 0.2226 | 0.377451 | 0.309374 | -0.42032 | 1.175225 |
| Hank’s – Sys H_2_O | 36, 138 | **<.0001** | 1.164251 | 0.280904 | 0.43989 | 1.888608 |
| Hank’s – Egg H_2_O | 36, 36 | 0.0711 | 0.638889 | 0.353783 | -0.2734 | 1.551178 |
| Sys H_2_O – Egg H_2_O | 138, 36 | 0.0616 | 0.525362 | 0.280904 | -0.19899 | 1.249719 |
| **C1 Angle** | | | | | | |
| **Comparison** | **(n)** | **p-Value** | **Mean Dif** | **Std Error Dif** | **Lower CL** | **Upper CL** |
| 1x E3 – 1x E3 MB | 380, 314 | 0.5282 | 0.66262 | 1.050266 | -2.0457 | 3.37091 |
| 1x E3 – 1x E2 | 380, 180 | 0.3903 | 1.07076 | 1.24607 | -2.1424 | 4.28397 |
| 1x E3 – 1x E2 MB | 380, 250 | 0.2699 | 1.23768 | 1.121463 | -1.6542 | 4.12957 |
| 1x E3 – 0.5x E2 MB | 380, 70 | 0.1749 | 2.43083 | 1.791191 | -2.1881 | 7.04972 |
| 1x E3 – Ringer’s | 380, 150 | **<.0001** | 9.32035 | 1.327935 | 5.896 | 12.74466 |
| 1x E3 – Hi Ca^2+^ Ringer’s | 380, 120 | **<.0001** | 18.10702 | 1.442045 | 14.3885 | 21.82558 |
| 1x E3 – Bath Soln | 380, 68 | **<.0001** | 9.33251 | 1.813298 | 4.6566 | 14.00841 |
| 1x E3 – Hank’s | 380, 36 | 0.6016 | 1.25409 | 2.401486 | -4.9386 | 7.44674 |
| 1x E3 – Sys H_2_O | 380, 138 | 0.3977 | 1.15774 | 1.368705 | -2.3717 | 4.68718 |
| 1x E3 – Egg H_2_O | 380, 36 | 0.119 | 3.74591 | 2.401486 | -2.4467 | 9.93855 |
| 1x E3 MB – 1x E2 | 314, 180 | 0.7513 | 0.40814 | 1.287475 | -2.9118 | 3.72811 |
| 1x E3 MB – 1x E2 MB | 314, 250 | 0.1037 | 1.90031 | 1.167296 | -1.1098 | 4.91038 |
| 1x E3 MB – 0.5x E2 MB | 314, 70 | 0.0894 | 3.09345 | 1.820238 | -1.6003 | 7.78725 |
| 1x E3 MB – Ringer’s | 314, 150 | **<.0001** | 9.98297 | 1.366863 | 6.4583 | 13.50766 |
| 1x E3 MB – Hi Ca^2+^ Ringer’s | 314, 120 | **<.0001** | 18.76964 | 1.477971 | 14.9584 | 22.58084 |
| 1x E3 MB – Bath Soln | 314, 68 | **<.0001** | 9.99513 | 1.841996 | 5.2452 | 14.74503 |
| 1x E3 MB – Hank’s | 314, 36 | 0.8072 | 0.59147 | 2.423228 | -5.6572 | 6.84018 |
| 1x E3 MB – Sys H_2_O | 314, 138 | 0.1958 | 1.82036 | 1.406505 | -1.8066 | 5.44728 |
| 1x E3 MB – Egg H_2_O | 314, 36 | 0.069 | 4.40853 | 2.423228 | -1.8402 | 10.65724 |
| 1x E2 – 1x E2 MB | 180, 250 | 0.0866 | 2.30844 | 1.346184 | -1.1629 | 5.77981 |
| 1x E2 – 0.5x E2 MB | 180, 70 | 0.0712 | 3.50159 | 1.939818 | -1.5006 | 8.50374 |
| 1x E2 – Ringer’s | 180, 150 | **<.0001** | 10.39111 | 1.522479 | 6.4651 | 14.31709 |
| 1x E2 – Hi Ca^2+^ Ringer’s | 180, 120 | **<.0001** | 19.17778 | 1.622969 | 14.9927 | 23.36288 |
| 1x E2 – Bath Soln | 180, 68 | **<.0001** | 10.40327 | 1.96025 | 5.3484 | 15.45811 |
| 1x E2 – Hank’s | 180, 36 | 0.9419 | 0.18333 | 2.514292 | -6.3002 | 6.66687 |
| 1x E2 – Sys H_2_O | 180, 138 | 0.1528 | 2.2285 | 1.558167 | -1.7895 | 6.24651 |
| 1x E2 – Egg H_2_O | 180, 36 | 0.0556 | 4.81667 | 2.514292 | -1.6669 | 11.3002 |
| 1x E2 MB – 0.5x E2 MB | 250, 70 | 0.5218 | 1.19314 | 1.862226 | -3.6089 | 5.99521 |
| 1x E2 MB – Ringer’s | 250, 150 | **<.0001** | 8.08267 | 1.422298 | 4.415 | 11.75031 |
| 1x E2 MB – Hi Ca^2+^ Ringer’s | 250, 120 | **<.0001** | 16.86933 | 1.529384 | 12.9256 | 20.81311 |
| 1x E2 MB – Bath Soln | 250, 68 | **<.0001** | 8.09482 | 1.883499 | 3.2379 | 12.95175 |
| 1x E2 MB – Hank’s | 250, 36 | 0.3102 | 2.49178 | 2.454924 | -3.8387 | 8.82222 |
| 1x E2 MB – Sys H_2_O | 250, 138 | 0.9564 | 0.07994 | 1.460436 | -3.686 | 3.84593 |
| 1x E2 MB – Egg H_2_O | 250, 36 | 0.3071 | 2.50822 | 2.454924 | -3.8222 | 8.83867 |
| 0.5x E2 MB – Ringer’s | 70, 150 | **0.0006** | 6.88952 | 1.993394 | 1.7492 | 12.02983 |
| 0.5x E2 MB – Hi Ca^2+^ Ringer’s | 70, 120 | **<.0001** | 15.67619 | 2.071159 | 10.3353 | 21.01703 |
| 0.5x E2 MB – Bath Soln | 70, 68 | **0.0033** | 6.90168 | 2.344836 | 0.8551 | 12.94824 |
| 0.5x E2 MB – Hank’s | 70, 36 | 0.1922 | 3.68492 | 2.824418 | -3.5983 | 10.96817 |
| 0.5x E2 MB – Sys H_2_O | 70, 138 | 0.5288 | 1.27308 | 2.020782 | -3.9379 | 6.48402 |
| 0.5x E2 MB – Egg H_2_O | 70, 36 | 0.6416 | 1.31508 | 2.824418 | -5.9682 | 8.59833 |
| Ringer’s – Hi Ca^2+^ Ringer’s | 150, 120 | **<.0001** | 8.78667 | 1.686638 | 4.4374 | 13.13595 |
| Ringer’s – Bath Soln | 150, 68 | 0.9952 | 0.01216 | 2.013282 | -5.1794 | 5.20375 |
| Ringer’s – Hank’s | 150, 36 | **<.0001** | 10.57444 | 2.555853 | 3.9837 | 17.16515 |
| Ringer’s – Sys H_2_O | 150, 138 | **<.0001** | 8.16261 | 1.624379 | 3.9739 | 12.35135 |
| Ringer’s – Egg H_2_O | 150, 36 | 0.0293 | 5.57444 | 2.555853 | -1.0163 | 12.16515 |
| Hi Ca^2+^ Ringer’s – Bath Soln | 120, 68 | **<.0001** | 8.77451 | 2.090308 | 3.3843 | 14.16473 |
| Hi Ca^2+^ Ringer’s – Hank’s | 120, 36 | **<.0001** | 19.36111 | 2.616958 | 12.6128 | 26.10939 |
| Hi Ca^2+^ Ringer’s – Sys H_2_O | 120, 138 | **<.0001** | 16.94928 | 1.718921 | 12.5167 | 21.38181 |
| Hi Ca^2+^ Ringer’s – Egg H_2_O | 120, 36 | **<.0001** | 14.36111 | 2.616958 | 7.6128 | 21.10939 |
| Bath Soln – Hank’s | 68, 36 | **0.0002** | 10.5866 | 2.83849 | 3.2671 | 17.90614 |
| Bath Soln – Sys H_2_O | 68, 138 | **<.0001** | 8.17477 | 2.040403 | 2.9132 | 13.4363 |
| Bath Soln – Egg H_2_O | 68, 36 | 0.0492 | 5.5866 | 2.83849 | -1.7329 | 12.90614 |
| Hank’s – Sys H_2_O | 36, 138 | 0.3495 | 2.41184 | 2.577272 | -4.2341 | 9.05778 |
| Hank’s – Egg H_2_O | 36, 36 | 0.1236 | 5 | 3.245937 | -3.3702 | 13.37021 |
| Sys H_2_O – Egg H_2_O | 138, 36 | 0.3154 | 2.58816 | 2.577272 | -4.0578 | 9.2341 |
| **C1 Curvature** | | | | | | |
| **Comparison** | **(n)** | **p-Value** | **Mean Dif** | **Std Error Dif** | **Lower CL** | **Upper CL** |
| 1x E3 – 1x E3 MB | 380, 314 | 0.1409 | 2.25825 | 1.532881 | -1.6945 | 6.21104 |
| 1x E3 – 1x E2 | 380, 180 | 0.1055 | 2.94561 | 1.818659 | -1.7441 | 7.63533 |
| 1x E3 – 1x E2 MB | 380, 250 | 0.5754 | 0.91695 | 1.636793 | -3.3038 | 5.13769 |
| 1x E3 – 0.5x E2 MB | 380, 71 | 0.3817 | 2.27387 | 2.598679 | -4.4273 | 8.975 |
| 1x E3 – Ringer’s | 380, 150 | **<.0001** | 7.93772 | 1.938143 | 2.9399 | 12.93555 |
| 1x E3 – Hi Ca^2+^ Ringer’s | 380, 122 | **<.0001** | 19.93007 | 2.091536 | 14.5367 | 25.32345 |
| 1x E3 – Bath Soln | 380, 68 | **<.0001** | 13.3387 | 2.646537 | 6.5142 | 20.16324 |
| 1x E3 – Hank’s | 380, 36 | 0.3582 | 3.22105 | 3.505007 | -5.8172 | 12.2593 |
| 1x E3 – Sys H_2_O | 380, 138 | **<.0001** | 7.84786 | 1.997647 | 2.6966 | 12.99913 |
| 1x E3 – Egg H_2_O | 380, 36 | 0.388 | 3.02661 | 3.505007 | -6.0116 | 12.06486 |
| 1x E3 MB – 1x E2 | 314, 180 | 0.7146 | 0.68737 | 1.87909 | -4.1582 | 5.53292 |
| 1x E3 MB – 1x E2 MB | 314, 250 | 0.4312 | 1.3413 | 1.703688 | -3.0519 | 5.73454 |
| 1x E3 MB – 0.5x E2 MB | 314, 71 | 0.0864 | 4.53212 | 2.641324 | -2.279 | 11.34321 |
| 1x E3 MB – Ringer’s | 314, 150 | **<.0001** | 10.19597 | 1.994958 | 5.0516 | 15.3403 |
| 1x E3 MB – Hi Ca^2+^ Ringer’s | 314, 122 | **<.0001** | 22.18832 | 2.144291 | 16.6589 | 27.71773 |
| 1x E3 MB – Bath Soln | 314, 68 | **<.0001** | 15.59695 | 2.688423 | 8.6644 | 22.5295 |
| 1x E3 MB – Hank’s | 314, 36 | 0.1215 | 5.4793 | 3.536741 | -3.6408 | 14.59938 |
| 1x E3 MB – Sys H_2_O | 314, 138 | **<.0001** | 10.10611 | 2.052816 | 4.8126 | 15.39964 |
| 1x E3 MB – Egg H_2_O | 314, 36 | 0.1353 | 5.28485 | 3.536741 | -3.8352 | 14.40493 |
| 1x E2 – 1x E2 MB | 180, 250 | 0.302 | 2.02867 | 1.964777 | -3.0378 | 7.09517 |
| 1x E2 – 0.5x E2 MB | 180, 71 | 0.0641 | 5.21948 | 2.816804 | -2.0441 | 12.48309 |
| 1x E2 – Ringer’s | 180, 150 | **<.0001** | 10.88333 | 2.222083 | 5.1533 | 16.61335 |
| 1x E2 – Hi Ca^2+^ Ringer’s | 180, 122 | **<.0001** | 22.87568 | 2.35707 | 16.7976 | 28.95378 |
| 1x E2 – Bath Soln | 180, 68 | **<.0001** | 16.28431 | 2.861017 | 8.9067 | 23.66192 |
| 1x E2 – Hank’s | 180, 36 | 0.093 | 6.16667 | 3.66965 | -3.2961 | 15.62947 |
| 1x E2 – Sys H_2_O | 180, 138 | **<.0001** | 10.79348 | 2.274171 | 4.9291 | 16.65781 |
| 1x E2 – Egg H_2_O | 180, 36 | 0.1038 | 5.97222 | 3.66965 | -3.4906 | 15.43503 |
| 1x E2 MB – 0.5x E2 MB | 250, 71 | 0.238 | 3.19082 | 2.702954 | -3.7792 | 10.16084 |
| 1x E2 MB – Ringer’s | 250, 150 | **<.0001** | 8.85467 | 2.075867 | 3.5017 | 14.20764 |
| 1x E2 MB – Hi Ca^2+^ Ringer’s | 250, 122 | **<.0001** | 20.84702 | 2.219764 | 15.123 | 26.57105 |
| 1x E2 MB – Bath Soln | 250, 68 | **<.0001** | 14.25565 | 2.748998 | 7.1669 | 21.3444 |
| 1x E2 MB – Hank’s | 250, 36 | 0.2483 | 4.138 | 3.583002 | -5.1014 | 13.37737 |
| 1x E2 MB – Sys H_2_O | 250, 138 | **<.0001** | 8.76481 | 2.131531 | 3.2683 | 14.26132 |
| 1x E2 MB – Egg H_2_O | 250, 36 | 0.2712 | 3.94356 | 3.583002 | -5.2958 | 13.18293 |
| 0.5x E2 MB – Ringer’s | 71, 150 | 0.0506 | 5.66385 | 2.895387 | -1.8024 | 13.13009 |
| 0.5x E2 MB – Hi Ca^2+^ Ringer’s | 71, 122 | **<.0001** | 17.6562 | 3.000232 | 9.9196 | 25.3928 |
| 0.5x E2 MB – Bath Soln | 71, 68 | **0.0012** | 11.06483 | 3.410429 | 2.2705 | 19.85919 |
| 0.5x E2 MB – Hank’s | 71, 36 | 0.8179 | 0.94718 | 4.112413 | -9.6574 | 11.55173 |
| 0.5x E2 MB – Sys H_2_O | 71, 138 | 0.0578 | 5.57399 | 2.935551 | -1.9958 | 13.14381 |
| 0.5x E2 MB – Egg H_2_O | 71, 36 | 0.8548 | 0.75274 | 4.112413 | -9.8518 | 11.35728 |
| Ringer’s – Hi Ca^2+^ Ringer’s | 150, 122 | **<.0001** | 11.99235 | 2.450441 | 5.6735 | 18.31122 |
| Ringer’s – Bath Soln | 150, 68 | 0.0662 | 5.40098 | 2.938417 | -2.1762 | 12.97818 |
| Ringer’s – Hank’s | 150, 36 | 0.2063 | 4.71667 | 3.730309 | -4.9026 | 14.33589 |
| Ringer’s – Sys H_2_O | 150, 138 | 0.9698 | 0.08986 | 2.370808 | -6.0237 | 6.20338 |
| Ringer’s – Egg H_2_O | 150, 36 | 0.1882 | 4.91111 | 3.730309 | -4.7081 | 14.53034 |
| Hi Ca^2+^ Ringer’s – Bath Soln | 122, 68 | 0.0304 | 6.59137 | 3.041779 | -1.2524 | 14.43511 |
| Hi Ca^2+^ Ringer’s – Hank’s | 122, 36 | **<.0001** | 16.70902 | 3.812261 | 6.8785 | 26.53957 |
| Hi Ca^2+^ Ringer’s – Sys H_2_O | 122, 138 | **<.0001** | 12.0822 | 2.49777 | 5.6413 | 18.52312 |
| Hi Ca^2+^ Ringer’s – Egg H_2_O | 122, 36 | **<.0001** | 16.90346 | 3.812261 | 7.0729 | 26.73402 |
| Bath Soln – Hank’s | 68, 36 | 0.0147 | 10.11765 | 4.142822 | -0.5653 | 20.80061 |
| Bath Soln – Sys H_2_O | 68, 138 | 0.0654 | 5.49084 | 2.978001 | -2.1884 | 13.17011 |
| Bath Soln – Egg H_2_O | 68, 36 | 0.0129 | 10.31209 | 4.142822 | -0.3709 | 20.99505 |
| Hank’s – Sys H_2_O | 36, 138 | 0.2189 | 4.62681 | 3.76157 | -5.073 | 14.32665 |
| Hank’s – Egg H_2_O | 36, 36 | 0.9673 | 0.19444 | 4.737498 | -12.022 | 12.41088 |
| Sys H_2_O – Egg H_2_O | 138, 36 | 0.2001 | 4.82126 | 3.76157 | -4.8786 | 14.52109 |
| **SLC Distance** | | | | | | |
| **Comparison** | **(n)** | **p-Value** | **Mean Dif** | **Std Error Dif** | **Lower CL** | **Upper CL** |
| 1x E3 – 1x E3 MB | 380, 314 | **<.0001** | 0.443746 | 0.064574 | 0.27723 | 0.610261 |
| 1x E3 – 1x E2 | 380, 180 | 0.0174 | 0.182319 | 0.076613 | -0.01524 | 0.379878 |
| 1x E3 – 1x E2 MB | 380, 250 | 0.0169 | 0.164882 | 0.068952 | -0.01292 | 0.342685 |
| 1x E3 – 0.5x E2 MB | 380, 70 | 0.7887 | 0.029348 | 0.109472 | -0.25294 | 0.31164 |
| 1x E3 – Ringer’s | 380, 150 | **<.0001** | 0.596743 | 0.081646 | 0.386205 | 0.807282 |
| 1x E3 – Hi Ca^2+^ Ringer’s | 380, 120 | **<.0001** | 0.594346 | 0.088108 | 0.367144 | 0.821547 |
| 1x E3 – Bath Soln | 380, 68 | 0.0408 | 0.228251 | 0.111488 | -0.05924 | 0.515742 |
| 1x E3 – Hank’s | 380, 36 | 0.3374 | 0.141694 | 0.147652 | -0.23905 | 0.52244 |
| 1x E3 – Sys H_2_O | 380, 138 | 0.4931 | 0.057695 | 0.084153 | -0.15931 | 0.274697 |
| 1x E3 – Egg H_2_O | 380, 36 | 0.4546 | 0.110444 | 0.147652 | -0.2703 | 0.49119 |
| 1x E3 MB – 1x E2 | 314, 180 | **<.0001** | 0.626065 | 0.079159 | 0.421941 | 0.830189 |
| 1x E3 MB – 1x E2 MB | 314, 250 | **<.0001** | 0.608627 | 0.07177 | 0.423557 | 0.793697 |
| 1x E3 MB – 0.5x E2 MB | 314, 70 | **<.0001** | 0.473094 | 0.111269 | 0.186169 | 0.760018 |
| 1x E3 MB – Ringer’s | 314, 150 | 0.0688 | 0.152998 | 0.08404 | -0.06371 | 0.369708 |
| 1x E3 MB – Hi Ca^2+^ Ringer’s | 314, 120 | 0.0957 | 0.1506 | 0.090331 | -0.08233 | 0.383532 |
| 1x E3 MB – Bath Soln | 314, 68 | 0.0572 | 0.215495 | 0.113253 | -0.07655 | 0.507536 |
| 1x E3 MB – Hank’s | 314, 36 | **<.0001** | 0.58544 | 0.148989 | 0.201247 | 0.969633 |
| 1x E3 MB – Sys H_2_O | 314, 138 | **<.0001** | 0.386051 | 0.086477 | 0.163056 | 0.609047 |
| 1x E3 MB – Egg H_2_O | 314, 36 | **0.0002** | 0.55419 | 0.148989 | 0.169997 | 0.938383 |
| 1x E2 – 1x E2 MB | 180, 250 | 0.8332 | 0.017438 | 0.082768 | -0.19599 | 0.230869 |
| 1x E2 – 0.5x E2 MB | 180, 70 | 0.1975 | 0.152971 | 0.118661 | -0.15302 | 0.458958 |
| 1x E2 – Ringer’s | 180, 150 | **<.0001** | 0.779063 | 0.093608 | 0.53768 | 1.020445 |
| 1x E2 – Hi Ca^2+^ Ringer’s | 180, 120 | **<.0001** | 0.776665 | 0.099294 | 0.520619 | 1.032711 |
| 1x E2 – Bath Soln | 180, 68 | **0.0007** | 0.41057 | 0.120523 | 0.09978 | 0.721359 |
| 1x E2 – Hank’s | 180, 36 | 0.7927 | 0.040625 | 0.154588 | -0.35801 | 0.439256 |
| 1x E2 – Sys H_2_O | 180, 138 | 0.0123 | 0.240014 | 0.095802 | -0.00703 | 0.487055 |
| 1x E2 – Egg H_2_O | 180, 36 | 0.642 | 0.071875 | 0.154588 | -0.32676 | 0.470506 |
| 1x E2 MB – 0.5x E2 MB | 250, 70 | 0.2341 | 0.135534 | 0.113865 | -0.15809 | 0.429153 |
| 1x E2 MB – Ringer’s | 250, 150 | **<.0001** | 0.761625 | 0.087448 | 0.536125 | 0.987125 |
| 1x E2 MB – Hi Ca^2+^ Ringer’s | 250, 120 | **<.0001** | 0.759228 | 0.09351 | 0.518097 | 1.000358 |
| 1x E2 MB – Bath Soln | 250, 68 | **0.0007** | 0.393132 | 0.115804 | 0.094511 | 0.691753 |
| 1x E2 MB – Hank’s | 250, 36 | 0.8779 | 0.023188 | 0.150938 | -0.36603 | 0.412406 |
| 1x E2 MB – Sys H_2_O | 250, 138 | 0.0133 | 0.222576 | 0.089793 | -0.00897 | 0.454122 |
| 1x E2 MB – Egg H_2_O | 250, 36 | 0.7184 | 0.054438 | 0.150938 | -0.33478 | 0.443656 |
| 0.5x E2 MB – Ringer’s | 70, 150 | **<.0001** | 0.626092 | 0.121971 | 0.311568 | 0.940615 |
| 0.5x E2 MB – Hi Ca^2+^ Ringer’s | 70, 120 | **<.0001** | 0.623694 | 0.126388 | 0.297782 | 0.949606 |
| 0.5x E2 MB – Bath Soln | 70, 68 | 0.0731 | 0.257599 | 0.143668 | -0.11287 | 0.628071 |
| 0.5x E2 MB – Hank’s | 70, 36 | 0.5167 | 0.112346 | 0.17324 | -0.33438 | 0.559073 |
| 0.5x E2 MB – Sys H_2_O | 70, 138 | 0.4816 | 0.087043 | 0.123663 | -0.23184 | 0.405929 |
| 0.5x E2 MB – Egg H_2_O | 70, 36 | 0.6398 | 0.081096 | 0.17324 | -0.36563 | 0.527823 |
| Ringer’s – Hi Ca^2+^ Ringer’s | 150, 120 | 0.9815 | 0.002398 | 0.103227 | -0.26379 | 0.268587 |
| Ringer’s – Bath Soln | 150, 68 | **0.003** | 0.368493 | 0.123784 | 0.049295 | 0.68769 |
| Ringer’s – Hank’s | 150, 36 | **<.0001** | 0.738438 | 0.157143 | 0.333218 | 1.143657 |
| Ringer’s – Sys H_2_O | 150, 138 | **<.0001** | 0.539049 | 0.099873 | 0.28151 | 0.796588 |
| Ringer’s – Egg H_2_O | 150, 36 | **<.0001** | 0.707188 | 0.157143 | 0.301968 | 1.112407 |
| Hi Ca^2+^ Ringer’s – Bath Soln | 120, 68 | **0.0043** | 0.366095 | 0.128138 | 0.03567 | 0.696521 |
| Hi Ca^2+^ Ringer’s – Hank’s | 120, 36 | **<.0001** | 0.73604 | 0.160596 | 0.321918 | 1.150162 |
| Hi Ca^2+^ Ringer’s – Sys H_2_O | 120, 138 | **<.0001** | 0.536651 | 0.105221 | 0.265321 | 0.807982 |
| Hi Ca^2+^ Ringer’s – Egg H_2_O | 120, 36 | **<.0001** | 0.70479 | 0.160596 | 0.290668 | 1.118912 |
| Bath Soln – Hank’s | 68, 36 | 0.0342 | 0.369945 | 0.174521 | -0.08009 | 0.819976 |
| Bath Soln – Sys H_2_O | 68, 138 | 0.1742 | 0.170556 | 0.125451 | -0.15294 | 0.494054 |
| Bath Soln – Egg H_2_O | 68, 36 | 0.0525 | 0.338695 | 0.174521 | -0.11134 | 0.788726 |
| Hank’s – Sys H_2_O | 36, 138 | 0.2085 | 0.199389 | 0.15846 | -0.20923 | 0.608004 |
| Hank’s – Egg H_2_O | 36, 36 | 0.8756 | 0.03125 | 0.199572 | -0.48338 | 0.54588 |
| Sys H_2_O – Egg H_2_O | 138, 36 | 0.2888 | 0.168139 | 0.15846 | -0.24048 | 0.576754 |
| **C1 Max Angular Velocity** | | | | | | |
| **Comparison** | **(n)** | **p-Value** | **Mean Dif** | **Std Error Dif** | **Lower CL** | **Upper CL** |
| 1x E3 – 1x E3 MB | 357, 292 | **<.0001** | 1.30942 | 0.22511 | 0.72889 | 1.889945 |
| 1x E3 – 1x E2 | 357, 168 | 0.2491 | 0.307774 | 0.266925 | -0.38059 | 0.996133 |
| 1x E3 – 1x E2 MB | 357, 236 | **<.0001** | 1.047349 | 0.239351 | 0.4301 | 1.6646 |
| 1x E3 – 0.5x E2 MB | 357, 69 | **0.0001** | 1.443631 | 0.375183 | 0.47609 | 2.411175 |
| 1x E3 – Ringer’s | 357, 146 | **<.0001** | 1.789357 | 0.280266 | 1.06659 | 2.512123 |
| 1x E3 – Hi Ca^2+^ Ringer’s | 357, 111 | **<.0001** | 2.69831 | 0.310045 | 1.89875 | 3.497871 |
| 1x E3 – Bath Soln | 357, 62 | **<.0001** | 1.753947 | 0.392532 | 0.74167 | 2.766229 |
| 1x E3 – Hank’s | 357, 35 | 0.0459 | 1.009753 | 0.505327 | -0.29341 | 2.312917 |
| 1x E3 – Sys H_2_O | 357, 128 | **0.0079** | 0.781629 | 0.29392 | 0.02365 | 1.539606 |
| 1x E3 – Egg H_2_O | 357, 34 | 0.2815 | 0.551667 | 0.51205 | -0.76884 | 1.872169 |
| 1x E3 MB – 1x E2 | 292, 268 | **0.0003** | 1.001646 | 0.276268 | 0.28919 | 1.714101 |
| 1x E3 MB – 1x E2 MB | 292, 236 | 0.2941 | 0.26207 | 0.249728 | -0.38194 | 0.906082 |
| 1x E3 MB – 0.5x E2 MB | 292, 69 | 0.7253 | 0.134212 | 0.381887 | -0.85062 | 1.119043 |
| 1x E3 MB – Ringer’s | 292, 146 | 0.0972 | 0.479937 | 0.289179 | -0.26581 | 1.225688 |
| 1x E3 MB – Hi Ca^2+^ Ringer’s | 292, 111 | **<.0001** | 1.388891 | 0.318125 | 0.56849 | 2.209287 |
| 1x E3 MB – Bath Soln | 292, 62 | 0.2653 | 0.444527 | 0.398944 | -0.58429 | 1.473345 |
| 1x E3 MB – Hank’s | 292, 35 | **<.0001** | 2.319172 | 0.510324 | 1.00312 | 3.635224 |
| 1x E3 MB – Sys H_2_O | 292, 128 | 0.0811 | 0.527791 | 0.302431 | -0.25213 | 1.307715 |
| 1x E3 MB – Egg H_2_O | 292, 34 | 0.1429 | 0.757753 | 0.516982 | -0.57547 | 2.090974 |
| 1x E2 – 1x E2 MB | 168, 236 | 0.0103 | 0.739576 | 0.28799 | -0.00311 | 1.48226 |
| 1x E2 – 0.5x E2 MB | 168, 69 | **0.0054** | 1.135858 | 0.407936 | 0.08385 | 2.187866 |
| 1x E2 – Ringer’s | 168, 146 | **<.0001** | 1.481583 | 0.322798 | 0.64913 | 2.314033 |
| 1x E2 – Hi Ca^2+^ Ringer’s | 168, 111 | **<.0001** | 2.390537 | 0.348966 | 1.4906 | 3.29047 |
| 1x E2 – Bath Soln | 168, 62 | **0.0007** | 1.446173 | 0.423946 | 0.35288 | 2.539469 |
| 1x E2 – Hank’s | 168, 35 | 0.013 | 1.317526 | 0.530099 | -0.04952 | 2.684574 |
| 1x E2 – Sys H_2_O | 168, 128 | 0.1571 | 0.473855 | 0.334722 | -0.38934 | 1.337054 |
| 1x E2 – Egg H_2_O | 168, 34 | 0.6495 | 0.243893 | 0.536512 | -1.13969 | 1.627478 |
| 1x E2 MB – 0.5x E2 MB | 236, 69 | 0.3103 | 0.396282 | 0.390451 | -0.61064 | 1.403199 |
| 1x E2 MB – Ringer’s | 236, 146 | 0.0136 | 0.742007 | 0.300398 | -0.03267 | 1.51669 |
| 1x E2 MB – Hi Ca^2+^ Ringer’s | 236, 111 | **<.0001** | 1.650961 | 0.328356 | 0.80418 | 2.497743 |
| 1x E2 MB – Bath Soln | 236, 62 | 0.0828 | 0.706597 | 0.407149 | -0.34338 | 1.756576 |
| 1x E2 MB – Hank’s | 236, 35 | **<.0001** | 2.057102 | 0.516764 | 0.72444 | 3.389761 |
| 1x E2 MB – Sys H_2_O | 236, 128 | 0.3963 | 0.26572 | 0.313175 | -0.54191 | 1.073353 |
| 1x E2 MB – Egg H_2_O | 236, 34 | 0.3437 | 0.495683 | 0.52334 | -0.85393 | 1.8453 |
| 0.5x E2 MB – Ringer’s | 69, 146 | 0.4069 | 0.345726 | 0.416789 | -0.72911 | 1.420562 |
| 0.5x E2 MB – Hi Ca^2+^ Ringer’s | 69, 111 | **0.0042** | 1.254679 | 0.437369 | 0.12677 | 2.382589 |
| 0.5x E2 MB – Bath Soln | 69, 62 | 0.5343 | 0.310316 | 0.499244 | -0.97716 | 1.597793 |
| 0.5x E2 MB – Hank’s | 69, 35 | **<.0001** | 2.453384 | 0.592046 | 0.92658 | 3.980185 |
| 0.5x E2 MB – Sys H_2_O | 69, 128 | 0.1205 | 0.662002 | 0.42609 | -0.43682 | 1.760826 |
| 0.5x E2 MB – Egg H_2_O | 69, 34 | 0.1359 | 0.891965 | 0.597795 | -0.64966 | 2.43359 |
| Ringer’s – Hi Ca^2+^ Ringer’s | 146, 111 | 0.0115 | 0.908953 | 0.359274 | -0.01756 | 1.83547 |
| Ringer’s – Bath Soln | 146, 62 | 0.9348 | 0.03541 | 0.432471 | -1.07987 | 1.15069 |
| Ringer’s – Hank’s | 146, 35 | **<.0001** | 2.79911 | 0.536941 | 1.41442 | 4.183802 |
| Ringer’s – Sys H_2_O | 146, 128 | **0.0036** | 1.007728 | 0.345455 | 0.11685 | 1.898606 |
| Ringer’s – Egg H_2_O | 146, 34 | 0.0228 | 1.23769 | 0.543273 | -0.16333 | 2.638711 |
| Hi Ca^2+^ Ringer’s – Bath Soln | 111, 62 | 0.037 | 0.944364 | 0.452338 | -0.22215 | 2.110878 |
| Hi Ca^2+^ Ringer’s – Hank’s | 111, 35 | **<.0001** | 3.708063 | 0.553068 | 2.28178 | 5.134345 |
| Hi Ca^2+^ Ringer’s – Sys H_2_O | 111, 128 | **<.0001** | 1.916681 | 0.370024 | 0.96244 | 2.870919 |
| Hi Ca^2+^ Ringer’s – Egg H_2_O | 111, 34 | **0.0001** | 2.146644 | 0.559217 | 0.7045 | 3.588783 |
| Bath Soln – Hank’s | 62, 35 | **<.0001** | 2.7637 | 0.603189 | 1.20816 | 4.319237 |
| Bath Soln – Sys H_2_O | 62, 128 | 0.0278 | 0.972318 | 0.441442 | -0.1661 | 2.110732 |
| Bath Soln – Egg H_2_O | 62, 34 | 0.0485 | 1.20228 | 0.608833 | -0.36781 | 2.77237 |
| Hank’s – Sys H_2_O | 35, 128 | **0.001** | 1.791382 | 0.544192 | 0.38799 | 3.194774 |
| Hank’s – Egg H_2_O | 35, 34 | 0.0232 | 1.56142 | 0.686987 | -0.21022 | 3.333059 |
| Sys H_2_O – Egg H_2_O | 128, 34 | 0.6762 | 0.229962 | 0.550441 | -1.18954 | 1.649469 |
| **C2 Angle** | | | | | | |
| **Comparison** | **(n)** | **p-Value** | **Mean Dif** | **Std Error Dif** | **Lower CL** | **Upper CL** |
| 1x E3 – 1x E3 MB | 380, 314 | **<.0001** | 5.82105 | 1.164471 | 2.8183 | 8.82384 |
| 1x E3 – 1x E2 | 380, 180 | 0.9576 | 0.07339 | 1.381566 | -3.4892 | 3.636 |
| 1x E3 – 1x E2 MB | 380, 250 | 0.3506 | 1.16105 | 1.243409 | -2.0453 | 4.3674 |
| 1x E3 – 0.5x E2 MB | 380, 70 | 0.0545 | 3.8218 | 1.985962 | -1.2993 | 8.94295 |
| 1x E3 – Ringer’s | 380, 150 | 0.2395 | 1.73228 | 1.472333 | -2.0644 | 5.52895 |
| 1x E3 – Hi Ca^2+^ Ringer’s | 380, 120 | 0.1411 | 2.35395 | 1.598852 | -1.769 | 6.47686 |
| 1x E3 – Bath Soln | 380, 68 | **<.0001** | 10.87012 | 2.010473 | 5.6858 | 16.05448 |
| 1x E3 – Hank’s | 380, 36 | 0.0178 | 6.31784 | 2.66262 | -0.5482 | 13.18386 |
| 1x E3 – Sys H_2_O | 380, 138 | **<.0001** | 7.14996 | 1.517536 | 3.2367 | 11.06319 |
| 1x E3 – Egg H_2_O | 380, 36 | **0.0015** | 8.48772 | 2.66262 | 1.6217 | 15.35374 |
| 1x E3 MB – 1x E2 | 314, 180 | **<.0001** | 5.89444 | 1.427473 | 2.2135 | 9.57543 |
| 1x E3 MB – 1x E2 MB | 314, 250 | **0.0003** | 4.66 | 1.294227 | 1.3226 | 7.99739 |
| 1x E3 MB – 0.5x E2 MB | 314, 70 | **<.0001** | 9.64286 | 2.018168 | 4.4387 | 14.84705 |
| 1x E3 MB – Ringer’s | 314, 150 | **<.0001** | 7.55333 | 1.515494 | 3.6454 | 11.4613 |
| 1x E3 MB – Hi Ca^2+^ Ringer’s | 314, 120 | **<.0001** | 8.175 | 1.638683 | 3.9494 | 12.40063 |
| 1x E3 MB – Bath Soln | 314, 68 | **<.0001** | 16.69118 | 2.042293 | 11.4248 | 21.95758 |
| 1x E3 MB – Hank’s | 314, 36 | **<.0001** | 12.13889 | 2.686727 | 5.2107 | 19.06708 |
| 1x E3 MB – Sys H_2_O | 314, 138 | **<.0001** | 12.97101 | 1.559446 | 8.9497 | 16.99232 |
| 1x E3 MB – Egg H_2_O | 314, 36 | 0.3211 | 2.66667 | 2.686727 | -4.2615 | 9.59486 |
| 1x E2 – 1x E2 MB | 180, 250 | 0.4083 | 1.23444 | 1.492566 | -2.6144 | 5.08328 |
| 1x E2 – 0.5x E2 MB | 180, 70 | 0.0815 | 3.74841 | 2.150752 | -1.7977 | 9.2945 |
| 1x E2 – Ringer’s | 180, 150 | 0.3259 | 1.65889 | 1.688032 | -2.694 | 6.01177 |
| 1x E2 – Hi Ca^2+^ Ringer’s | 180, 120 | 0.2052 | 2.28056 | 1.799448 | -2.3596 | 6.92074 |
| 1x E2 – Bath Soln | 180, 68 | **<.0001** | 10.79673 | 2.173405 | 5.1922 | 16.40123 |
| 1x E2 – Hank’s | 180, 36 | 0.0252 | 6.24444 | 2.787693 | -0.9441 | 13.43299 |
| 1x E2 – Sys H_2_O | 180, 138 | **<.0001** | 7.07657 | 1.727601 | 2.6217 | 11.53149 |
| 1x E2 – Egg H_2_O | 180, 36 | **0.0022** | 8.56111 | 2.787693 | 1.3726 | 15.74966 |
| 1x E2 MB – 0.5x E2 MB | 250, 70 | 0.0159 | 4.98286 | 2.064722 | -0.3414 | 10.3071 |
| 1x E2 MB – Ringer’s | 250, 150 | 0.0667 | 2.89333 | 1.576957 | -1.1731 | 6.95979 |
| 1x E2 MB – Hi Ca^2+^ Ringer’s | 250, 120 | 0.0383 | 3.515 | 1.695687 | -0.8576 | 7.88762 |
| 1x E2 MB – Bath Soln | 250, 68 | **<.0001** | 12.03118 | 2.088309 | 6.6461 | 17.41624 |
| 1x E2 MB – Hank’s | 250, 36 | **0.0061** | 7.47889 | 2.72187 | 0.4601 | 14.4977 |
| 1x E2 MB – Sys H_2_O | 250, 138 | **<.0001** | 8.31101 | 1.619242 | 4.1355 | 12.48651 |
| 1x E2 MB – Egg H_2_O | 250, 36 | **0.0072** | 7.32667 | 2.72187 | 0.3079 | 14.34548 |
| 0.5x E2 MB – Ringer’s | 70, 150 | 0.3446 | 2.08952 | 2.210153 | -3.6097 | 7.78878 |
| 0.5x E2 MB – Hi Ca^2+^ Ringer’s | 70, 120 | 0.5228 | 1.46786 | 2.296375 | -4.4537 | 7.38946 |
| 0.5x E2 MB – Bath Soln | 70, 68 | **0.0068** | 7.04832 | 2.59981 | 0.3443 | 13.75238 |
| 0.5x E2 MB – Hank’s | 70, 36 | 0.4255 | 2.49603 | 3.131542 | -5.5792 | 10.57125 |
| 0.5x E2 MB – Sys H_2_O | 70, 138 | 0.1376 | 3.32816 | 2.240519 | -2.4494 | 9.10572 |
| 0.5x E2 MB – Egg H_2_O | 70, 36 | **<.0001** | 12.30952 | 3.131542 | 4.2343 | 20.38475 |
| Ringer’s – Hi Ca^2+^ Ringer’s | 150, 120 | 0.7396 | 0.62167 | 1.870041 | -4.2006 | 5.44389 |
| Ringer’s – Bath Soln | 150, 68 | **<.0001** | 9.13784 | 2.232203 | 3.3817 | 14.89396 |
| Ringer’s – Hank’s | 150, 36 | 0.1058 | 4.58556 | 2.833774 | -2.7218 | 11.89293 |
| Ringer’s – Sys H_2_O | 150, 138 | **0.0027** | 5.41768 | 1.801012 | 0.7735 | 10.0619 |
| Ringer’s – Egg H_2_O | 150, 36 | **0.0003** | 10.22 | 2.833774 | 2.9126 | 17.52737 |
| Hi Ca^2+^ Ringer’s – Bath Soln | 120, 68 | **0.0002** | 8.51618 | 2.317605 | 2.5398 | 14.49252 |
| Hi Ca^2+^ Ringer’s – Hank’s | 120, 36 | 0.1721 | 3.96389 | 2.901523 | -3.5182 | 11.44597 |
| Hi Ca^2+^ Ringer’s – Sys H_2_O | 120, 138 | 0.0119 | 4.79601 | 1.905835 | -0.1185 | 9.71054 |
| Hi Ca^2+^ Ringer’s – Egg H_2_O | 120, 36 | **0.0002** | 10.84167 | 2.901523 | 3.3596 | 18.32374 |
| Bath Soln – Hank’s | 68, 36 | 0.1482 | 4.55229 | 3.147143 | -3.5632 | 12.66774 |
| Bath Soln – Sys H_2_O | 68, 138 | 0.1003 | 3.72016 | 2.262274 | -2.1135 | 9.55383 |
| Bath Soln – Egg H_2_O | 68, 36 | **<.0001** | 19.35784 | 3.147143 | 11.2424 | 27.4733 |
| Hank’s – Sys H_2_O | 36, 138 | 0.7709 | 0.83213 | 2.857521 | -6.5365 | 8.20074 |
| Hank’s – Egg H_2_O | 36, 36 | **<.0001** | 14.80556 | 3.598896 | 5.5252 | 24.08593 |
| Sys H_2_O – Egg H_2_O | 138, 36 | **<.0001** | 15.63768 | 2.857521 | 8.2691 | 23.00629 |
| **Normalized Total Distance (spontaneous)** | | | | | | |
| **Comparison** | **(n)** | **p-Value** | **Mean Dif** | **Std Error Dif** | **Lower CL** | **Upper CL** |
| 1x E3 – 1x E3 MB | 398, 382 | 0.6513 | 0.021736 | 0.048087 | -0.10226 | 0.145726 |
| 1x E3 – 1x E2 | 398, 193 | **<.0001** | 0.455295 | 0.058849 | 0.30356 | 0.607034 |
| 1x E3 – 1x E2 MB | 398, 265 | **0.0074** | 0.14253 | 0.053193 | 0.00537 | 0.279687 |
| 1x E3 – 0.5x E2 MB | 398, 72 | **<.0001** | 0.593193 | 0.085922 | 0.37165 | 0.81474 |
| 1x E3 – Ringer’s | 398, 154 | **<.0001** | 0.404966 | 0.06367 | 0.2408 | 0.569136 |
| 1x E3 – Hi Ca^2+^ Ringer’s | 398, 140 | **<.0001** | 0.606402 | 0.065925 | 0.43642 | 0.776386 |
| 1x E3 – Bath Soln | 398, 69 | **<.0001** | 0.645152 | 0.08749 | 0.41956 | 0.87074 |
| 1x E3 – Hank’s | 398, 36 | **<.0001** | 0.742146 | 0.116766 | 0.44107 | 1.043222 |
| 1x E3 – Sys H_2_O | 398, 141 | **<.0001** | 0.392857 | 0.065752 | 0.22332 | 0.562396 |
| 1x E3 – Egg H_2_O | 398, 36 | **0.0006** | 0.400777 | 0.116766 | 0.0997 | 0.701853 |
| 1x E3 MB – 1x E2 | 382, 193 | **<.0001** | 0.47703 | 0.059276 | 0.32419 | 0.629871 |
| 1x E3 MB – 1x E2 MB | 382, 265 | **0.0022** | 0.164266 | 0.053666 | 0.02589 | 0.30264 |
| 1x E3 MB – 0.5x E2 MB | 382, 72 | **<.0001** | 0.571458 | 0.086216 | 0.34916 | 0.793761 |
| 1x E3 MB – Ringer’s | 382, 154 | **<.0001** | 0.383231 | 0.064065 | 0.21804 | 0.548419 |
| 1x E3 MB – Hi Ca^2+^ Ringer’s | 382, 140 | **<.0001** | 0.584666 | 0.066307 | 0.4137 | 0.755635 |
| 1x E3 MB – Bath Soln | 382, 69 | **<.0001** | 0.623416 | 0.087778 | 0.39709 | 0.849747 |
| 1x E3 MB – Hank’s | 382, 36 | **<.0001** | 0.763882 | 0.116982 | 0.46225 | 1.065515 |
| 1x E3 MB – Sys H_2_O | 382, 141 | **<.0001** | 0.371122 | 0.066135 | 0.2006 | 0.541647 |
| 1x E3 MB – Egg H_2_O | 382, 36 | **0.0003** | 0.422513 | 0.116982 | 0.12088 | 0.724146 |
| 1x E2 – 1x E2 MB | 193, 265 | **<.0001** | 0.312765 | 0.063489 | 0.14906 | 0.476467 |
| 1x E2 – 0.5x E2 MB | 193, 72 | **<.0001** | 1.048488 | 0.09265 | 0.8096 | 1.287381 |
| 1x E2 – Ringer’s | 193, 154 | **<.0001** | 0.860261 | 0.072492 | 0.67334 | 1.047178 |
| 1x E2 – Hi Ca^2+^ Ringer’s | 193, 140 | **<.0001** | 1.061696 | 0.074481 | 0.86965 | 1.253742 |
| 1x E2 – Bath Soln | 193, 69 | **<.0001** | 1.100446 | 0.094105 | 0.8578 | 1.343092 |
| 1x E2 – Hank’s | 193, 36 | 0.0186 | 0.286851 | 0.121802 | -0.02721 | 0.600911 |
| 1x E2 – Sys H_2_O | 193, 141 | **<.0001** | 0.848152 | 0.074328 | 0.6565 | 1.039802 |
| 1x E2 – Egg H_2_O | 193, 36 | 0.6545 | 0.054518 | 0.121802 | -0.25954 | 0.368577 |
| 1x E2 MB – 0.5x E2 MB | 265, 72 | **<.0001** | 0.735724 | 0.089164 | 0.50582 | 0.96563 |
| 1x E2 MB – Ringer’s | 265, 154 | **<.0001** | 0.547496 | 0.067981 | 0.37221 | 0.722783 |
| 1x E2 MB – Hi Ca^2+^ Ringer’s | 265, 140 | **<.0001** | 0.748932 | 0.070098 | 0.56819 | 0.929676 |
| 1x E2 MB – Bath Soln | 265, 69 | **<.0001** | 0.787682 | 0.090676 | 0.55388 | 1.021485 |
| 1x E2 MB – Hank’s | 265, 36 | **<.0001** | 0.599616 | 0.119172 | 0.29234 | 0.906896 |
| 1x E2 MB – Sys H_2_O | 265, 141 | **<.0001** | 0.535388 | 0.069935 | 0.35506 | 0.715712 |
| 1x E2 MB – Egg H_2_O | 265, 36 | 0.0304 | 0.258247 | 0.119172 | -0.04903 | 0.565527 |
| 0.5x E2 MB – Ringer’s | 72, 154 | 0.0495 | 0.188227 | 0.095784 | -0.05875 | 0.435202 |
| 0.5x E2 MB – Hi Ca^2+^ Ringer’s | 72, 140 | 0.892 | 0.013208 | 0.097298 | -0.23767 | 0.264086 |
| 0.5x E2 MB – Bath Soln | 72, 69 | 0.6458 | 0.051958 | 0.113027 | -0.23948 | 0.343394 |
| 0.5x E2 MB – Hank’s | 72, 36 | **<.0001** | 1.33534 | 0.136949 | 0.98222 | 1.688457 |
| 0.5x E2 MB – Sys H_2_O | 72, 141 | 0.0394 | 0.200336 | 0.097181 | -0.05024 | 0.450912 |
| 0.5x E2 MB – Egg H_2_O | 72, 36 | **<.0001** | 0.993971 | 0.136949 | 0.64085 | 1.347088 |
| Ringer’s – Hi Ca^2+^ Ringer’s | 154, 140 | 0.0102 | 0.201435 | 0.078346 | -0.00058 | 0.403446 |
| Ringer’s – Bath Soln | 154, 69 | 0.0136 | 0.240186 | 0.097192 | -0.01042 | 0.490792 |
| Ringer’s – Hank’s | 154, 36 | **<.0001** | 1.147112 | 0.124203 | 0.82686 | 1.467363 |
| Ringer’s – Sys H_2_O | 154, 141 | 0.877 | 0.012109 | 0.0782 | -0.18953 | 0.213744 |
| Ringer’s – Egg H_2_O | 154, 36 | **<.0001** | 0.805743 | 0.124203 | 0.48549 | 1.125994 |
| Hi Ca^2+^ Ringer’s – Bath Soln | 140, 69 | 0.6946 | 0.03875 | 0.098685 | -0.2157 | 0.293204 |
| Hi Ca^2+^ Ringer’s – Hank’s | 140, 36 | **<.0001** | 1.348548 | 0.125374 | 1.02528 | 1.671818 |
| Hi Ca^2+^ Ringer’s – Sys H_2_O | 140, 141 | **0.0077** | 0.213544 | 0.080047 | 0.00715 | 0.419942 |
| Hi Ca^2+^ Ringer’s – Egg H_2_O | 140, 36 | **<.0001** | 1.007179 | 0.125374 | 0.68391 | 1.330449 |
| Bath Soln – Hank’s | 69, 36 | **<.0001** | 1.387298 | 0.137938 | 1.03163 | 1.742965 |
| Bath Soln – Sys H_2_O | 69, 141 | 0.0106 | 0.252294 | 0.098569 | -0.00186 | 0.50645 |
| Bath Soln – Egg H_2_O | 69, 36 | **<.0001** | 1.045929 | 0.137938 | 0.69026 | 1.401596 |
| Hank’s – Sys H_2_O | 36, 141 | **<.0001** | 1.135003 | 0.125283 | 0.81197 | 1.458039 |
| Hank’s – Egg H_2_O | 36, 36 | 0.031 | 0.341369 | 0.158135 | -0.06638 | 0.749114 |
| Sys H_2_O – Egg H_2_O | 141, 36 | **<.0001** | 0.793635 | 0.125283 | 0.4706 | 1.11667 |
| **Swim Frequency** | | | | | | |
| **Comparison** | **(n)** | **p-Value** | **Mean Dif** | **Std Error Dif** | **Lower CL** | **Upper CL** |
| 1x E3 – 1x E3 MB | 319, 373 | **0.0003** | 2.26503 | 0.626435 | 0.64937 | 3.8807 |
| 1x E3 – 1x E2 | 319, 216 | 0.0516 | 1.41031 | 0.723814 | -0.45651 | 3.27713 |
| 1x E3 – 1x E2 MB | 319, 250 | 0.9769 | 0.02009 | 0.693846 | -1.76944 | 1.80962 |
| 1x E3 – 0.5x E2 MB | 319, 28 | **<.0001** | 8.01191 | 1.619059 | 3.83612 | 12.1877 |
| 1x E3 – Ringer’s | 319, 93 | 0.3413 | 0.92142 | 0.96802 | -1.57524 | 3.41809 |
| 1x E3 – Hi Ca^2+^ Ringer’s | 319, 87 | 0.0175 | 2.36395 | 0.993529 | -0.19851 | 4.92641 |
| 1x E3 – Bath Soln | 319, 17 | 0.0119 | 5.14721 | 2.044665 | -0.12629 | 10.4207 |
| 1x E3 – Hank’s | 319, 30 | 0.0262 | 3.49142 | 1.568661 | -0.55439 | 7.53723 |
| 1x E3 – Sys H_2_O | 319, 46 | **<.0001** | 6.98583 | 1.295521 | 3.64449 | 10.32716 |
| 1x E3 – Egg H_2_O | 319, 36 | **<.0001** | 5.83809 | 1.444242 | 2.11318 | 9.563 |
| 1x E3 MB – 1x E2 | 373, 216 | 0.2238 | 0.85472 | 0.702342 | -0.95672 | 2.66616 |
| 1x E3 MB – 1x E2 MB | 373, 250 | **0.0008** | 2.24494 | 0.671417 | 0.51326 | 3.97663 |
| 1x E3 MB – 0.5x E2 MB | 319, 28 | **<.0001** | 10.27694 | 1.609575 | 6.12561 | 14.42827 |
| 1x E3 MB – Ringer’s | 319, 93 | 0.1584 | 1.34361 | 0.952071 | -1.11192 | 3.79914 |
| 1x E3 MB – Hi Ca^2+^ Ringer’s | 319, 87 | 0.9194 | 0.09892 | 0.977996 | -2.42348 | 2.62131 |
| 1x E3 MB – Bath Soln | 319, 17 | **0.0003** | 7.41224 | 2.037163 | 2.1581 | 12.66638 |
| 1x E3 MB – Hank’s | 319, 30 | 0.4316 | 1.22639 | 1.55887 | -2.79416 | 5.24694 |
| 1x E3 MB – Sys H_2_O | 319, 46 | **<.0001** | 9.25086 | 1.283648 | 5.94014 | 12.56157 |
| 1x E3 MB – Egg H_2_O | 319, 36 | 0.0128 | 3.57306 | 1.433601 | -0.12441 | 7.27052 |
| 1x E2 – 1x E2 MB | 216, 250 | 0.0687 | 1.39022 | 0.763077 | -0.57787 | 3.35831 |
| 1x E2 – 0.5x E2 MB | 216, 28 | **<.0001** | 9.42222 | 1.649914 | 5.16685 | 13.67759 |
| 1x E2 – Ringer’s | 216, 93 | 0.6314 | 0.48889 | 1.018787 | -2.13871 | 3.11649 |
| 1x E2 – Hi Ca^2+^ Ringer’s | 216, 87 | 0.3607 | 0.95364 | 1.043055 | -1.73655 | 3.64383 |
| 1x E2 – Bath Soln | 216, 17 | **0.0016** | 6.55752 | 2.069184 | 1.22079 | 11.89424 |
| 1x E2 – Hank’s | 216, 30 | 0.1937 | 2.08111 | 1.600488 | -2.04678 | 6.209 |
| 1x E2 – Sys H_2_O | 216, 46 | **<.0001** | 8.39614 | 1.333882 | 4.95586 | 11.83641 |
| 1x E2 – Egg H_2_O | 216, 36 | **0.0028** | 4.42778 | 1.478749 | 0.61387 | 8.24169 |
| 1x E2 MB – 0.5x E2 MB | 250, 28 | **<.0001** | 8.032 | 1.636989 | 3.80997 | 12.25403 |
| 1x E2 MB – Ringer’s | 250, 93 | 0.3665 | 0.90133 | 0.997718 | -1.67193 | 3.4746 |
| 1x E2 MB – Hi Ca^2+^ Ringer’s | 250, 87 | 0.022 | 2.34386 | 1.022487 | -0.29328 | 4.98101 |
| 1x E2 MB – Bath Soln | 250, 17 | 0.0122 | 5.16729 | 2.058892 | -0.14289 | 10.47748 |
| 1x E2 MB – Hank’s | 250, 30 | 0.0289 | 3.47133 | 1.58716 | -0.62219 | 7.56485 |
| 1x E2 MB – Sys H_2_O | 250, 46 | **<.0001** | 7.00591 | 1.31786 | 3.60696 | 10.40487 |
| 1x E2 MB – Egg H_2_O | 250, 36 | **<.0001** | 5.818 | 1.464314 | 2.04132 | 9.59468 |
| 0.5x E2 MB – Ringer’s | 28, 93 | **<.0001** | 8.93333 | 1.770698 | 4.36644 | 13.50022 |
| 0.5x E2 MB – Hi Ca^2+^ Ringer’s | 28, 87 | **<.0001** | 10.37586 | 1.784772 | 5.77267 | 14.97905 |
| 0.5x E2 MB – Bath Soln | 28, 17 | 0.2569 | 2.86471 | 2.525662 | -3.64935 | 9.37876 |
| 0.5x E2 MB – Hank’s | 28, 30 | **<.0001** | 11.50333 | 2.158474 | 5.93631 | 17.07035 |
| 0.5x E2 MB – Sys H_2_O | 28, 46 | 0.6023 | 1.02609 | 1.96893 | -4.05207 | 6.10425 |
| 0.5x E2 MB – Egg H_2_O | 28, 36 | **<.0001** | 13.85 | 2.069818 | 8.51164 | 19.18836 |
| Ringer’s – Hi Ca^2+^ Ringer’s | 93, 87 | 0.2392 | 1.44253 | 1.225202 | -1.71745 | 4.6025 |
| Ringer’s – Bath Soln | 93, 17 | **0.0052** | 6.06863 | 2.16672 | 0.48034 | 11.65692 |
| Ringer’s – Hank’s | 93, 30 | 0.1364 | 2.57 | 1.724737 | -1.87835 | 7.01835 |
| Ringer’s – Sys H_2_O | 93, 46 | **<.0001** | 7.90725 | 1.480674 | 4.08837 | 11.72612 |
| Ringer’s – Egg H_2_O | 93, 36 | **0.0023** | 4.91667 | 1.612407 | 0.75803 | 9.0753 |
| Hi Ca^2+^ Ringer’s – Bath Soln | 87, 17 | **0.0006** | 7.51116 | 2.178236 | 1.89317 | 13.12915 |
| Hi Ca^2+^ Ringer’s – Hank’s | 87, 30 | 0.5169 | 1.12747 | 1.739182 | -3.35813 | 5.61308 |
| Hi Ca^2+^ Ringer’s – Sys H_2_O | 87, 46 | **<.0001** | 9.34978 | 1.497475 | 5.48757 | 13.21198 |
| Hi Ca^2+^ Ringer’s – Egg H_2_O | 87, 36 | 0.033 | 3.47414 | 1.627849 | -0.72432 | 7.6726 |
| Bath Soln – Hank’s | 17, 30 | **0.0005** | 8.63863 | 2.493654 | 2.20713 | 15.07013 |
| Bath Soln – Sys H_2_O | 17, 46 | 0.4305 | 1.83862 | 2.331521 | -4.17471 | 7.85195 |
| Bath Soln – Egg H_2_O | 17, 36 | **<.0001** | 10.98529 | 2.417323 | 4.75066 | 17.21992 |
| Hank’s – Sys H_2_O | 30, 46 | **<.0001** | 10.47725 | 1.927701 | 5.50542 | 15.44907 |
| Hank’s – Egg H_2_O | 30, 36 | 0.248 | 2.34667 | 2.030638 | -2.89065 | 7.58398 |
| Sys H_2_O – Egg H_2_O | 46, 36 | **<.0001** | 12.82391 | 1.827886 | 8.10953 | 17.5383 |
| **Turn Frequency** | | | | | | |
| **Comparison** | **(n)** | **p-Value** | **Mean Dif** | **Std Error Dif** | **Lower CL** | **Upper CL** |
| 1x E3 – 1x E3 MB | 319, 373 | **<.0001** | 5.32425 | 0.901239 | 2.99982 | 7.64867 |
| 1x E3 – 1x E2 | 319, 216 | 0.2349 | 1.23739 | 1.041337 | -1.44837 | 3.92315 |
| 1x E3 – 1x E2 MB | 319, 250 | 0.0363 | 2.09144 | 0.998223 | -0.48312 | 4.666 |
| 1x E3 – 0.5x E2 MB | 319, 28 | 0.0632 | 4.33018 | 2.329309 | -1.67744 | 10.33781 |
| 1x E3 – Ringer’s | 319, 93 | **<.0001** | 11.31713 | 1.392671 | 7.72522 | 14.90903 |
| 1x E3 – Hi Ca^2+^ Ringer’s | 319, 87 | **<.0001** | 12.39028 | 1.42937 | 8.70373 | 16.07684 |
| 1x E3 – Bath Soln | 319, 17 | **<.0001** | 14.31422 | 2.94162 | 6.72735 | 21.90108 |
| 1x E3 – Hank’s | 319, 30 | 0.6837 | 0.91971 | 2.256802 | -4.90091 | 6.74033 |
| 1x E3 – Sys H_2_O | 319, 46 | 0.031 | 4.02521 | 1.863841 | -0.7819 | 8.83233 |
| 1x E3 – Egg H_2_O | 319, 36 | **<.0001** | 8.84526 | 2.077803 | 3.48631 | 14.20422 |
| 1x E3 MB – 1x E2 | 373, 216 | **<.0001** | 4.08685 | 1.010446 | 1.48077 | 6.69294 |
| 1x E3 MB – 1x E2 MB | 373, 250 | **0.0008** | 3.23281 | 0.965954 | 0.74147 | 5.72414 |
| 1x E3 MB – 0.5x E2 MB | 319, 28 | 0.6678 | 0.99406 | 2.315663 | -4.97837 | 6.9665 |
| 1x E3 MB – Ringer’s | 319, 93 | **<.0001** | 5.99288 | 1.369726 | 2.46016 | 9.5256 |
| 1x E3 MB – Hi Ca^2+^ Ringer’s | 319, 87 | **<.0001** | 7.06603 | 1.407024 | 3.43711 | 10.69496 |
| 1x E3 MB – Bath Soln | 319, 17 | **0.0022** | 8.98997 | 2.930827 | 1.43094 | 16.549 |
| 1x E3 MB – Hank’s | 319, 30 | 0.0497 | 4.40454 | 2.242715 | -1.37975 | 10.18883 |
| 1x E3 MB – Sys H_2_O | 319, 46 | 0.4819 | 1.29903 | 1.84676 | -3.46403 | 6.0621 |
| 1x E3 MB – Egg H_2_O | 319, 36 | 0.088 | 3.52102 | 2.062494 | -1.79846 | 8.84049 |
| 1x E2 – 1x E2 MB | 216, 250 | 0.4367 | 0.85405 | 1.097824 | -1.9774 | 3.6855 |
| 1x E2 – 0.5x E2 MB | 216, 28 | 0.1928 | 3.09279 | 2.373699 | -3.02933 | 9.21491 |
| 1x E2 – Ringer’s | 216, 93 | **<.0001** | 10.07973 | 1.465708 | 6.29946 | 13.86001 |
| 1x E2 – Hi Ca^2+^ Ringer’s | 216, 87 | **<.0001** | 11.15289 | 1.500623 | 7.28256 | 15.02321 |
| 1x E2 – Bath Soln | 216, 17 | **<.0001** | 13.07682 | 2.976894 | 5.39898 | 20.75467 |
| 1x E2 – Hank’s | 216, 30 | 0.8903 | 0.31769 | 2.302591 | -5.62103 | 6.2564 |
| 1x E2 – Sys H_2_O | 216, 46 | 0.1465 | 2.78782 | 1.919029 | -2.16163 | 7.73728 |
| 1x E2 – Egg H_2_O | 216, 36 | **0.0004** | 7.60787 | 2.127447 | 2.12087 | 13.09487 |
| 1x E2 MB – 0.5x E2 MB | 250, 28 | 0.342 | 2.23874 | 2.355104 | -3.83541 | 8.3129 |
| 1x E2 MB – Ringer’s | 250, 93 | **<.0001** | 9.22569 | 1.435398 | 5.52359 | 12.92779 |
| 1x E2 MB – Hi Ca^2+^ Ringer’s | 250, 87 | **<.0001** | 10.29884 | 1.471032 | 6.50484 | 14.09285 |
| 1x E2 MB – Bath Soln | 250, 17 | **<.0001** | 12.22278 | 2.962087 | 4.58312 | 19.86243 |
| 1x E2 MB – Hank’s | 250, 30 | 0.6079 | 1.17173 | 2.283416 | -4.71753 | 7.061 |
| 1x E2 MB – Sys H_2_O | 250, 46 | 0.3079 | 1.93377 | 1.895979 | -2.95623 | 6.82378 |
| 1x E2 MB – Egg H_2_O | 250, 36 | **0.0014** | 6.75382 | 2.106679 | 1.32039 | 12.18726 |
| 0.5x E2 MB – Ringer’s | 28, 93 | **0.0062** | 6.98694 | 2.547469 | 0.41665 | 13.55724 |
| 0.5x E2 MB – Hi Ca^2+^ Ringer’s | 28, 87 | **0.0017** | 8.0601 | 2.567716 | 1.43758 | 14.68261 |
| 0.5x E2 MB – Bath Soln | 28, 17 | **0.0061** | 9.98403 | 3.63362 | 0.6124 | 19.35567 |
| 0.5x E2 MB – Hank’s | 28, 30 | 0.2723 | 3.41048 | 3.105353 | -4.59868 | 11.41964 |
| 0.5x E2 MB – Sys H_2_O | 28, 46 | 0.9143 | 0.30497 | 2.832661 | -7.00088 | 7.61082 |
| 0.5x E2 MB – Egg H_2_O | 28, 36 | 0.1297 | 4.51508 | 2.977806 | -3.16512 | 12.19528 |
| Ringer’s – Hi Ca^2+^ Ringer’s | 93, 87 | 0.5427 | 1.07316 | 1.762673 | -3.47304 | 5.61935 |
| Ringer’s – Bath Soln | 93, 17 | 0.3365 | 2.99709 | 3.117217 | -5.04267 | 11.03685 |
| Ringer’s – Hank’s | 93, 30 | **<.0001** | 10.39742 | 2.481345 | 3.99767 | 16.79717 |
| Ringer’s – Sys H_2_O | 93, 46 | **0.0006** | 7.29191 | 2.130216 | 1.79777 | 12.78605 |
| Ringer’s – Egg H_2_O | 93, 36 | 0.2868 | 2.47186 | 2.319737 | -3.51108 | 8.45481 |
| Hi Ca^2+^ Ringer’s – Bath Soln | 87, 17 | 0.5394 | 1.92394 | 3.133786 | -6.15856 | 10.00643 |
| Hi Ca^2+^ Ringer’s – Hank’s | 87, 30 | **<.0001** | 11.47057 | 2.502127 | 5.01722 | 17.92393 |
| Hi Ca^2+^ Ringer’s – Sys H_2_O | 87, 46 | **0.0001** | 8.36507 | 2.154388 | 2.80858 | 13.92155 |
| Hi Ca^2+^ Ringer’s – Egg H_2_O | 87, 36 | 0.1303 | 3.54502 | 2.341954 | -2.49522 | 9.58526 |
| Bath Soln – Hank’s | 17, 30 | **0.0002** | 13.39451 | 3.587571 | 4.14164 | 22.64738 |
| Bath Soln – Sys H_2_O | 17, 46 | **0.0022** | 10.289 | 3.354313 | 1.63774 | 18.94027 |
| Bath Soln – Egg H_2_O | 17, 36 | 0.116 | 5.46895 | 3.477754 | -3.50068 | 14.43859 |
| Hank’s – Sys H_2_O | 30, 46 | 0.263 | 3.10551 | 2.773345 | -4.04736 | 10.25837 |
| Hank’s – Egg H_2_O | 30, 36 | **0.0067** | 7.92556 | 2.921439 | 0.39074 | 15.46037 |
| Sys H_2_O – Egg H_2_O | 46, 36 | 0.067 | 4.82005 | 2.629744 | -1.96245 | 11.60254 |

**Supplemental Table 3. Multiple comparisons between media types**. Multiple comparisons and their reported values for the acoustic startle response (normalized SLC and LLC Index), pre-pulse inhibition, short-term habituation (habituation half-life), SLC kinematics (latency, C1 angle, C1 curvature, C1 distance, C1 max angular velocity, and C2 angle), and general locomotor behaviors (normalized total distance, swim and turn frequency) with statistically significant p-values in bold (α=0.01, ANOVA with student’s t each pair test for multiple comparisons).

| **Normalized SLC Index (by media)** | | | | | | |
| --- | --- | --- | --- | --- | --- | --- |
| **Comparison** | **(n)** | **p-Value** | **Mean Dif** | **Std Error Dif** | **Lower CL** | **Upper CL** |
| **1x E3** | | | | | | |
| *chd7^+/+^* – *chd7^ncu101/+^* | 18, 25 | 0.1281 | 0.150641 | 0.097499 | -0.04475 | 0.346034 |
| *chd7^+/+^* – *chd7^ncu101/ncu101^* | 18, 15 | 0.2328 | 0.133035 | 0.110268 | -0.08795 | 0.354017 |
| *chd7^ncu101/+^* – *chd7^ncu101/ncu101^* | 25, 15 | 0.8649 | 0.017606 | 0.103012 | -0.18884 | 0.224047 |
| **1x E3 MB** | | | | | | |
| *chd7^+/+^* – *chd7^ncu101/+^* | 18, 38 | 0.8215 | 0.019854 | 0.087656 | -0.15511 | 0.194817 |
| *chd7^+/+^* – *chd7^ncu101/ncu101^* | 18, 14 | 0.2309 | 0.131972 | 0.109167 | -0.08593 | 0.34987 |
| *chd7^ncu101/+^* – *chd7^ncu101/ncu101^* | 38, 14 | 0.2459 | 0.112118 | 0.095777 | -0.07905 | 0.30329 |
| **1x E2** | | | | | | |
| *chd7^+/+^* – *chd7^ncu101/+^* | 20, 36 | 0.9668 | 0.0030369 | 0.0727820 | -0.142362 | 0.1484356 |
| *chd7^+/+^* – *chd7^ncu101/ncu101^* | 20, 11 | 0.1228 | 0.1531875 | 0.0979638 | -0.042518 | 0.3488927 |
| *chd7^ncu101/+^* – *chd7^ncu101/ncu101^* | 36, 11 | 0.0871 | 0.1562244 | 0.0899078 | -0.023387 | 0.3358359 |
| **1x E2 MB** | | | | | | |
| *chd7^+/+^* – *chd7^ncu101/+^* | 13, 33 | 0.7226 | 0.035262 | 0.098862 | -0.16243 | 0.23295 |
| *chd7^+/+^* – *chd7^ncu101/ncu101^* | 13, 18 | 0.3169 | 0.110891 | 0.109889 | -0.10885 | 0.330627 |
| *chd7^ncu101/+^* – *chd7^ncu101/ncu101^* | 33, 18 | 0.1037 | 0.146154 | 0.088465 | -0.03074 | 0.323051 |
| **Hank’s** | | | | | | |
| *chd7^+/+^* – *chd7^ncu101/+^* | 18, 33 | 0.2155 | 0.101761 | 0.081352 | -0.06071 | 0.264232 |
| *chd7^+/+^* – *chd7^ncu101/ncu101^* | 18, 17 | 0.1153 | 0.149874 | 0.093897 | -0.03765 | 0.337399 |
| *chd7^ncu101/+^* – *chd7^ncu101/ncu101^* | 33, 17 | **0.0034** | 0.251635 | 0.082886 | 0.086101 | 0.41717 |
| **Normalized SLC Index (by genotype)** | | | | | | |
| **Comparison** | **(n)** | **p-Value** | **Mean Dif** | **Std Error Dif** | **Lower CL** | **Upper CL** |
| ***chd7^+/+^*** | | | | | | |
| 1x E3 – 1x E3 MB | 18, 18 | **0.0399** | 0.198976 | 0.095282 | 0.00943 | 0.388522 |
| 1x E3 – 1x E2 MB | 18, 13 | **0.0453** | 0.211491 | 0.104041 | 0.00452 | 0.418461 |
| 1x E3 – 1x E2 | 18, 20 | 0.4812 | 0.065719 | 0.092869 | -0.11903 | 0.250465 |
| 1x E3 – Hank’s | 18, 18 | **0.0057** | 0.270317 | 0.095282 | 0.080771 | 0.459862 |
| 1x E2 – 1x E3 MB | 20, 18 | 0.1551 | 0.133257 | 0.092869 | -0.05149 | 0.318003 |
| 1x E2 – 1x E2 MB | 20, 13 | 0.1561 | 0.145771 | 0.101836 | -0.05681 | 0.348356 |
| 1x E2 – Hank’s | 20, 18 | **0.0304** | 0.204598 | 0.092869 | 0.019851 | 0.389344 |
| 1x E3 MB – 1x E2 MB | 18, 13 | 0.9046 | 0.012515 | 0.104041 | -0.19446 | 0.219485 |
| 1x E3 MB – Hank’s | 18, 18 | 0.4562 | 0.071341 | 0.095282 | -0.11821 | 0.260886 |
| 1x E2 MB – Hank’s | 13, 18 | 0.5733 | 0.058826 | 0.104041 | -0.14814 | 0.265796 |
| ***chd7^ncu101/+^*** | | | | | | |
| 1x E3 – 1x E3 MB | 25, 38 | 0.3765 | 0.068189 | 0.07689 | -0.08366 | 0.22004 |
| 1x E3 – 1x E2 MB | 25, 33 | 0.747 | 0.025587 | 0.079168 | -0.13076 | 0.181936 |
| 1x E3 – 1x E2 | 25, 36 | 0.2595 | 0.087959 | 0.077733 | -0.06556 | 0.241474 |
| 1x E3 – Hank’s | 25, 33 | 0.8213 | 0.017914 | 0.079168 | -0.13843 | 0.174263 |
| 1x E2 – 1x E3 MB | 36, 38 | **0.0259** | 0.156148 | 0.069444 | 0.019003 | 0.293293 |
| 1x E2 – 1x E2 MB | 36, 33 | 0.1166 | 0.113546 | 0.071958 | -0.02856 | 0.255655 |
| 1x E2 – Hank’s | 36, 33 | 0.1432 | 0.105873 | 0.071958 | -0.03624 | 0.247983 |
| 1x E3 MB – 1x E2 MB | 38, 33 | 0.5496 | 0.042602 | 0.071046 | -0.09771 | 0.182912 |
| 1x E3 MB – Hank’s | 38, 33 | 0.4802 | 0.050275 | 0.071046 | -0.09004 | 0.190584 |
| 1x E2 MB – Hank’s | 33, 33 | 0.917 | 0.007673 | 0.073505 | -0.13749 | 0.152839 |
| ***chd7^ncu101/ncu101^*** | | | | | | |
| 1x E3 – 1x E3 MB | 15, 14 | **0.0668** | 0.197913 | 0.106273 | -0.01404 | 0.409867 |
| 1x E3 – 1x E2 MB | 15, 18 | **0.0624** | 0.189347 | 0.099979 | -0.01005 | 0.388747 |
| 1x E3 – 1x E2 | 15, 11 | 0.4519 | 0.085872 | 0.113521 | -0.14054 | 0.312282 |
| 1x E3 – Hank’s | 15, 17 | **0.006** | 0.287156 | 0.101306 | 0.085107 | 0.489205 |
| 1x E2 – 1x E3 MB | 11, 14 | 0.3342 | 0.112042 | 0.115224 | -0.11776 | 0.341848 |
| 1x E2 – 1x E2 MB | 11, 18 | 0.3477 | 0.103475 | 0.109446 | -0.11481 | 0.321757 |
| 1x E2 – Hank’s | 11, 17 | 0.0732 | 0.201284 | 0.11066 | -0.01942 | 0.421988 |
| 1x E3 MB – 1x E2 MB | 14, 18 | 0.9332 | 0.008567 | 0.101908 | -0.19468 | 0.211815 |
| 1x E3 MB – Hank’s | 14, 17 | 0.3902 | 0.089243 | 0.103211 | -0.1166 | 0.295089 |
| 1x E2 MB – Hank’s | 18, 17 | 0.3154 | 0.097809 | 0.096717 | -0.09509 | 0.290706 |
| **Normalized LLC Index (by media)** | | | | | | |
| **Comparison** | **(n)** | **p-Value** | **Mean Dif** | **Std Error Dif** | **Lower CL** | **Upper CL** |
| **1x E3** | | | | | | |
| *chd7^+/+^* – *chd7^ncu101/+^* | 18, 25 | 0.061601 | 0.216364 | -0.372 | 0.495205 | 0.7769 |
| *chd7^+/+^* – *chd7^ncu101/ncu101^* | 18, 15 | 0.229351 | 0.244699 | -0.26104 | 0.719738 | 0.3527 |
| *chd7^ncu101/+^* – *chd7^ncu101/ncu101^* | 25, 15 | 0.290952 | 0.228598 | -0.16717 | 0.749072 | 0.2085 |
| **1x E3 MB** | | | | | | |
| *chd7^+/+^* – *chd7^ncu101/+^* | 18, 38 | **0.016362** | 0.226689 | -0.43611 | 0.468836 | 0.9427 |
| *chd7^+/+^* – *chd7^ncu101/ncu101^* | 18, 14 | 0.649858 | 0.282319 | 0.086347 | 1.213369 | 0.0245 |
| *chd7^ncu101/+^* – *chd7^ncu101/ncu101^* | 38, 14 | 0.66622 | 0.247692 | 0.171826 | 1.160615 | 0.009 |
| **1x E2** | | | | | | |
| *chd7^+/+^* – *chd7^ncu101/+^* | 20, 36 | 0.0873 | 0.326871 | 0.188269 | -0.04924 | 0.70298 |
| *chd7^+/+^* – *chd7^ncu101/ncu101^* | 20, 11 | 0.0932 | 0.431854 | 0.253408 | -0.07439 | 0.938094 |
| *chd7^ncu101/+^* – *chd7^ncu101/ncu101^* | 36, 11 | **0.0018** | 0.758724 | 0.232569 | 0.294114 | 1.223334 |
| **1x E2 MB** | | | | | | |
| *chd7^+/+^* – *chd7^ncu101/+^* | 13, 33 | 0.09686 | 0.270983 | -0.445 | 0.638724 | 0.722 |
| *chd7^+/+^* – *chd7^ncu101/ncu101^* | 13, 18 | 0.949868 | 0.301207 | 0.347568 | 1.552168 | 0.0025 |
| *chd7^ncu101/+^* – *chd7^ncu101/ncu101^* | 33, 18 | 0.853008 | 0.242484 | 0.368131 | 1.337885 | 0.0008 |
| **Hank’s** | | | | | | |
| *chd7^+/+^* – *chd7^ncu101/+^* | 18, 33 | 0.442481 | 0.219014 | 0.005081 | 0.879882 | 0.0475 |
| *chd7^+/+^* – *chd7^ncu101/ncu101^* | 18, 17 | 0.487082 | 0.252786 | -0.01777 | 0.99193 | 0.0584 |
| *chd7^ncu101/+^* – *chd7^ncu101/ncu101^* | 33, 17 | **0.044601** | 0.223143 | -0.40105 | 0.490248 | 0.8422 |
| **Normalized LLC Index (by genotype)** | | | | | | |
| **Comparison** | **(n)** | **p-Value** | **Mean Dif** | **Std Error Dif** | **Lower CL** | **Upper CL** |
| ***chd7^+/+^*** | | | | | | |
| 1x E3 – 1x E3 MB | 18, 18 | **0.0175** | 0.610803 | 0.251841 | 0.109811 | 1.111796 |
| 1x E3 – 1x E2 MB | 18, 13 | **0.011** | 0.715283 | 0.274993 | 0.168235 | 1.262331 |
| 1x E3 – 1x E2 | 18, 20 | 0.9965 | 0.001075 | 0.245465 | -0.48723 | 0.489383 |
| 1x E3 – Hank’s | 18, 18 | **0.0034** | 0.75893 | 0.251841 | 0.257937 | 1.259923 |
| 1x E2 – 1x E3 MB | 20, 18 | **0.015** | 0.609728 | 0.245465 | 0.121421 | 1.098035 |
| 1x E2 – 1x E2 MB | 20, 13 | **0.0096** | 0.714208 | 0.269165 | 0.178753 | 1.249663 |
| 1x E2 – Hank’s | 20, 18 | **0.0028** | 0.757855 | 0.245465 | 0.269547 | 1.246162 |
| 1x E3 MB – 1x E2 MB | 18, 13 | 0.705 | 0.10448 | 0.274993 | -0.44257 | 0.651528 |
| 1x E3 MB – Hank’s | 18, 18 | 0.558 | 0.148127 | 0.251841 | -0.35287 | 0.649119 |
| 1x E2 MB – Hank’s | 13, 18 | 0.8743 | 0.043647 | 0.274993 | -0.5034 | 0.590695 |
| ***chd7^ncu101/+^*** | | | | | | |
| 1x E3 – 1x E3 MB | 25, 38 | **0.0047** | 0.565564 | 0.197473 | 0.175575 | 0.955554 |
| 1x E3 – 1x E2 MB | 25, 33 | **0.0069** | 0.556822 | 0.203323 | 0.155279 | 0.958364 |
| 1x E3 – 1x E2 | 25, 36 | 0.1841 | 0.266345 | 0.199638 | -0.12792 | 0.660609 |
| 1x E3 – Hank’s | 25, 33 | 0.2119 | 0.254847 | 0.203323 | -0.1467 | 0.65639 |
| 1x E2 – 1x E3 MB | 36, 38 | 0.0954 | 0.29922 | 0.178349 | -0.053 | 0.651442 |
| 1x E2 – 1x E2 MB | 36, 33 | 0.118 | 0.290477 | 0.184806 | -0.0745 | 0.65545 |
| 1x E2 – Hank’s | 36, 33 | 0.9505 | 0.011497 | 0.184806 | -0.35348 | 0.37647 |
| 1x E3 MB – 1x E2 MB | 38, 33 | 0.9618 | 0.008743 | 0.182465 | -0.35161 | 0.369093 |
| 1x E3 MB – Hank’s | 38, 33 | 0.0905 | 0.310717 | 0.182465 | -0.04963 | 0.671067 |
| 1x E2 MB – Hank’s | 33, 33 | 0.1117 | 0.301974 | 0.18878 | -0.07085 | 0.674797 |
| ***chd7^ncu101/ncu101^*** | | | | | | |
| 1x E3 – 1x E3 MB | 15, 14 | 0.4736 | 0.190296 | 0.264125 | -0.33649 | 0.717076 |
| 1x E3 – 1x E2 MB | 15, 18 | 0.9833 | 0.005234 | 0.248482 | -0.49035 | 0.500816 |
| 1x E3 – 1x E2 | 15, 11 | 0.4776 | 0.201428 | 0.28214 | -0.36128 | 0.764138 |
| 1x E3 – Hank’s | 15, 17 | 0.0504 | 0.501199 | 0.251782 | -0.00097 | 1.003363 |
| 1x E2 – 1x E3 MB | 11, 14 | 0.1757 | 0.391723 | 0.286372 | -0.17943 | 0.962873 |
| 1x E2 – 1x E2 MB | 11, 18 | 0.4731 | 0.196193 | 0.272011 | -0.34632 | 0.738702 |
| 1x E2 – Hank’s | 11, 17 | **0.0128** | 0.702626 | 0.275029 | 0.154098 | 1.251154 |
| 1x E3 MB – 1x E2 MB | 14, 18 | 0.4427 | 0.19553 | 0.253277 | -0.30961 | 0.700674 |
| 1x E3 MB – Hank’s | 14, 17 | 0.2296 | 0.310903 | 0.256515 | -0.2007 | 0.822506 |
| 1x E2 MB – Hank’s | 18, 17 | **0.0387** | 0.506433 | 0.240377 | 0.027016 | 0.98585 |
| **Normalized Total Distance (by media)** | | | | | | |
| **Comparison** | **(n)** | **p-Value** | **Mean Dif** | **Std Error Dif** | **Lower CL** | **Upper CL** |
| **1x E3** | | | | | | |
| *chd7^+/+^* – *chd7^ncu101/+^* | 18, 26 | 0.8604 | 0.077804 | 0.440424 | -0.8038 | 0.959409 |
| *chd7^+/+^* – *chd7^ncu101/ncu101^* | 18, 17 | **0.0341** | 1.054542 | 0.485782 | 0.082144 | 2.026939 |
| *chd7^ncu101/+^* – *chd7^ncu101/ncu101^* | 26, 17 | **0.0142** | 1.132346 | 0.448013 | 0.23555 | 2.029142 |
| **1x E3 MB** | | | | | | |
| *chd7^+/+^* – *chd7^ncu101/+^* | 18, 38 | 0.1562 | 0.741691 | 0.517177 | -0.2906 | 1.77398 |
| *chd7^+/+^* – *chd7^ncu101/ncu101^* | 18, 14 | 0.5891 | 0.349571 | 0.644092 | -0.93604 | 1.635183 |
| *chd7^ncu101/+^* – *chd7^ncu101/ncu101^* | 38, 14 | 0.4901 | 0.392121 | 0.565091 | -0.73581 | 1.520048 |
| **1x E2** | | | | | | |
| *chd7^+/+^* – *chd7^ncu101/+^* | 21, 38 | 0.7581 | 0.099976 | 0.323235 | -0.5452 | 0.745155 |
| *chd7^+/+^* – *chd7^ncu101/ncu101^* | 21, 11 | 0.5294 | 0.279752 | 0.442448 | -0.60338 | 1.162882 |
| *chd7^ncu101/+^* – *chd7^ncu101/ncu101^* | 38, 11 | 0.3542 | 0.379728 | 0.407008 | -0.43266 | 1.19212 |
| **1x E2 MB** | | | | | | |
| *chd7^+/+^* – *chd7^ncu101/+^* | 13, 36 | 0.138 | 1.06754 | 0.711026 | -0.35207 | 2.487149 |
| *chd7^+/+^* – *chd7^ncu101/ncu101^* | 13, 20 | 0.3914 | 0.675402 | 0.782854 | -0.88762 | 2.23842 |
| *chd7^ncu101/+^* – *chd7^ncu101/ncu101^* | 36, 20 | 0.5245 | 0.392137 | 0.612827 | -0.83141 | 1.615687 |
| **Hank’s** | | | | | | |
| *chd7^+/+^* – *chd7^ncu101/+^* | 18, 33 | 0.6695 | 0.216849 | 0.505781 | -0.79298 | 1.226674 |
| *chd7^+/+^* – *chd7^ncu101/ncu101^* | 18, 18 | 0.0729 | 1.048604 | 0.575373 | -0.10017 | 2.197373 |
| *chd7^ncu101/+^* – *chd7^ncu101/ncu101^* | 33, 18 | 0.1048 | 0.831755 | 0.505781 | -0.17807 | 1.841579 |
| **Normalized Total Distance (by genotype)** | | | | | | |
| **Comparison** | **(n)** | **p-Value** | **Mean Dif** | **Std Error Dif** | **Lower CL** | **Upper CL** |
| ***chd7^+/+^*** | | | | | | |
| 1x E3 – 1x E3 MB | 18, 18 | 0.2871 | 0.563367 | 0.525822 | -0.48247 | 1.609205 |
| 1x E3 – 1x E2 MB | 18, 13 | **0.0001** | 2.309603 | 0.57416 | 1.16762 | 3.451585 |
| 1x E3 – 1x E2 | 18, 21 | 0.4326 | 0.39956 | 0.506695 | -0.60824 | 1.407355 |
| 1x E3 – Hank’s | 18, 18 | 0.0802 | 0.931303 | 0.525822 | -0.11454 | 1.977142 |
| 1x E2 – 1x E3 MB | 21, 18 | 0.7473 | 0.163807 | 0.506695 | -0.84399 | 1.171602 |
| 1x E2 – 1x E2 MB | 21, 13 | **0.0009** | 1.910044 | 0.556696 | 0.8028 | 3.01729 |
| 1x E2 – Hank’s | 21, 18 | 0.297 | 0.531744 | 0.506695 | -0.47605 | 1.539539 |
| 1x E3 MB – 1x E2 MB | 18, 13 | **0.0032** | 1.746237 | 0.57416 | 0.60426 | 2.888218 |
| 1x E3 MB – Hank’s | 18, 18 | 0.486 | 0.367937 | 0.525822 | -0.6779 | 1.413776 |
| 1x E2 MB – Hank’s | 13, 18 | **0.0186** | 1.3783 | 0.57416 | 0.23632 | 2.520281 |
| ***chd7^ncu101/+^*** | | | | | | |
| 1x E3 – 1x E3 MB | 26, 38 | **0.0004** | 1.382862 | 0.383587 | 0.625523 | 2.1402 |
| 1x E3 – 1x E2 MB | 26, 36 | **0.0008** | 1.319868 | 0.387892 | 0.554031 | 2.085705 |
| 1x E3 – 1x E2 | 26, 38 | 0.1342 | 0.57734 | 0.383587 | -0.18 | 1.334679 |
| 1x E3 – Hank’s | 26, 33 | **0.0023** | 1.225957 | 0.395217 | 0.445658 | 2.006255 |
| 1x E2 – 1x E3 MB | 38, 38 | **0.021** | 0.805522 | 0.345761 | 0.122866 | 1.488177 |
| 1x E2 – 1x E2 MB | 38, 36 | **0.0356** | 0.742528 | 0.35053 | 0.050456 | 1.4346 |
| 1x E2 – Hank’s | 38, 33 | 0.0723 | 0.648616 | 0.358619 | -0.05943 | 1.356658 |
| 1x E3 MB – 1x E2 MB | 38, 36 | 0.8576 | 0.062994 | 0.35053 | -0.62908 | 0.755066 |
| 1x E3 MB – Hank’s | 38, 33 | 0.6623 | 0.156905 | 0.358619 | -0.55114 | 0.864947 |
| 1x E2 MB – Hank’s | 36, 33 | 0.7963 | 0.093911 | 0.363219 | -0.62321 | 0.811036 |
| ***chd7^ncu101/ncu101^*** | | | | | | |
| 1x E3 – 1x E3 MB | 17, 14 | 0.8593 | 0.141605 | 0.796137 | -1.44438 | 1.72759 |
| 1x E3 – 1x E2 MB | 17, 20 | 0.4282 | 0.579659 | 0.727707 | -0.87001 | 2.029326 |
| 1x E3 – 1x E2 | 17, 11 | 0.277 | 0.934734 | 0.853598 | -0.76572 | 2.635189 |
| 1x E3 – Hank’s | 17, 18 | 0.2187 | 0.925365 | 0.746051 | -0.56084 | 2.411575 |
| 1x E2 – 1x E3 MB | 11, 14 | 0.3751 | 0.793129 | 0.888802 | -0.97745 | 2.563713 |
| 1x E2 – 1x E2 MB | 11, 20 | **0.0714** | 1.514393 | 0.828066 | -0.1352 | 3.163984 |
| 1x E2 – Hank’s | 11, 18 | **0.0306** | 1.860099 | 0.844232 | 0.1783 | 3.541895 |
| 1x E3 MB – 1x E2 MB | 14, 20 | 0.3511 | 0.721264 | 0.768698 | -0.81006 | 2.252589 |
| 1x E3 MB – Hank’s | 14, 18 | 0.1788 | 1.06697 | 0.786086 | -0.49899 | 2.632933 |
| 1x E2 MB – Hank’s | 20, 18 | 0.631 | 0.345706 | 0.716697 | -1.08203 | 1.77344 |
| **Swim Frequency (by media)** | | | | | | |
| **Comparison** | **(n)** | **p-Value** | **Mean Dif** | **Std Error Dif** | **Lower CL** | **Upper CL** |
| **1x E3** | | | | | | |
| *chd7^+/+^* – *chd7^ncu101/+^* | 18, 26 | 0.6184 | 1.235043 | 2.466323 | -3.70184 | 6.171924 |
| *chd7^+/+^* – *chd7^ncu101/ncu101^* | 18, 17 | 0.989 | 0.037582 | 2.720319 | -5.40773 | 5.482892 |
| *chd7^ncu101/+^* – *chd7^ncu101/ncu101^* | 26, 17 | 0.6139 | 1.272624 | 2.508821 | -3.74933 | 6.294575 |
| **1x E3 MB** | | | | | | |
| *chd7^+/+^* – *chd7^ncu101/+^* | 18, 38 | 0.1812 | 1.847368 | 1.366933 | -0.8818 | 4.576538 |
| *chd7^+/+^* – *chd7^ncu101/ncu101^* | 18, 14 | **0.0105** | 4.580769 | 1.738818 | 1.10911 | 8.052432 |
| *chd7^ncu101/+^* – *chd7^ncu101/ncu101^* | 38, 14 | 0.0796 | 2.733401 | 1.53498 | -0.33129 | 5.798088 |
| **1x E2** | | | | | | |
| *chd7^+/+^* – *chd7^ncu101/+^* | 21, 38 | 0.1315 | 3.245739 | 2.125951 | -0.99768 | 7.489155 |
| *chd7^+/+^* – *chd7^ncu101/ncu101^* | 21, 11 | 0.5042 | 1.954113 | 2.910031 | -3.85433 | 7.762559 |
| *chd7^ncu101/+^* – *chd7^ncu101/ncu101^* | 38, 11 | 0.631 | 1.291627 | 2.676939 | -4.05157 | 6.63482 |
| **1x E2 MB** | | | | | | |
| *chd7^+/+^* – *chd7^ncu101/+^* | 13, 36 | 0.3049 | 1.954525 | 1.889697 | -1.82058 | 5.729629 |
| *chd7^+/+^* – *chd7^ncu101/ncu101^* | 13, 20 | 0.6911 | 0.824231 | 2.064547 | -3.30018 | 4.948638 |
| *chd7^ncu101/+^* – *chd7^ncu101/ncu101^* | 36, 20 | 0.4914 | 1.130294 | 1.63304 | -2.13208 | 4.392667 |
| **Hank’s** | | | | | | |
| *chd7^+/+^* – *chd7^ncu101/+^* | 18, 33 | 0.2111 | 2.44798 | 1.938642 | -1.42264 | 6.318604 |
| *chd7^+/+^* – *chd7^ncu101/ncu101^* | 18, 18 | 0.2916 | 2.344444 | 2.205386 | -2.05875 | 6.747639 |
| *chd7^ncu101/+^* – *chd7^ncu101/ncu101^* | 33, 18 | 0.9576 | 0.103535 | 1.938642 | -3.76709 | 3.974159 |
| **Swim Frequency (by genotype)** | | | | | | |
| **Comparison** | **(n)** | **p-Value** | **Mean Dif** | **Std Error Dif** | **Lower CL** | **Upper CL** |
| ***chd7^+/+^*** | | | | | | |
| 1x E3 – 1x E3 MB | 18, 18 | 0.1615 | 3.011111 | 2.13166 | -1.22868 | 7.250898 |
| 1x E3 – 1x E2 MB | 18, 13 | 0.9762 | 0.069658 | 2.32762 | -4.55989 | 4.699202 |
| 1x E3 – 1x E2 | 18, 21 | 0.6444 | 0.951587 | 2.054119 | -3.13397 | 5.037148 |
| 1x E3 – Hank’s | 18, 18 | 0.8761 | 0.333333 | 2.13166 | -3.90645 | 4.57312 |
| 1x E2 – 1x E3 MB | 21, 18 | 0.319 | 2.059524 | 2.054119 | -2.02604 | 6.145084 |
| 1x E2 – 1x E2 MB | 21, 13 | 0.6521 | 1.021245 | 2.256822 | -3.46748 | 5.509974 |
| 1x E2 – Hank’s | 21, 18 | 0.7642 | 0.618254 | 2.054119 | -3.46731 | 4.703814 |
| 1x E3 MB – 1x E2 MB | 18, 13 | 0.1893 | 3.080769 | 2.32762 | -1.54877 | 7.710313 |
| 1x E3 MB – Hank’s | 18, 18 | 0.2126 | 2.677778 | 2.13166 | -1.56201 | 6.917564 |
| 1x E2 MB – Hank’s | 13, 18 | 0.863 | 0.402991 | 2.32762 | -4.22655 | 5.032535 |
| ***chd7^ncu101/+^*** | | | | | | |
| 1x E3 – 1x E3 MB | 26, 38 | 0.1758 | 2.398785 | 1.764141 | -1.08457 | 5.882143 |
| 1x E3 – 1x E2 MB | 26, 36 | 0.6627 | 0.78914 | 1.805807 | -2.77649 | 4.354768 |
| 1x E3 – 1x E2 | 26, 38 | 0.5491 | 1.059109 | 1.764141 | -2.42425 | 4.542467 |
| 1x E3 – Hank’s | 26, 33 | 0.6291 | 0.879604 | 1.817624 | -2.70936 | 4.468566 |
| 1x E2 – 1x E3 MB | 38, 38 | 0.0311 | 3.457895 | 1.590175 | 0.31804 | 6.597751 |
| 1x E2 – 1x E2 MB | 38, 36 | 0.8692 | 0.269969 | 1.636277 | -2.96092 | 3.500854 |
| 1x E2 – Hank’s | 38, 33 | 0.9135 | 0.179506 | 1.64931 | -3.07711 | 3.436125 |
| 1x E3 MB – 1x E2 MB | 38, 36 | 0.0531 | 3.187926 | 1.636277 | -0.04296 | 6.418811 |
| 1x E3 MB – Hank’s | 38, 33 | **0.0485** | 3.278389 | 1.64931 | 0.02177 | 6.535008 |
| 1x E2 MB – Hank’s | 36, 33 | 0.9575 | 0.090463 | 1.693803 | -3.25401 | 3.434935 |
| ***chd7^ncu101/ncu101^*** | | | | | | |
| 1x E3 – 1x E3 MB | 17, 14 | 0.5037 | 1.60724 | 2.392046 | -3.15902 | 6.373495 |
| 1x E3 – 1x E2 MB | 17, 20 | 0.6649 | 0.931471 | 2.141738 | -3.33604 | 5.198977 |
| 1x E3 – 1x E2 | 17, 11 | 0.6801 | 1.040107 | 2.512252 | -3.96566 | 6.045879 |
| 1x E3 – Hank’s | 17, 18 | 0.3538 | 2.048693 | 2.195727 | -2.32639 | 6.423774 |
| 1x E2 – 1x E3 MB | 11, 14 | 0.8317 | 0.567133 | 2.659761 | -4.73256 | 5.866822 |
| 1x E2 – 1x E2 MB | 11, 20 | 0.9646 | 0.108636 | 2.437107 | -4.7474 | 4.964677 |
| 1x E2 – Hank’s | 11, 18 | 0.686 | 1.008586 | 2.484685 | -3.94226 | 5.95943 |
| 1x E3 MB – 1x E2 MB | 14, 20 | 0.771 | 0.675769 | 2.312998 | -3.93298 | 5.284519 |
| 1x E3 MB – Hank’s | 14, 18 | 0.8523 | 0.441453 | 2.363077 | -4.26708 | 5.149987 |
| 1x E2 MB – Hank’s | 20, 18 | 0.5979 | 1.117222 | 2.109335 | -3.08572 | 5.320164 |
| **Turn Frequency (by media)** | | | | | | |
| **Comparison** | **(n)** | **p-Value** | **Mean Dif** | **Std Error Dif** | **Lower CL** | **Upper CL** |
| **1x E3** | | | | | | |
| *chd7^+/+^* – *chd7^ncu101/+^* | 18, 26 | 0.5115 | 3.535897 | 5.352362 | -7.178 | 14.24981 |
| *chd7^+/+^* – *chd7^ncu101/ncu101^* | 18, 17 | 0.9811 | 0.140196 | 5.903579 | -11.6771 | 11.95749 |
| *chd7^ncu101/+^* – *chd7^ncu101/ncu101^* | 26, 17 | 0.5353 | 3.395701 | 5.44459 | -7.5028 | 14.29423 |
| **1x E3 MB** | | | | | | |
| *chd7^+/+^* – *chd7^ncu101/+^* | 18, 38 | 0.268 | 5.79737 | 5.189855 | -4.56451 | 16.15925 |
| *chd7^+/+^* – *chd7^ncu101/ncu101^* | 18, 14 | 0.3449 | 6.28077 | 6.601797 | -6.90014 | 19.46168 |
| *chd7^ncu101/+^* – *chd7^ncu101/ncu101^* | 38, 14 | **0.0421** | 12.07814 | 5.827883 | 0.44239 | 23.71388 |
| **1x E2** | | | | | | |
| *chd7^+/+^* – *chd7^ncu101/+^* | 21, 38 | 0.2099 | 6.34699 | 5.013997 | -3.66099 | 16.35497 |
| *chd7^+/+^* – *chd7^ncu101/ncu101^* | 21, 11 | 0.3248 | 6.80779 | 6.86323 | -6.89127 | 20.50686 |
| *chd7^ncu101/+^* – *chd7^ncu101/ncu101^* | 38, 11 | **0.041** | 13.15478 | 6.313489 | 0.55301 | 25.75656 |
| **1x E2 MB** | | | | | | |
| *chd7^+/+^* – *chd7^ncu101/+^* | 13, 36 | 0.384 | 5.0905 | 5.806713 | -6.50975 | 16.69074 |
| *chd7^+/+^* – *chd7^ncu101/ncu101^* | 13, 20 | **0.033** | 13.82462 | 6.343996 | 1.15103 | 26.4982 |
| *chd7^ncu101/+^* – *chd7^ncu101/ncu101^* | 36, 20 | 0.0866 | 8.73412 | 5.01805 | -1.29059 | 18.75883 |
| **Hank’s** | | | | | | |
| *chd7^+/+^* – *chd7^ncu101/+^* | 18, 33 | 0.7942 | 1.394949 | 5.32582 | -9.23839 | 12.02829 |
| *chd7^+/+^* – *chd7^ncu101/ncu101^* | 18, 18 | 0.5872 | 3.305556 | 6.058615 | -8.79086 | 15.40197 |
| *chd7^ncu101/+^* – *chd7^ncu101/ncu101^* | 33, 18 | 0.3807 | 4.700505 | 5.32582 | -5.93284 | 15.33385 |
| **Turn Frequency (by genotype)** | | | | | | |
| **Comparison** | **(n)** | **p-Value** | **Mean Dif** | **Std Error Dif** | **Lower CL** | **Upper CL** |
| ***chd7^+/+^*** | | | | | | |
| 1x E3 – 1x E3 MB | 18, 18 | 0.431 | 4.93333 | 6.234543 | -7.46692 | 17.33359 |
| 1x E3 – 1x E2 MB | 18, 13 | **0.0058** | 19.26795 | 6.807675 | 5.72776 | 32.80814 |
| 1x E3 – 1x E2 | 18, 21 | 0.6718 | 2.55476 | 6.007756 | -9.39442 | 14.50395 |
| 1x E3 – Hank’s | 18, 18 | 0.0704 | 11.42778 | 6.234543 | -0.97248 | 23.82803 |
| 1x E2 – 1x E3 MB | 21, 18 | 0.6932 | 2.37857 | 6.007756 | -9.57061 | 14.32776 |
| 1x E2 – 1x E2 MB | 21, 13 | **0.0132** | 16.71319 | 6.600609 | 3.58484 | 29.84153 |
| 1x E2 – Hank’s | 21, 18 | 0.1435 | 8.87302 | 6.007756 | -3.07617 | 20.8222 |
| 1x E3 MB – 1x E2 MB | 18, 13 | **0.0383** | 14.33462 | 6.807675 | 0.79442 | 27.87481 |
| 1x E3 MB – Hank’s | 18, 18 | 0.3006 | 6.49444 | 6.234543 | -5.90581 | 18.8947 |
| 1x E2 MB – Hank’s | 13, 18 | 0.2528 | 7.84017 | 6.807675 | -5.70002 | 21.38036 |
| ***chd7^ncu101/+^*** | | | | | | |
| 1x E3 – 1x E3 MB | 26, 38 | **0.0016** | 14.2666 | 4.444851 | 5.49009 | 23.04311 |
| 1x E3 – 1x E2 MB | 26, 36 | **0.0001** | 17.71335 | 4.54983 | 8.72955 | 26.69715 |
| 1x E3 – 1x E2 | 26, 38 | **0.0058** | 12.43765 | 4.444851 | 3.66114 | 21.21416 |
| 1x E3 – Hank’s | 26, 33 | **0.0005** | 16.35862 | 4.579606 | 7.31604 | 25.40121 |
| 1x E2 – 1x E3 MB | 38, 38 | 0.6486 | 1.82895 | 4.006535 | -6.08209 | 9.73999 |
| 1x E2 – 1x E2 MB | 38, 36 | 0.2025 | 5.2757 | 4.12269 | -2.8647 | 13.41609 |
| 1x E2 – Hank’s | 38, 33 | 0.3468 | 3.92097 | 4.155527 | -4.28426 | 12.1262 |
| 1x E3 MB – 1x E2 MB | 38, 36 | 0.4043 | 3.44675 | 4.12269 | -4.69365 | 11.58714 |
| 1x E3 MB – Hank’s | 38, 33 | 0.6153 | 2.09203 | 4.155527 | -6.11321 | 10.29726 |
| 1x E2 MB – Hank’s | 36, 33 | 0.7513 | 1.35472 | 4.267629 | -7.07186 | 9.78131 |
| ***chd7^ncu101/ncu101^*** | | | | | | |
| 1x E3 – 1x E3 MB | 17, 14 | 0.8596 | 1.20724 | 6.800608 | -12.3433 | 14.75775 |
| 1x E3 – 1x E2 MB | 17, 20 | 0.3621 | 5.58353 | 6.088982 | -6.549 | 17.71609 |
| 1x E3 – 1x E2 | 17, 11 | 0.5665 | 4.11283 | 7.142356 | -10.1186 | 18.34429 |
| 1x E3 – Hank’s | 17, 18 | 0.1897 | 8.26242 | 6.242472 | -4.176 | 20.70081 |
| 1x E2 – 1x E3 MB | 11, 14 | 0.7019 | 2.90559 | 7.561725 | -12.1615 | 17.97266 |
| 1x E2 – 1x E2 MB | 11, 20 | 0.1659 | 9.69636 | 6.928716 | -4.1094 | 23.50213 |
| 1x E2 – Hank’s | 11, 18 | 0.0839 | 12.37525 | 7.063983 | -1.7 | 26.45055 |
| 1x E3 MB – 1x E2 MB | 14, 20 | 0.3051 | 6.79077 | 6.575875 | -6.3119 | 19.89349 |
| 1x E3 MB – Hank’s | 14, 18 | 0.1629 | 9.46966 | 6.71825 | -3.9167 | 22.85606 |
| 1x E2 MB – Hank’s | 20, 18 | 0.6564 | 2.67889 | 5.996859 | -9.2701 | 14.62789 |

**Supplemental Table 4.** **Multiple comparisons of *chd7* mutants within and between media types.** Multiple comparisons of *chd7^+/+^, chd7^ncu101/+^,* and *chd7^ncu101/ncu101^*  in 1x E3, 1x E3 MB, 1x E2, 1x E2 MB, of Hank’s and their reported values for the acoustic startle response (normalized SLC and LLC Index), and general locomotor behaviors (total distance, swim and turn frequency) with statistically significant p-values in bold (α=0.05, ANOVA with student’s t each pair test for multiple comparisons).
